# X-CODE: a dual RNA barcoding system for multi-platform clonal tracking and spatial phenotyping

**DOI:** 10.64898/2026.02.10.705126

**Authors:** Roberto Rota, Elizabeth Flittner, Matteo Tartagni, Marwane Bourdim, Emma Westlund, James S. McKenzie, Anthony Devlin, Archana Thankamony, Nicole Pandell, Florian Gabel, Misha Siddiqui, Dalia Rosano, Alexander Wilkes, Ivan Andrew, Ka Lok Choi, Laurence Game, Chiara Gorrini, Luca Magnani, Zoltan Takats, Marco Bezzi

**Author notes:** Equal contribution (Roberto Rota, Elizabeth Flittner).

## Abstract

Experimental dissection of clonal dynamics in complex tissues requires barcoding systems that are scalable, compatible with different analytical platforms, providing phenotypic and spatial resolution. Here we introduce X-CODE, a dual-expressed RNA barcoding system designed to enable high-complexity clonal tracking across sequencing-based, cytometric, and spatial imaging modalities within a unified experimental framework. X-CODE combines a combinatorial, probe-detectable long RNA barcode with a matched short sequencing barcode, enabling seamless integration of probe-based readouts with sequencing and barcode-guided clonal retrieval. We demonstrate robust X-CODE detection by mass cytometry and imaging-based platforms, including spatial RNA barcode readout using via a repurposed Akoya PhenoCycler-Fusion protocol. In addition, we show compatibility with MALDI mass spectrometry imaging for co-registration of clonal and metabolic information. We further demonstrate the feasibility of X-CODE detection within probe-based spatial transcriptomics using the 10x Genomics Xenium platform. Applied to an *in vivo* model of androgen deprivation in prostate cancer, X-CODE reveals clonal architecture, selection and clone-specific phenotypic and metabolic plasticity underlying castration resistance. Together, X-CODE provides a flexible and broadly accessible platform for integrated clonal analysis across spatial, phenotypic, and molecular dimensions.

## INTRODUCTION

Understanding how complex tissues arise, maintain themselves, and adapt over time has long relied on the ability to follow the descendants of individual cells. Lineage tracing approaches, first established in developmental systems, provided a framework to connect cell of origin with fate, position, and function^1^. These principles have increasingly been adopted in cancer research, where tumours are now recognised as dynamic, evolving populations shaped by continuous selection^2,3^.

Lineage relationships in cancer can be reconstructed retrospectively from patient samples by analysing endogenous somatic features, including genetic, mitochondrial, or epigenetic variation^4–11^. Such approaches have delivered important insights into tumour evolution and clonal architecture and will continue to advance with improvements in sequencing technologies and computational inference^12^. However, retrospective strategies are inherently limited in their ability to functionalise defined genetic or therapeutic perturbations or to directly interrogate how interactions with the tumour microenvironment shape clonal behaviour. By contrast, prospective lineage tracing introduces heritable labels into cells prior to perturbation, enabling direct observation of how individual clones develop and respond to selective pressures such as drug treatment, environmental stress, or microenvironmental change in controlled preclinical systems^13^.

Molecular barcoding has emerged as the dominant strategy for prospective lineage tracing, allowing large numbers of clones to be tracked simultaneously within heterogeneous populations^14–16^. When combined with single-cell RNA sequencing (scRNA-seq), prospective barcoding enables clonal fitness to be linked to molecular state^17–20^, providing a powerful framework for studying tumour adaptation and resistance. Such studies have revealed pronounced clonal asymmetries during tumour progression and therapy, often identifying rare populations that disproportionately contribute to long-term survival.

Current barcoding technologies broadly fall into two classes: evolvable and static systems. Evolvable barcodes incorporate arrays of mutable sites that accumulate edits across successive cell divisions, enabling reconstruction of lineage relationships base d on shared mutation patterns (e.g. ZOMBIE^21^, KPtracer^22^, GESTALT^23^, CARLIN^24^, macsGESTALT^25^). Advances using base or prime editors have improved the predictability of these edits (e.g. baseMEMOIR^26^ ; PEtracer^27^, DNA Typewriter^28^). Nevertheless, evolvable systems face inherent limitations related to recording capacity, saturation over time, dependence on sustained genome editing activity, and the need for system optimisation.

Static barcode libraries provide stable, high-complexity identifiers that simplify experimental implementation and debarcoding but do not encode intra-generational lineage relationships. These systems have been implemented at the DNA, RNA, and protein levels, each with distinct advantages and limitations. DNA barcodes (e.g. ClonTracer^29^), among the earliest introduced, consist of variable or semi-variable DNA sequences delivered via viral vectors and rely exclusively on next-generation sequencing (NGS) for readout, limiting direct phenotypic or spatial characterisation. Transcribed RNA barcodes retain high complexity while enabling integration with scRNA-seq (e.g. LARRY^30^, ClonMapper^31^, CloneTracker^TM^). Notably, recent systems such as CaTCH^32^ and ClonMapper^31^, support barcode-specific clonal retrieval via VPR–Cas9-mediated reporter activation. However, RNA barcode platforms remain constrained by the cost, scalability, and limited spatial compatibility of sequencing-based readouts. Protein barcodes based on combinations of fluorescent proteins^33^ or combinatorial epitope arrays^34^ enable direct phenotypic and spatial readout using antibody-based platforms such as flow cytometry, fluorescence microscopy, mass cytometry, or multiplexed ion beam imaging. However, the limited number of orthogonal epitopes and the complexity of engineering large combinatorial constructs restrict achievable library size and, consequently, clonal resolution in large-scale studies.

Together, these limitations highlight a central bottleneck in current lineage-tracing technologies: the absence of a high-complexity barcode system that is agnostic to the readout platform while remaining compatible with scalable, cost-effective phenotypic and spatial assays, thereby enabling broad applicability across model systems and the wider scientific community.

Probe-based RNA detection offers an attractive route to address this gap. RNA in situ hybridisation approaches that incorporate signal amplification strategies, such as rolling circle amplification, enable sensitive and specific detection using relatively simple probe architectures^35–42^. Crucially, probe-based detection decouples barcode identity from detection modality, allowing the same probes to be paired with fluorophores or metal isotopes depending on the application. This creates the opportunity for a single barcode design to be read across orthogonal platforms, including flow cytometry, mass cytometry, and spatial imaging. However, the practical utility of probe-based strategies is constrained by probe number and structural complexity, which can substantially increase costs, lengthen experimental workflows, and limit compatibility with certain platforms. Achieving an effective balance between multiplexing capacity, detection sensitivity, and practical accessibility, therefore represents a central design challenge for probe-based barcode systems.

Here, we introduce X-CODE, a universal RNA barcoding system coupled to a dedicated probe-based detection strategy that enables robust, multi-platform barcode readout at single-cell and spatial resolution. X-CODE supports high-complexity static barcoding, integrates seamlessly with sequencing-based workflows, and is directly compatible with high-throughput phenotypic technologies, including CyTOF, multiplex immunofluorescence, imaging mass cytometry, the Akoya PhenoCycler-Fusion, and 10x Genomics Xenium. In addition, X-CODE enables barcode-specific clonal retrieval for downstream functional characterisation. We apply this framework to investigate the emergence of castration resistance in prostate cancer, a highly heterogeneous disease in which resistance to androgen-deprivation therapy arises through multiple evolutionary trajectories. By enabling clonal tracking within intact spatial architecture and across complementary analytical platforms, X-CODE provides a flexible and broadly accessible framework for interrogating development, tumour evolution, and therapeutic resistance.

## RESULTS

### Design and generation of a scalable, multiplatform-compatible barcode system

Achieving sufficient barcode diversity for clonal lineage tracing with probe-based RNA detection requires generating complexity through combinatorial design rather than large numbers of unique probes, given the practical limits on multiplexing imposed by probe chemistry and detection platforms. We therefore developed the X-CODE framework, in which unique barcode identities are encoded by combinations of modular Barcode Units (BUs) assembled from a limited set of short RNA sequence elements.

To enable probe-based detection of these BUs using a minimal probe set, we developed an RNA detection strategy integrating key principles from established probe-based RNA detection approaches and CRISPR screening readouts^41,43,44^. X-CODE employs a dual-probe architecture designed to ensure specificity and signal amplification (Fig. 1A). Each BU is targeted by two probes arranged in a “snail” configuration, comprising a padlock probe (“snail shell”) and a linear primer probe (“snail tail”) that must simultaneously bind the target RNA. Correct binding enables ligation of the padlock probe followed by rolling circle amplification (RCA), generating concatemeric DNA amplicons that are detected using fluorescently or metal-labelled probes. Each BU consists of a 20-bp variable binding sequence (VBS) targeted by a BU-specific padlock probe, followed by a 20-bp universal binding sequence (UBS) shared across all BUs and targeted by a single universal tail probe, enabling multiplexed detection with a limited probe repertoire.

**Figure 1.**
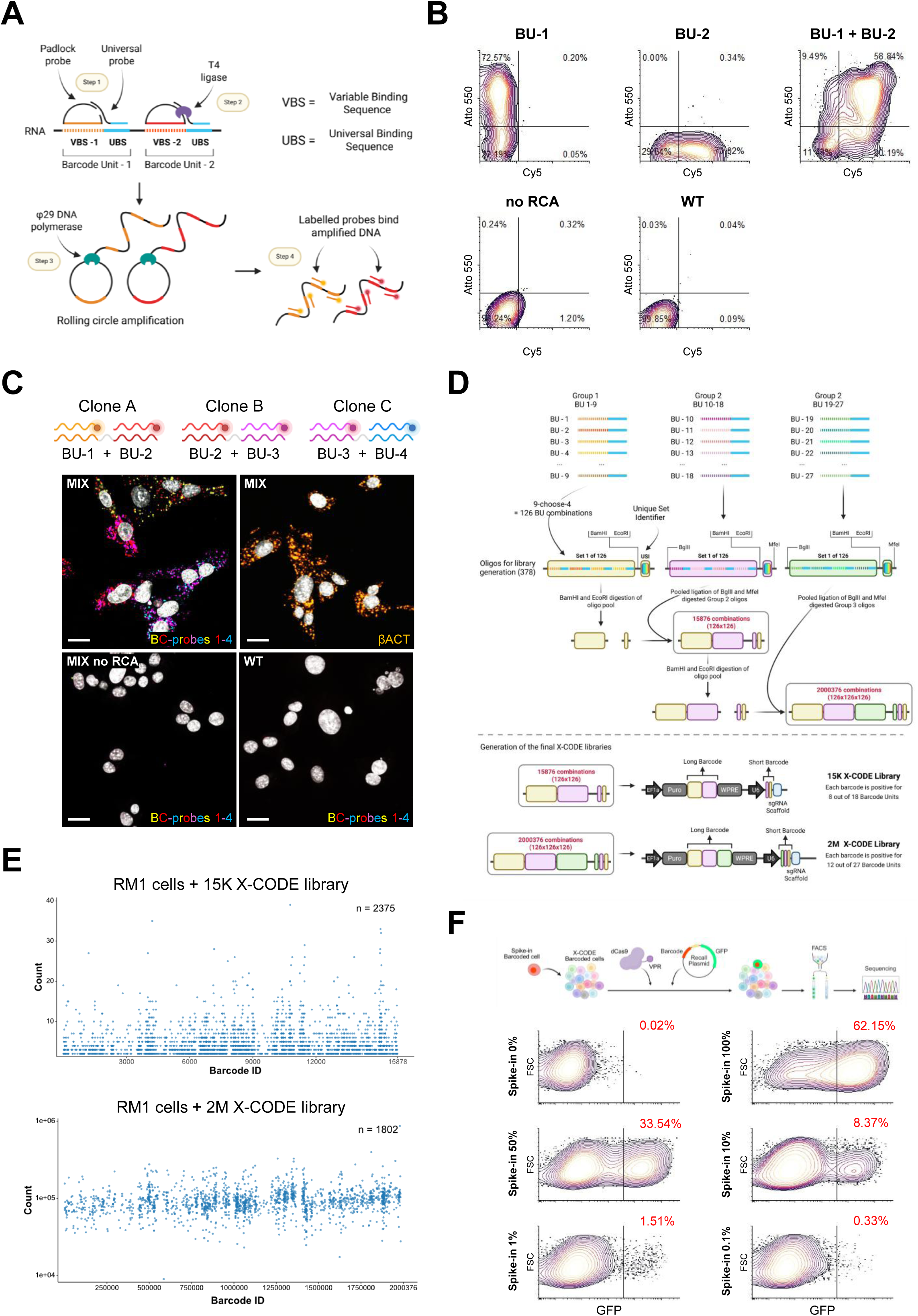
X-CODE concept and generation. **A**) Schematic of the X-CODE probe-based detection workflow (created with https://BioRender.com). Individual barcode units (BUs) are detected using complementary fluorescent probes following rolling circle amplification (RCA), enabling single- or multiple-barcode identification within individual cells. **B**) Flow cytometry contour plots of three barcoded TRAMP-C1 cell lines expressing BU-1, BU-2, or BU-1+BU-2. BU-1 and BU-2 were detected using Atto-550– and Cy5-conjugated probes, respectively. No RCA control was processed omitting the RCA step. WT sample are TRAMP-C1 cells without barcode. **C**) Fluorescence confocal imaging of three barcoded clones, each carrying a unique combination of two out of four barcode units (Clone A: BU-1+BU-2; Clone B: BU-2+BU-3; Clone C: BU-3+BU-4). Top left: equal mix sample containing all three clones processed with the X-CODE detection workflow. In yellow, BU-1; in red, BU-2; in magenta, BU-3; in cyan, BU-4. Top right: detection of β-actin using the X-CODE probe protocol. The mix no-RCA negative control demonstrates loss of signal in the absence of RCA, and WT cells lack barcodes. Fluorophores used for detection: BU-1–Alexa 488, BU-2–Atto 550, BU-3–Cy5, BU-4–AF750, β-actin–Alexa 488. Scale bars, 20 µM. **D**) Schematic representation of the combinatorial strategy developed for the assembly of the 15K and 2M barcode libraries (created with https://BioRender.com). **E**) NGS analysis of barcode representation in RM-1 cells transduced at low MOI with X-CODE libraries. Top: Scatter plot of NGS amplicon sequencing of short barcodes recovered from RM-1 cells transduced at low MOI with the 15K X-CODE library. Barcodes were PCR-amplified, gel-purified, and sequenced. The x axis shows individual barcode IDs, and the y-axis shows read counts. A total of 2,375 unique barcodes were detected; only barcodes with counts > 1 are plotted. Bottom: Equivalent analysis performed on RM-1 cells transduced at low MOI with the 2M X-CODE library. A total of 1,802 unique barcodes were identified; barcodes plotted are filtered for counts > 8,000. **F**) Clonal spike-in retrieval sensitivity assay using X-CODE recall plasmid activation. Top schematic: Experimental workflow (created with https://BioRender.com). A population of barcoded cells carrying a defined barcode was spiked into a bulk X-CODE–tagged cell population at defined ratios. Cells were transfected with dCas9-VPR and a barcode-specific recall plasmid, inducing GFP expression only in cells harbouring the cognate barcode. After 48 h, GFP⁺ cells were quantified by flow cytometry, sorted, and sequenced for validation. Bottom panels: Flow cytometry contour plots showing GFP activation across six spike-in dilution conditions (0%, 100%, 50%, 10%, 1%, 0.1%). The percentage refers to the fraction of spike-in cells in the initial mixture (red values). GFP⁺ enrichment scales with spike-in frequency, demonstrating barcode-specific recall sensitivity across >3 orders of magnitude, with minimal background at 0% and 0.1% spike-in control conditions.

To validate this detection strategy in suspension cells, we generated TRAMP-C1 cell lines expressing BU-1, BU-2, or the combined BU-1+BU-2 configuration. Flow cytometry demonstrated robust and specific detection of individual BUs, with dual-BU cells resolved by double-positive probe signal (Fig. 1B). By contrast, omission of the RCA step or analysis of wild-type cells lacking X-CODE constructs resulted in minimal background signal, confirming both RCA dependence and barcode specificity. We next assessed spatial detection of X-CODE barcodes by immunofluorescence imaging. Three cell lines carrying small-scale barcodes composed of unique combinations of two out of four BUs were mixed at equal proportions, seeded onto slides, and processed using the X-CODE imaging protocol. This analysis yielded high-quality signals for all four BUs and enabled discrimination of the three clones based on their unique BU combinations (Fig. 1C), demonstrating that combinatorial assembly of a limited pool of short, probe-detectable RNA elements supports spatially resolved barcode identification.

Following this validation, we developed a strategy for assembling high-complexity X-CODE libraries under three constraints: compatibility with probe-based detection platforms, scalable cloning without cost-prohibitive synthesis of thousands of unique long oligonucleotides, and incorporation of a compact secondary barcode compatible with sequencing-based readouts and barcode-specific clonal retrieval while remaining intrinsically linked to the probe-detectable barcode identity. To address these constraints, we selected 27 VBSs for BU construction, 23 were adapted from insert probes previously validated in the PLAYR system^43^, 4 were developed following the PLAYR criteria (Fig. 1D, top; Table 2 – 27_BUs). A single UBS was designed by combining two backbone-binding regions described in the same work and paired with each VBS to generate 27 barcode units, each 41 bp in length. All sequences were evaluated for GC content, predicted secondary structure, and annealing temperature. Library construction was achieved through a combinatorial, pooled ligation strategy^45^. The 27 BUs were divided into three groups of nine, and all combinations of four units within each group (C(9,4) = 126) were generated (Fig. 1D, top). Each combination was linked to a unique 4-bp unique set identifier (USI) contributing to the variable region of a secondary short barcode. These pools were sequentially combined through iterative cycles of pooled digestion and ligation (Fig. 1D, middle), expanding library complexity from 126 to 15,876 and ultimately to 2,000,376 unique barcodes, after which additional vector components were incorporated. In the final design, long and short barcodes are inherently matched, enabling direct cross - referencing across sequencing- and probe-based datasets. The short barcode is expressed downstream of a U6 promoter and upstream of a scaffold sequence, enabling barcode-specific clonal retrieval and providing a compact solution for DNA sequencing and scRNA-seq–based barcode recovery. Using this approach, we generated two libraries: a 15,876-member (15K) library and a 2,000,376-member (2M) library. Long barcodes are encoded by unique combinations of 8 out of 18 or 12 out of 27 BUs, respectively, while short barcodes consist of two or three 4-bp blocks separated by fixed 6-bp linker sequences (Fig. 1D, bottom).

Whole-plasmid sequencing confirmed correct library assembly and validated the expected pairing between long and short barcodes in both libraries (Supplementary Fig. 1A). Amplicon - based next-generation sequencing of short barcodes from cells transduced at low multiplicity of infection (MOI) further confirmed broad barcode representation and library balance (Fig. 1E).

Finally, to validate compatibility with barcode-specific clonal retrieval based on short barcodes of different lengths, we generated two clonal cell lines tagged with unique barcodes from either the 15K or 2M library. These clones were spiked at defined ratios into bulk populations tagged with the corresponding libraries and transfected with a barcode-specific recall reporter plasmid, designed as previously described^46^, together with dCas9–VPR. Forty-eight hours after transfection, cells were sorted based on EGFP expression and barcode identity confirmed by Sanger sequencing (Fig. 1F; Supplementary Fig. 1B,C). Barcode recall scaled with spike-in frequency across multiple orders of magnitude, while recovery efficiency decreased at higher spike-in fractions, consistent with observations reported for similar systems^31,32,47^.

Together, these results establish X-CODE as a modular, combinatorial RNA barcoding system that enables high barcode complexity using a limited set of probe-detectable elements, while maintaining compatibility with sequencing-based readouts and barcode-specific clonal retrieval.

### High-throughput X-CODE readout by mass cytometry

To determine whether X-CODE barcodes could be detected at high dimensionality while preserving simultaneous phenotypic profiling, we adapted the probe-based detection workflow for mass cytometry (CyTOF). CyTOF enables routine analysis of millions of single cells per day across many samples through sample multiplexing strategies such as TOBis^48^, while its >40-parameter capacity allows barcode detection to be combined with antibody-based phenotyping. Detection probes were conjugated to metal isotopes, enabling barcode readout to be integrated with standard antibody-based CyTOF panels in a single acquisition (Fig. 2A).

**Figure 2.**
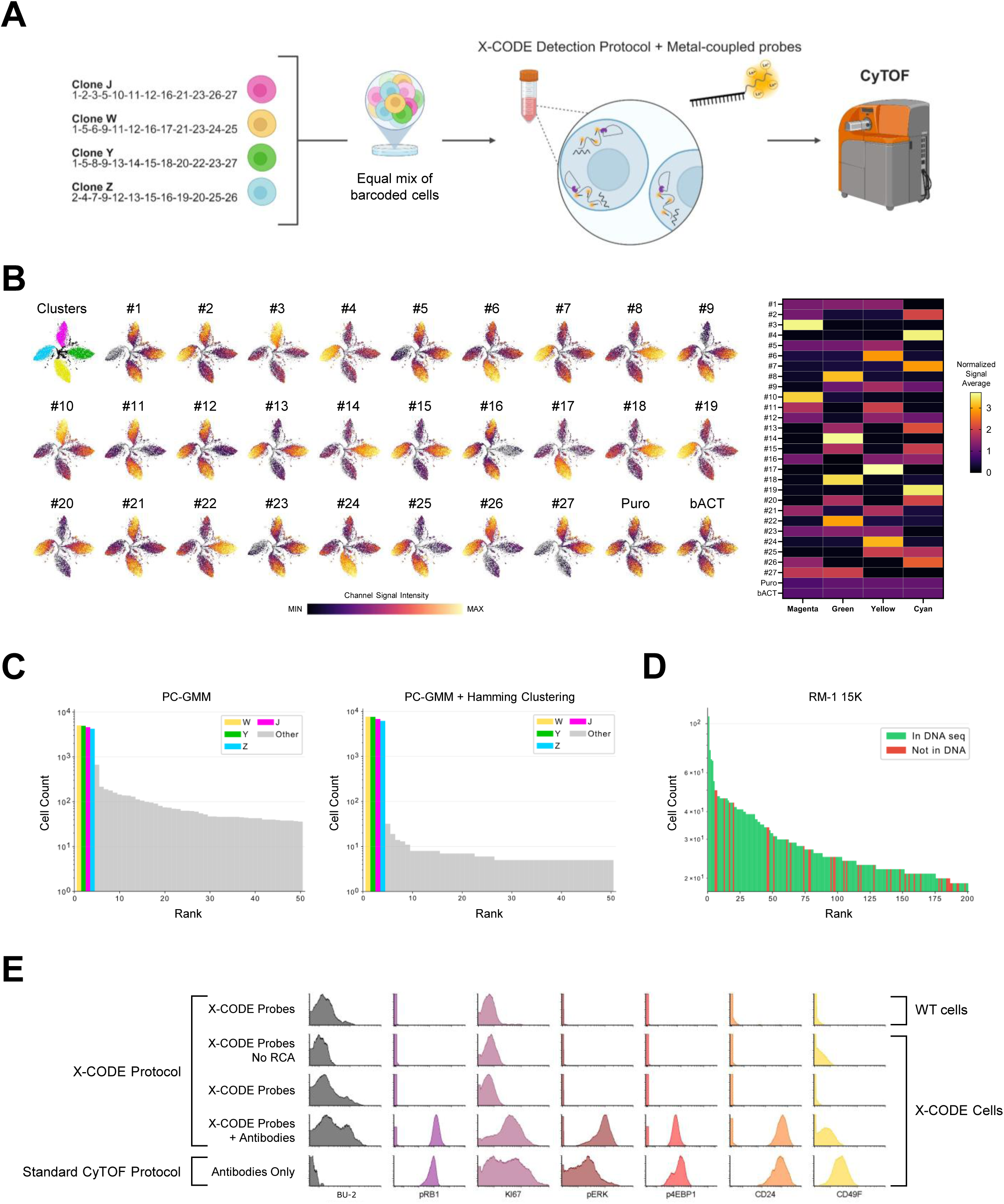
High-throughput X-CODE detection by CyTOF. **A**) Workflow illustrating validation of 2M BUs detection using the X-CODE probe-based protocol adapted for CyTOF (created with https://BioRender.com). Four barcoded clones (J, W, Y, Z) collectively represent the complete set of 27 BUs. The four clones were mixed equally and then analysed in a single run. Detection probes were conjugated in-house with metal isotopes to enable mass cytometry readout. **B**) t-SNE visualisation and cluster-specific heatmaps. Left: CyTOF data from the JWYZ mix acquisition were analysed using t-SNE, revealing four distinct cell clusters: magenta (top), cyan (left), yellow (bottom), and green (right). Each panel displays the same t-SNE, with colour intensity representing the signal intensity of a specific detection probe across the clusters. Right: heatmap displaying normalised average signal intensity for each detection probe across the four clusters. **C**) Hamming clustering improves barcode recovery in X-CODE 2M data. Left: Rank-count histogram for PC-GMM debarcoding before Hamming clustering (CTV: 0.030). Expected barcodes (J, W, Y, Z) are coloured; gray bars indicate non-expected patterns. Right: Rank-count histogram after sphere Hamming clustering (radius=4, ratio=2, min_count_center=500). CTV: 0.04. CTV = (1/2) Σi |pi - qi|, where pi is observed proportion and qi = 1/4 is expected uniform proportion. **D**) DNA concordance validates CyTOF debarcoding in 15K RM-1 dataset. Rank-count histogram for PC-GMM debarcoding with patterns coloured by DNA membership. Barcodes detected by PC-GMM (count ≥ 3: 3035), 162 matching DNA barcodes in the top 200. **E**) Histograms showing signal intensity distributions for the indicated markers under different processing conditions. Samples were prepared using, from top to bottom, (i) the full X-CODE protocol with X-CODE probes on WT cells, (ii) X-CODE probes on X-CODE cells without the RCA step, (iii) X-CODE probes on X-CODE cells with the complete X-CODE protocol, (iv) X-CODE probes together with antibodies using the full protocol on X-CODE cells, and (v) antibodies alone using the standard CyTOF workflow on X-CODE cells. The first panel represents a randomly selected BU target (BU-2); all remaining panels display antibody-derived signals.

Using the 2M X-CODE library, we assessed whether combinatorial barcode identities could be resolved in a pooled, single-run format. Four clonal RM-1-derived cell populations (J, W, Y and Z), together spanning all 27 barcode units, were mixed at equal proportions and analysed simultaneously. CyTOF analysis followed by dimensionality reduction using t-SNE revealed four distinct clusters (Fig. 2B). Projection of individual barcode unit signals onto the t-SNE embedding demonstrated clone-specific patterns, with each cluster defined by the expected combinatorial BU signature (Fig. 2B, right). Similar results were obtained using a lower-complexity X-CODE design in a different cell line. Three TRAMP-C1 clonal populations collectively spanning all 18 barcode units of the 15K library were profiled (Supplementary Fig. 2A), and barcode unit signals again segregated according to clone identity, with minimal background detected in wild-type cells and in no-RCA controls (Supplementary Fig. 2B).

To computationally debarcode CyTOF X-CODE data, we implemented two approaches: PC-GMM, a pattern-constrained Gaussian mixture model that fits two-component mixtures to each channel and selects the most likely valid 4-of-9 pattern; and an adaptation of PreMessa^49^, an iterative method that normalises channel intensities and selects the top-4 channels per block. We validated both methods on the 2M dataset containing four known barcodes with roughly uniform distribution. Initial debarcoding correctly identified all four expected patterns as the highest-abundance patterns but also produced numerous spurious low-count assignments arising from measurement noise (Fig. 2C, Supplementary Fig. 3A). Visualisation of pairwise Hamming distances revealed that spurious patterns cluster within small Hamming distances of the true barcodes (Supplementary Fig. 3B), motivating a postprocessing correction step. Application of Hamming clustering, which reassigns cells from low-count patterns to nearby high-count neighbours, consolidated assignments to the four expected barcodes and increased the fraction of cells assigned to expected patterns from 58.4% to 86.7% (Fig. 2C). To validate that our pipeline recovers true biological barcodes rather than computational artifacts, we analysed the 15K dataset where independent DNA sequencing provides ground truth. Hamming clustering was not applied to this dataset because the smaller barcode space increases the risk of true barcode neighbours, and simulations showed negligible improvement at this expected sub-library size. Rank-count analysis showed that the majority of high-abundance CyTOF-detected patterns correspond to NGS-verified barcodes for both methods (Fig. 2D, Supplementary Fig. 3C). Quantifying recovery across detection thresholds demonstrated that concordance far exceeds the hypergeometric null expectation, confirming genuine signal rather than chance (Supplementary Fig. 3D). Overall, PC-GMM detected 3,035 unique patterns, of which 1,701 (56%) were independently verified by DNA sequencing, representing 49% recall of the 3,493 NGS-confirmed barcodes (Supplementary Fig. 3E). Benchmarking on simulated data confirmed PC-GMM accuracy gains over PreMessa, particularly when channels exhibit bimodal distributions (Supplementary Fig. 3F, 3H), and that conservative Hamming clustering improves performance with minimal harmful reassignments (Supplementary Fig. 3G).

Finally, we examined whether X-CODE barcode detection could be integrated with conventional CyTOF immunophenotyping without compromising assay performance. Barcode signals were dependent on the rolling circle amplification step and absent in wild-type controls, while antibody-derived signals remained comparable whether acquired alone or together with X-CODE probes (Fig. 2E). These data indicate that X-CODE readout is compatible with standard antibody staining and CyTOF acquisition parameters.

Together, these analyses demonstrate that X-CODE supports high-throughput, combinatorial barcode detection by mass cytometry across multiple library scales while remaining compatible with simultaneous, high-dimensional protein phenotyping.

### Imaging-based X-CODE detection across cells, tissues, and MALDI-compatible workflows

Having established high-dimensional, sequencing-free X-CODE barcode detection in suspension using mass cytometry, we next extended this approach to spatially resolved imaging. To this end, X-CODE barcode detection was implemented using an established high-plex spatial imaging platform by repurposing the Akoya PhenoCycler-Fusion for direct RNA barcode readout. In contrast to its standard use, in which oligonucleotide-conjugated antibodies are detected through iterative hybridisation of fluorescent probes, the antibody detection oligos were replaced with fluorescent X-CODE detection probes designed to recognise rolling circle amplification amplicons generated by the X-CODE protocol. These fluorescent probes were processed using the unmodified Akoya PhenoCycler-Fusion fluidics, cycling, and imaging workflow, without altering instrument hardware or acquisition parameters (Fig. 3A, top).

**Figure 3.**
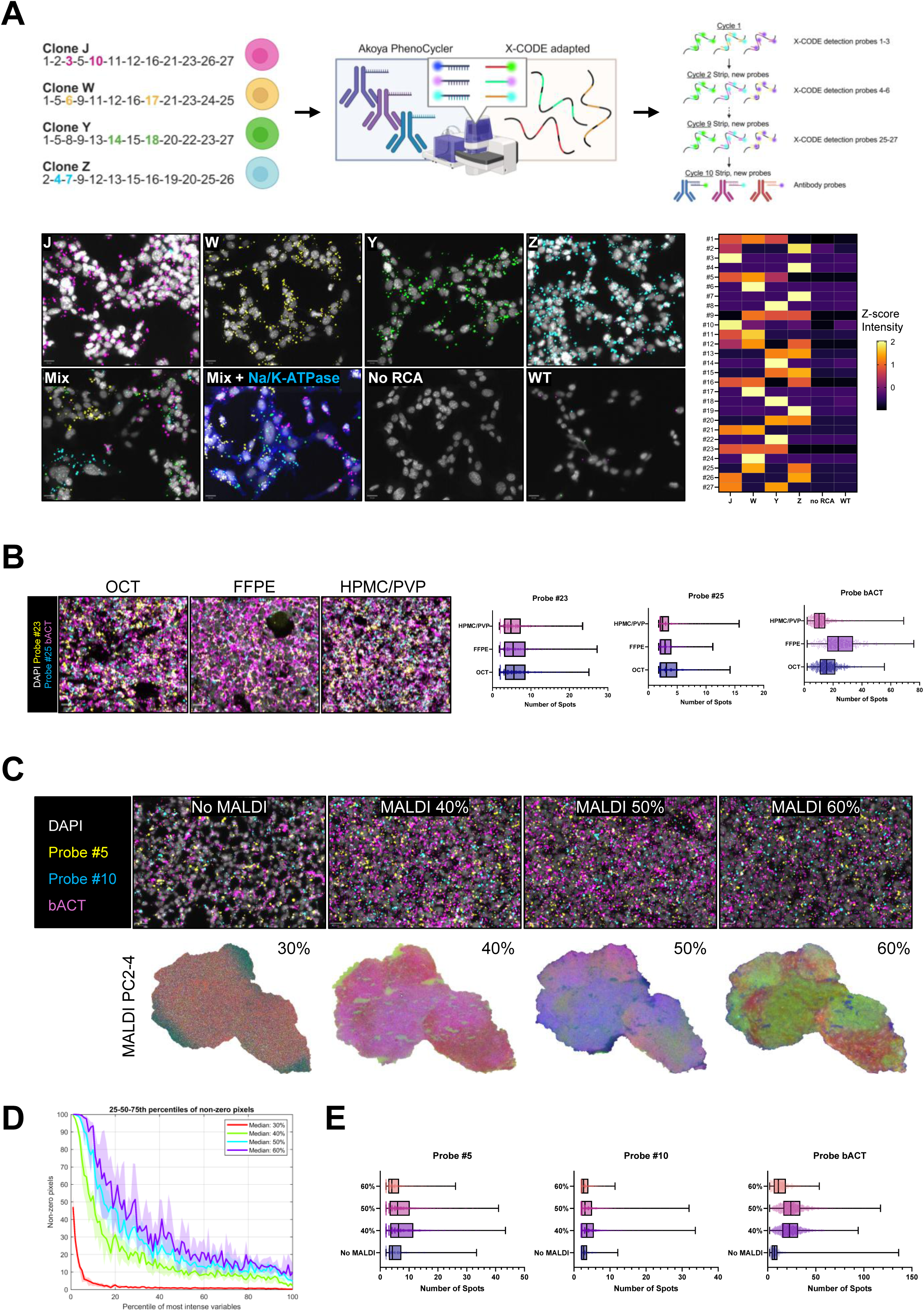
Imaging-based detection across cells, tissues and MALDI compatibility. **A)** Top: Schematic overview of the spatial X-CODE workflow adapted for the Akoya PhenoCycler platform (created with https://BioRender.com). Four barcoded clones (J, W, Y, Z) collectively represent the complete set of 27 BUs. Each clone is uniquely identified by two BUs and represented by a single colour: J (magenta), W (yellow), Y (green) and Z (cyan). Bottom: Representative fluorescence images of J, W, Y and Z barcoded cells across eight experimental conditions: J, W, Y, Z, WT, no-RCA, mix, and mix + Na/K-ATPase antibody. The no RCA control was processed without rolling circle amplification. The WT sample consists of untagged RM-1 cells. The mix sample contains an equal mixture of the four clones. The mix + Na/K-ATPase condition combines the X-CODE protocol with antibody staining and standard Akoya detection probes. In all conditions, each barcode is visualised by two BUs, uniquely colour-coded as in the schematic above. The heatmap displays Z-score normalised average spot intensity per cell for each probe across samples. **B)** Representative fluorescence images of mixed RM-1 J, W, Y, Z barcoded tumours from C57BL/6 mice processed using three tissue embedding methods: OCT, FFPE, and HPMC/PVP hydrogel (hydroxypropyl-methylcellulose/polyvinylpyrrolidone enriched). Nuclei are shown in white (DAPI), probe #23 in yellow, probe #25 in cyan, and β-actin in magenta. Box plots (right) display spot counts per probe per cell across embedding methods. **C)** Detection of the RM-1-15K library tumours post MALDI-MSI processing and acquisition. Top: Representative X-CODE spatial images of RM-1-15K tumours from C57BL/6 mice processed with MALDI at different laser power settings (No MALDI, 40%, 50%, 60%). DAPI in white, probe #5 in yellow, probe #10 in cyan, and β-actin in magenta. Below: MALDI PCA composite maps (PC2–4) for the same samples, highlighting spectral variance patterns. **D**) Sparsity distribution, where variables are ranked by intensity and grouped by centile. The X-axis represents top-ranked variables, and the Y-axis indicates the percentage of non-zero pixels, reflecting signal occupancy. **E**) Box plots show the number of detected spots per probe across conditions.

Using this approach, we first assessed barcode detection in cultured cells. Four clonal RM-1-derived X-CODE 2M populations (J, W, Y and Z), previously used for CyTOF analysis, were analysed. Each line carries a unique barcode encoded by a combination of 12-of-27 BUs, visualised in this defined mixture using two distinguishing barcode units per clone (Fig. 3A, top). In single-clone samples, barcode signals were robust and restricted to the expected line, while analysis of the mixed population enabled correct spatial resolution of all four clones (Fig. 3A bottom). Barcode signal was minimal in wild-type and no-RCA controls, and X-CODE detection was compatible with simultaneous antibody staining, exemplified by Na/K-ATPase co-detection. Quantification of Z-score–normalised spot intensity per cell confirmed clear BU-specific signal patterns across samples (Fig. 3A), while per-probe spot counts across conditions further demonstrated specificity and low background (Supplementary Fig. 4A). Similar results were obtained using imaging mass cytometry (IMC) in plated cells tagged with the X-CODE 15K library, where barcode detection was achieved using the same metal-conjugated probes employed for CyTOF analysis, demonstrating consistent spatial resolution and signal specificity across platforms (Supplementary Fig. 4B).

We next evaluated X-CODE performance in tissue sections and across standard embedding methods. Barcoded RM-1 J, W, Y and Z cells were used to generate tumours in immunocompetent mice, and tumour sections were processed using OCT, FFPE, or HPMC/PVP hydrogel embedding, the latter selected for compatibility with mass spectrometry imaging^50^. Across all embedding conditions, barcode signals for representative probes and the β-actin control were consistently detected by both fluorescence imaging and IMC, with spot counts and signal intensities varying between methods but remaining quantifiable and interpretable, as reflected by spot count distributions and mean spot intensity per cell (Fig. 3B; Supplementary Fig. 4C, D). Based on these results, we further compared OCT and HPMC/PVP hydrogel embedding in the context of combined X-CODE detection and protein staining using the Akoya PhenoCycler-Fusion. Incorporation of the antibody staining workflow did not interfere with X-CODE detection, and both embedding methods supported robust barcode readout (Supplementary Fig. 5A). Antibody signals were comparable between embeddings; however, individual antibodies showed variable performance, indicating that marker-specific optimisation may be required for reliable co-detection of barcodes and proteins on the same tissue section (Supplementary Fig. 5B).

Having established compatibility with spatial imaging across embedding conditions, we next explored the use of X-CODE in combination with matrix-assisted laser desorption/ionisation (MALDI) mass spectrometry imaging as an upstream application. MALDI provides label-free, spatially resolved molecular information, such as metabolites and lipids, that is orthogonal to antibody-based phenotyping, creating the opportunity to align spatial clonal architecture with underlying molecular landscapes within the same tissue section.

To assess compatibility between these modalities, four tissue sections were analysed by MALDI using increasing laser power settings prior to X-CODE detection. Preprocessing was performed to identify (approximately 1600) peaks across all 4 samples, and PCA visualisations demonstrate that laser powers ≥50% yielded coherent spatial patterns, whereas lower powers were less informative (Fig. 3C). The primary reason for this is related to the higher image sparsity at lower laser powers (Fig. 3D; Supplementary Fig. 5C); the most intense 1% variables have a median sparsity of <50% at the lowest laser power thus demonstrating insufficient ionisation across the sample. As the power increases, the variables become less sparse, as a result of increased ionisation. We then evaluated whether X-CODE probe detection remained effective following MALDI acquisition by measuring two representative probes across conditions on the same tissue section. Quantification of spot counts and mean probe intensity indicated that 50–60% laser power preserved X-CODE probe detection (Fig. 3E; Supplementary Fig. 5D). Together, these data support ≥50% laser power as a practical operating regime for combining MALDI imaging with X-CODE probe detection.

Together, these results demonstrate that X-CODE supports spatially resolved RNA barcode detection using repurposed Akoya PhenoCycler-Fusion hardware and IMC across cells and tissues, and that this workflow is compatible with MALDI mass spectrometry imaging as an upstream application, enabling integration of clonal identity with spatial molecular profiling on the same tissue section.

### X-CODE reveals clonal spatial organisation underlying tumour evolution during androgen deprivation *in vivo*

To apply X-CODE to an *in vivo* therapeutic setting, we investigated clonal dynamics and spatial phenotypes during androgen deprivation in the androgen-sensitive MyC-CaP mouse prostate cancer model. MyC-CaP cells were barcoded with the X-CODE 15K library, implanted into immunocompetent mice, and tumours were sampled longitudinally before and after chemical castration induced by Degarelix. Tumours were processed to maximise downstream flexibility, with portions allocated for DNA extraction, cell culture and line establishment, and the central tumour region split and embedded in OCT and HPMC/PVP hydrogel for spatial analyses. This experimental design enabled integrated, longitudinal analysis of clonal dynamics across multiple readouts using matched samples, providing a defined temporal window to examine clonal behaviour during treatment response and the emergence of castration-resistant disease (Fig. 4A). Consistent with the known androgen dependence of the MyC-CaP model^51^, tumour growth was suppressed following castration, with an initial phase of tumour regression followed by eventual relapse, whereas tumours in untreated mice continued to grow progressively (Fig. 4B).

**Figure 4.**
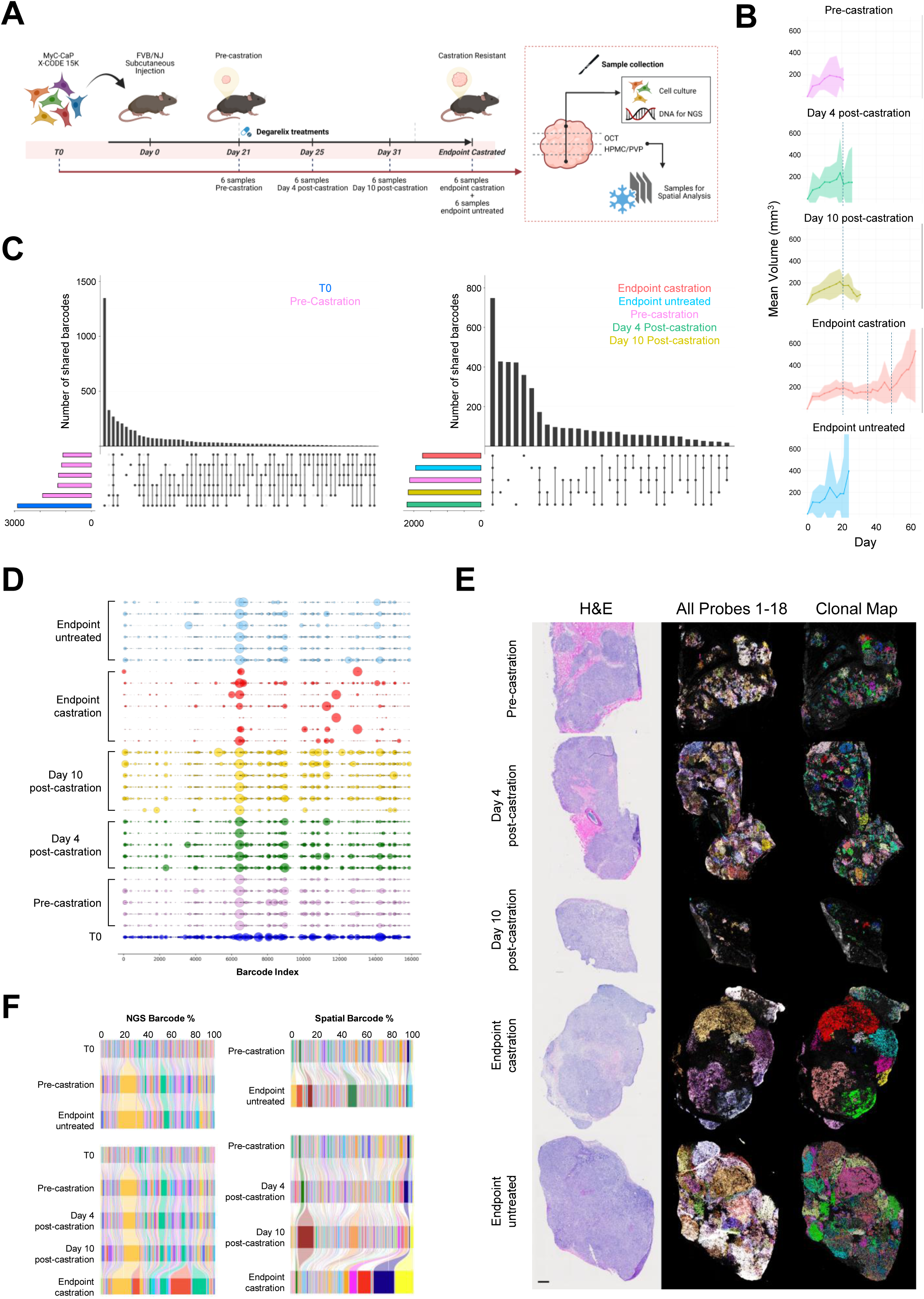
X-CODE resolves clonal dynamics during androgen deprivation in vivo. **A**) Experimental design for longitudinal clonal tracking in the MyC-CaP model (created with https://BioRender.com). MyC-CaP cells transduced with the 15K X-CODE library (T0) were injected subcutaneously into FVB/NJ mice. Tumours were collected at baseline prior to chemical castration with degarelix, at 4 days and 10 days post-castration, and at endpoint under castrated or untreated conditions. Tumours were sectioned for parallel DNA barcode sequencing and spatial analysis. **B**) Tumour growth curves across experimental groups showing mean tumour volume (n=5-6) over time for each experimental group. Shaded areas indicate 95% confidence intervals. **C**) UpSet plots illustrating barcode sharing across samples. Left panel: Intersection of unique barcodes between T0 (blue; MyC-CaP-15K cells before injection) and pre-castration samples (pink). Right panel: Intersection of unique barcodes shared across pooled samples from all timepoints, including T0, Pre-castration, Day 4 post-castration (green), Day 10 post-castration (yellow), Endpoint castration (red), and Endpoint untreated (light blue). **D**) Bubble plots showing barcode composition of all tumour samples across different timepoints, based on DNA sequencing data after debarcoding. Each row represents an individual sample, and bubbles indicate the relative abundance of each barcode (indexed along the x-axis). Samples are grouped and color-coded by timepoint: T0 (blue) – MyC-CaP-15K cells before injection; Pre-castration (purple); Day 4 post-castration (green); Day 10 post-castration (yellow); Endpoint castration (red); and Endpoint untreated (light blue). **E**) Representative tumour sections from samples across the experimental groups. Left: H&E staining showing overall tumour morphology. Middle: combined fluorescent signal, acquired using Akoya PhenoCycler, from all 18 X-CODE probes detecting the 15K library. Right: spatially resolved clonal maps following computational debarcoding, illustrating distinct spatial clonal distribution within tumours. Scale bar, 400 µm. **F**) Alluvial plots showing barcode composition dynamics. Each bar represents the normalised barcode frequencies calculated from pooled data across all tumour samples within the corresponding experimental group. Left panels: barcode composition derived from debarcoding of DNA sequencing (NGS) data. Right panels: barcode frequencies derived from debarcoding of spatial barcode data.

We first characterised clonal dynamics using DNA-level barcode sequencing across all tumours and time points. Comparison of barcode composition between the initial *in vitro* population (T0) and pre-castration tumours revealed that only a subset of barcoded clones reproducibly engrafted *in vivo*, with 369 barcodes shared across all pre-castration tumours, indicating early selection for clones with superior tumour-establishing capacity (Fig. 4C). Analysis across all subsequent samples identified 748 unique barcodes shared across pooled time points, demonstrating robust barcode recovery and enabling longitudinal tracking of clonal lineages across the experiment (Fig. 4C).

Analysis of clonal dynamics showed that barcode composition was highly similar across pre-castration, early post-castration (day 4 and day 10), and untreated endpoint samples (Fig. 4D). In contrast, castration-resistant endpoint tumours displayed pronounced clonal remodelling, characterised by the selective expansion of a limited subset of clones. This pattern is further supported by pairwise correlation analysis of barcode composition, which shows high similarity among non-resistant samples and a distinct profile in castration-resistant tumours (Supp. Fig. 6). Importantly, these expanding clones were not prominently enriched at early post-castration time points, indicating that the observed resistance is not associated with early selective outgrowth immediately following castration, but instead reflects differential clonal fitness under sustained androgen deprivation, consistent with differences in adaptive potential among clonal lineages.

Based on the clonal dynamics identified by sequencing, selected samples were subjected to spatial profiling using HPMC/PVP-embedded tumour sections (Fig. 4E). Following cell segmentation, subcellular spots corresponding to all 18 barcode units were quantified and barcode identities assigned using the 15K whitelist (Table 7 – 15K_Key). Cells with minor barcode unit mismatches that were spatially proximal to confidently assigned cells were grouped accordingly, enabling reconstruction of spatially resolved clonal maps (Fig. 4E). Pre-castration and untreated tumours exhibited highly clonal organisation characterised by numerous small clonal patches arranged in close proximity, with frequent spatial intermixing among neighbouring lineages. In contrast, castration-resistant tumours were organised into large tumour regions dominated by a limited number of clones, resulting in pronounced spatial segregation. This reorganisation of clonal structure mirrors the strong clonal selection observed by sequencing and indicates that resistant disease is sustained by the local expansion of a restricted set of adapted clonal lineages following androgen deprivation (Fig. 4E).

Comparison of clonal dynamics inferred from DNA sequencing and spatial debarcoding showed highly concordant patterns of tumour evolution in response to castration (Fig. 4F). In untreated tumours, barcode composition remained largely stable over time, indicating limited clonal selection during continued androgen-driven growth. In contrast, castration induced pronounced but delayed clonal selection: while barcode diversity was largely preserved at early post-castration time points, castration-resistant endpoint tumours were dominated by a restricted subset of barcodes, consistent with clonal outgrowth following a period of adaptation rather than immediate selection.

Together, these results establish X-CODE as a flexible, end-to-end platform for studying tumour evolution *in vivo*, enabling high-throughput clonal tracking across large cohorts and time points, informed samples selection for spatial analysis, and integrated resolution of clonal identity during therapy response.

### Clone-associated phenotypic and metabolic states during tumour evolution under androgen deprivation

Following characterisation of clonal dynamics, we integrated spatial clonal maps with multimodal tumour phenotyping to define phenotypic and metabolic states associated with dominant clones during androgen deprivation. Consecutive sections from the same HPMC/PVP-embedded tumour blocks were analysed to enable direct spatial alignment of clonal identity, multiplex immunophenotyping, and spatial metabolomics. Multiplexed imaging using the Akoya PhenoCycler-Fusion revealed marked differences in tumour architecture and microenvironmental organisation across treatment conditions (Fig. 5A). Pre-castration and untreated tumours were phenotypically homogeneous, displaying a uniform CK8⁺ luminal epithelial phenotype with variable but broadly distributed stromal and immune components. In contrast, castration-resistant endpoint tumours exhibited pronounced spatial heterogeneity, with discrete tumour regions showing reduced luminal marker expression and relative exclusion of immune and stromal cells, which instead accumulated in the areas between clonal regions. Whole-tumour quantification revealed a transient enrichment of CD8⁺ T cells at day 10 post-castration, whereas a castration-resistant tumour was characterised by enrichment of myeloid populations (Fig. 5B). Analysis of AR and Ki67 showed the expected temporal pattern following androgen deprivation, with reduced proliferation and cytoplasmic re-localisation of AR at early post-castration time points, and re-emergence of proliferative activity in resistant tumours. Notably, within a castration-resistant tumour dominated by three clones, distinct spatially resolved AR and proliferation states could be observed, including AR-negative proliferative regions, AR-cytoplasmic proliferative regions, and AR-cytoplasmic low-proliferative regions, indicating clone-associated modes of adaptation (Fig. 5A bottom panel). Whole-tumour analysis further revealed increased basal marker expression and synaptophysin positivity in castration-resistant samples, consistent with phenotypic plasticity under sustained androgen deprivation (Fig. 5B).

**Figure 5.**
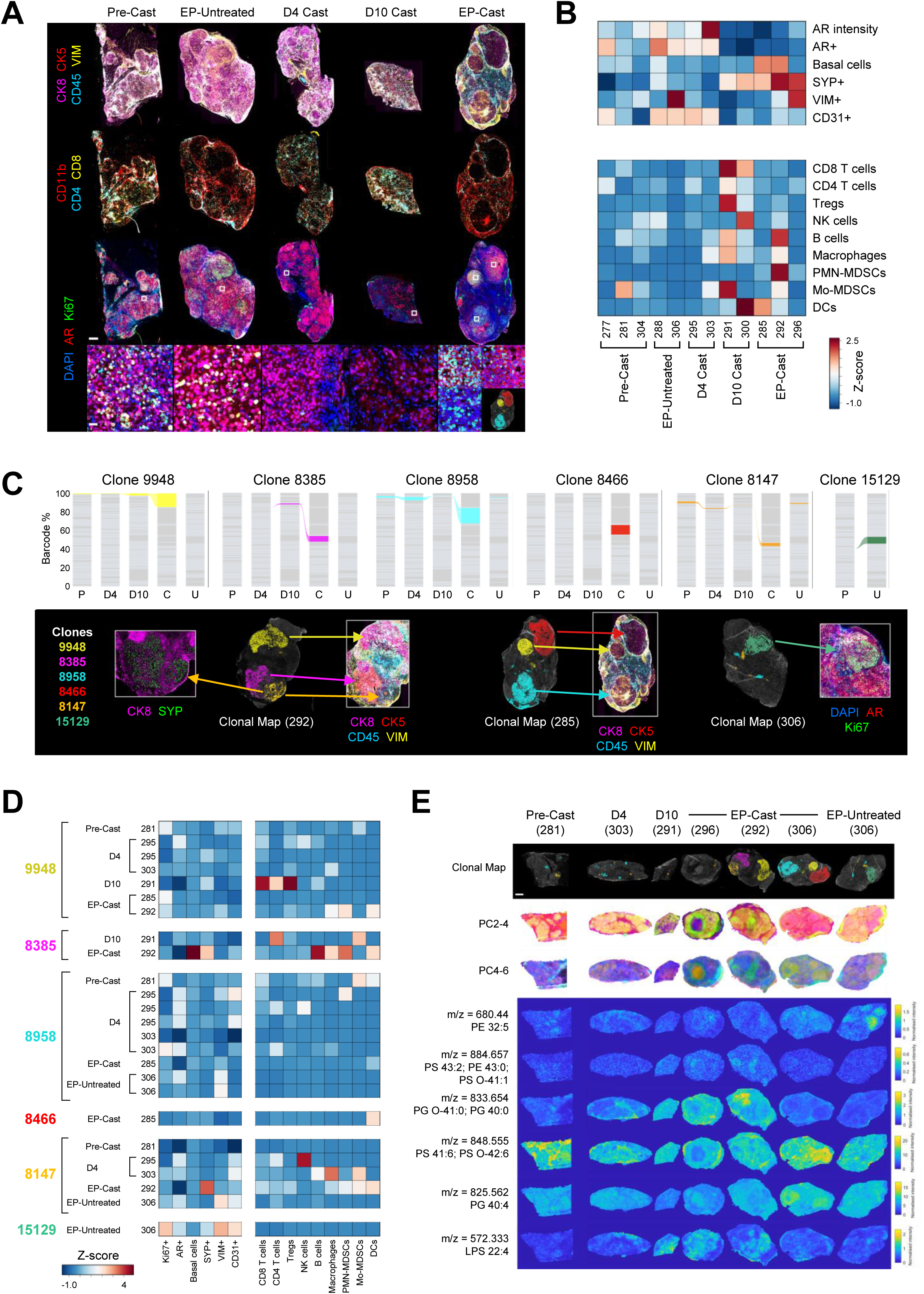
Clone-associated phenotypic and metabolic states during tumour evolution under androgen deprivation. **A**) Multiplexed imaging of representative tumour sections using the Akoya PhenoCycler. Sections were stained for epithelial (CK8, CK5), immune (CD45), stromal (VIM), androgen receptor (AR), and proliferation (Ki67) markers across experimental conditions: pre-castration (Pre-Cast), untreated endpoint (EP-Untreated), 4 days post-castration (D4 Cast), 10 days post-castration (D10 Cast), and castration endpoint (EP-Cast). Scale bar, 500 µm. Bottom panels show higher-magnification views of selected regions highlighting AR and Ki67 staining. Scale bar, 20 µm. Bottom-right panel shows the spatial clonal map of the three dominant clones identified in the EP-Cast tumour. **B**) Heatmaps summarising whole-tumour phenotypic and immune profiles across all analysed samples and experimental groups. Top: CD45-negative (non-immune) populations. Bottom: CD45-positive immune cell populations. Values represent per-feature Z-score normalisation across samples. **C**) Clonal dynamics across time points for selected clones of interest. Top: alluvial plots depicting the relative abundance of selected clones (IDs: 9948, 8385, 8958, 8466, 8147, and 15129) based on pooled data for each experimental group. Bottom: representative multiplexed images highlighting the spatial distribution and associated phenotypic features of clones of interest in treated tumours (292, 285) and an untreated tumour (306). **D**) Heatmaps summarising clone-specific phenotypic and immune profiles across experimental groups. Top: CD45-negative (non-immune) populations. Bottom: CD45-positive immune cell populations. Values represent per-feature Z-score normalisation across samples. **E**) Integration of spatial metabolomics with clonal maps (top; clone colours as in Fig. 5C). Upper panels show MALDI-derived principal component (PC) maps aligned to tumour sections across time points. Lower panels display representative ion images corresponding to clone-associated m/z features with putative annotations, illustrating spatially localised metabolic variables associated with the clones of interest. Scale bar, 800 µm.

**Figure 6.**
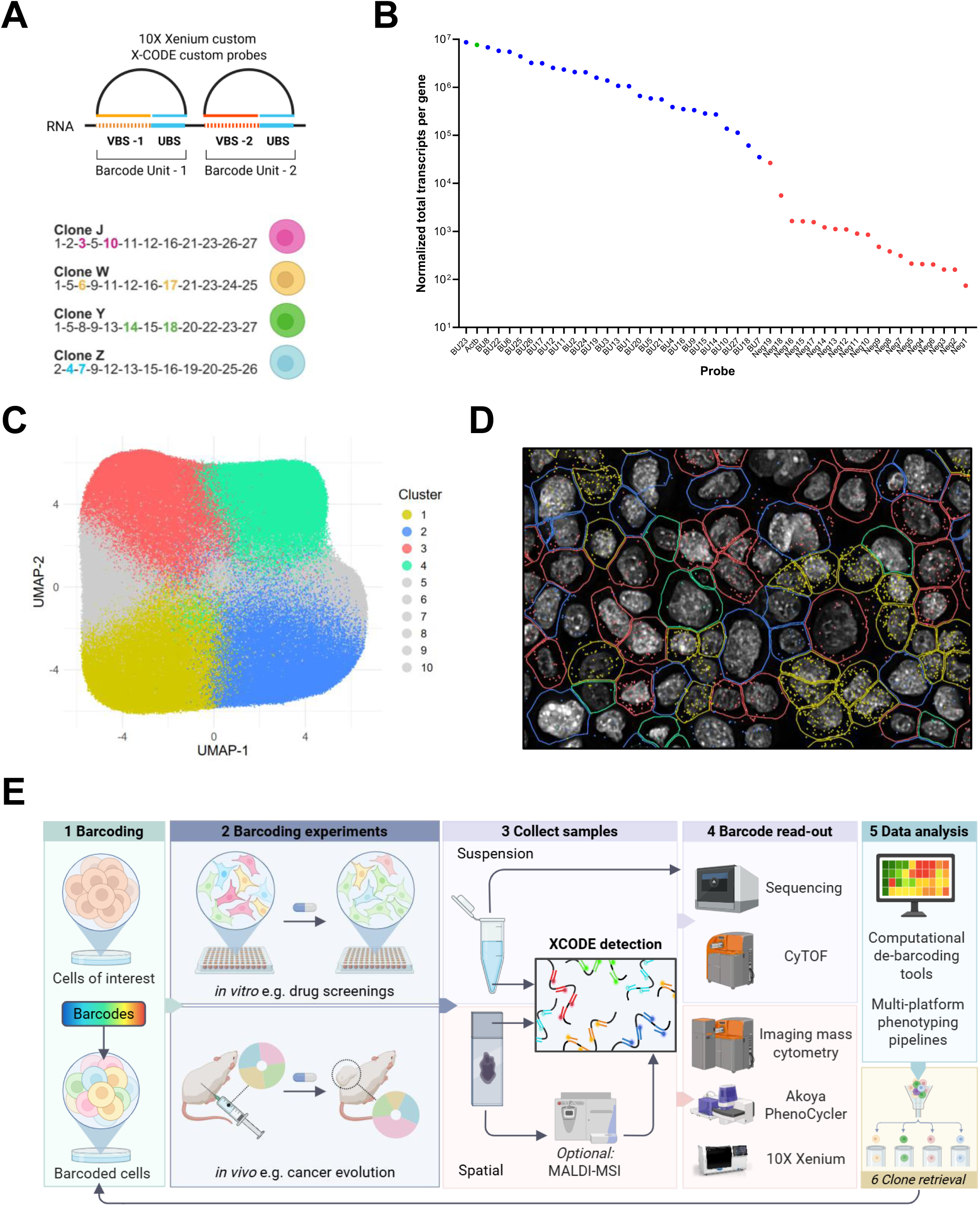
X-CODE compatibility with 10X Xenium spatial transcriptomics. **A**) Schematic showing integration of X-CODE barcoding with 10x Genomics Xenium probes. Each probe targets the entire barcode unit, with the ligation point positioned at the boundary between the variable barcode sequence (VBS) and the universal barcode sequence (UBS). 5’ RNA-binding domain (RBD) and 3’ RBD span between 16-23 bp. Four barcoded clones (J, W, Y, Z) collectively represent the complete set of 27 BUs. Each clone is uniquely identified by two BUs and represented by a single colour: J (magenta), W (yellow), Y (green) and Z (cyan). **B**) Probe performance across barcode units. Normalised detection counts per probe are plotted for all barcode probes of the 15K library, highlighting detection above negative controls. Normalised detection counts for each BU = (BU total transcript count) ÷ (number of times the BU appears in J, W, Y, Z, clones). **C**) UMAP visualization of spatial transcriptomic clusters obtained from equally mixed J, W, Y, Z cells STAMPed on a 10x Xenium slide. Cells are grouped into 10 clusters based on Xenium spatial transcriptomic data, illustrating distinct transcriptional profiles. Clusters 1–4 include the majority of cells and spot analysis (D) reveals that they correspond to clones W (yellow), J (blue), Y (red), and Z (light green), respectively. **D**) Representative 10X Xenium image showing cell segmentation boundaries and barcode probe signals overlaid on tissue architecture, enabling spatial localization of barcoded clones. Clone J (BU 3 and BU 10) is shown in blue, Clone W (BU-6 and BU-17) in yellow, Clone Z (BU-4 and BU-7) in light green, and Clone Y (BU-18 and BU-22) in red. **E**) Workflow for X-CODE integration with multi-platform analysis. Overview of the experimental pipeline: (1) barcoding cells of interest, (2) performing *in vitro* or *in vivo* experiments, (3) collecting samples, (4) barcode read-out via sequencing or probe-based detection platforms (e.g., Xenium, imaging mass cytometry, Akoya PhenoCycler, CyTOF), (5) computational debarcoding and multi-platform phenotyping and (6) clonal retrieval and functional characterization.

Guided by clonal dynamics, we next focused on individual clones with detection across time points and enrichment at either untreated or castration-resistant endpoints (Fig. 5C,D). Clones enriched in castration-resistant tumours displayed distinct spatially resolved phenotypic associations. Clone 9948, dominant in two resistant tumours, exhibited divergent states across tumours, ranging from luminal regions with associated myeloid infiltration to immune-excluded regions with reduced CK8 expression. Clone 8385 showed a pronounced basal phenotype with increased synaptophysin expression and myeloid infiltration, features not apparent at earlier time points. Clone 8958, although detected across multiple conditions, displayed a selective increase in VIM-positive cells at the resistant endpoint. Clone 8466, detected exclusively at castration-resistant endpoint, exhibited marked loss of epithelial markers together with the strongest induction of synaptophysin. In contrast, clone 15129, detected only in untreated endpoint tumours, was highly proliferative, immune excluded, and associated with increased stromal and vascular features. These observations indicate that MyC-CaP dominant clones are associated with distinct spatial phenotypic configurations rather than converging on a single resistant state.

To further characterise these clone-associated states, we performed MALDI mass spectrometry imaging on matched tumour sections spanning pre-castration, early post-castration, untreated endpoint, and castration-resistant endpoint samples, and co-registered metabolic profiles with spatial clonal maps (Fig. 5E). Principal component analysis revealed tumour- and clone-specific metabolic patterning, with spatially localised features aligning with clonal regions. For each dominant castration-resistant clone, we identified m/z signals selectively enriched within that clone relative to others (Fig. 5E; Supp. Fig. 7). These clone - associated features corresponded to lipid species with putative annotations including phosphatidylserines, phosphatidylglycerols, phosphatidylinositols, ceramide-linked phosphoethanolamines, and lysophospholipids. Different dominant clones were associated with distinct lipid signatures, indicating that clonal dominance is compatible with multiple, non-overlapping metabolic configurations linked to membrane composition, signalling, and stress response pathways.

Together, these data demonstrate that prostate cancer evolution under androgen deprivation does not converge on a single phenotypic or metabolic programme. Instead, castration - resistant tumours comprise multiple dominant clones, each occupying distinct spatial niches and associated with specific phenotypic and metabolic states. These findings support a model in which resistance reflects multiple evolutionary solutions to high fitness, shaped by clonal adaptive potential and microenvironmental context rather than uniform metabolic convergence. X-CODE enables direct resolution of this heterogeneity in situ, integrating clonal identity with spatial phenotype and metabolism during tumour evolution under therapy.

### Compatibility of X-CODE with probe-based spatial transcriptomics using 10x Xenium

While mass cytometry and multiplex immunofluorescence enable high-throughput, sequencing-free readout of X-CODE barcodes, these platforms are inherently constrained by the limited number of detectable parameters. In practice, a substantial fraction of available channels must be allocated to X-CODE probes (18 for the 15K library and 27 for the 2M library), leaving limited capacity for phenotypic characterisation and necessitating strong *a priori* selection of antibody panels. By contrast, sequencing-based spatial transcriptomic approaches provide substantially richer phenotypic information but remain costly, can suffer from incomplete barcode recovery, and typically do not achieve true single-cell resolution in their most established implementations. Probe-based spatial transcriptomic platforms such as 10x Genomics Xenium address several of these limitations by enabling single-cell–resolved, highly multiplexed RNA detection using predefined but extensible probe panels comprising hundreds to thousands of targets. Importantly, Xenium relies on a detection chemistry based on padlock probe hybridisation, ligation, and rolling circle amplification, conceptually aligned with the X-CODE detection strategy. We therefore explored the compatibility of X-CODE with Xenium as a route to combine high-complexity clonal barcoding with rich, spatially resolved transcriptomic phenotyping. Because Xenium panels cannot practically accommodate the millions of unique sequences used in conventional barcode libraries, this approach enables X-CODE to function as a compact, probe-detectable barcode system compatible with probe-based spatial transcriptomics.

To enable detection of X-CODE barcodes using Xenium, we adapted the X-CODE detection strategy to match the Xenium probe architecture. In Xenium, target detection relies on a single probe design in which the 5′ and 3′ probe arms hybridize to adjacent regions of the target RNA, bringing the probe ends into close proximity in an inverted U-shaped configuration that enables ligation and subsequent rolling circle amplification. Because this architecture differs from the dual-probe design used for other X-CODE readouts, we generated a custom Xenium probe panel by redesigning all BU hybridization probes to conform to the Xenium chemistry (Fig. 5A, top).

We validated this approach using the same four clonal RM-1-derived X-CODE 2M populations (J, W, Y, and Z) used in previous experiments. These clones were mixed, attached onto Xenium slides using the STAMP protocol^52^, and processed using the custom X-CODE Xenium panel in combination with a standard 10x Xenium mouse gene expression panel. Inclusion of this panel ensured that X-CODE probes were evaluated under conditions representative of a typical Xenium experiment (Fig. 5A, bottom).

Quantification of X-CODE probe performance demonstrated robust and specific detection of all 27 BU probes, which consistently outperformed negative control probes when assessed by normalised transcript counts (Fig. 5B). These data indicate that the redesigned BU probes are efficiently detected by the Xenium chemistry and are compatible with standard Xenium workflows.

To assess whether barcode identity could be recovered in an unbiased manner, we analysed the combined dataset comprising both X-CODE probes and the Xenium mouse gene panel using the Xenium software pipeline. UMAP-based dimensionality reduction identified ten transcriptionally distinct cell clusters, four of which were dominant in the dataset (Fig. 5C). Importantly, spatial projection of barcode-specific BU signals onto these clusters revealed that each dominant cluster was selectively enriched for the barcode units corresponding to one of the four X-CODE barcodes (J, W, Y, or Z), with colours matching the expected clone identities (Fig. 5D). Thus, unbiased clustering of Xenium data correctly resolved the four clonal populations based solely on barcode detection, demonstrating specific and robust barcode recovery within a highly multiplexed spatial transcriptomic dataset.

In summary, X-CODE provides a flexible framework that supports clonal barcoding across experimental contexts and enables barcode readout using complementary detection platforms, including sequencing-based screening and probe-based, single-cell spatial transcriptomics (Fig. 5E). Because long, probe-detectable barcodes and short sequencing barcodes are intrinsically paired within the same construct, clonal identities can be directly integrated across sequencing and non-sequencing readouts, enabling coordinated analysis of clonal dynamics, spatial organisation, and phenotype within a unified experimental workflow. Cells of interest are first barcoded using X-CODE and subjected to experimental perturbations *in vitro* or *in vivo*. Samples can then be analysed using either sequencing-based approaches or probe-based detection platforms, including mass cytometry, imaging-based antibody platforms, or probe-based spatial transcriptomics. Across all probe-based modalities, barcode detection begins with the same set of BU-specific hybridisation probes (18 plus one universal probe for the 15K library, or 27 plus one for the 2M library), followed by platform-specific detection probe-sets and reagents. Importantly, X-CODE also supports barcode-specific clonal retrieval from collected samples, enabling functional follow-up of clones identified by either sequencing or spatial analysis.

## DISCUSSION

Understanding how cell populations evolve during development and under selective pressure requires experimental systems that can resolve lineage identity alongside phenotypic state and spatial context. While lineage tracing has become central to developmental and cancer biology^53^, existing approaches have imposed practical trade-offs between clonal resolution, phenotypic depth, spatial information, and experimental scalability^31,54–56^. In this study, we introduce the RNA barcode X-CODE as a unifying framework for high-complexity clonal tracking across sequencing-based, cytometric, and spatial imaging platforms within a single, coherent experimental design.

Current spatially resolved barcoding strategies span a range of complementary approaches, each optimised for specific experimental objectives^40,54,56,57^. Protein-based barcoding systems enable spatial detection through fluorophore-conjugated antibodies, while probe-based RNA barcodes have enabled spatially resolved optical pooled CRISPR screens and detection of evolvable barcodes, providing powerful tools for functional perturbation studies and dynamic lineage recording, respectively. However, practical bottlenecks differ across implementations, with trade-offs between library scale, probe number, imaging complexity, and analytical requirements shaping their applicability to large-scale clonal analyses.

X-CODE addresses several of these challenges through a dual-expressed RNA barcoding strategy, in which a long barcode is optimised for probe-based detection and a matched short barcode enables sequencing-based readout and barcode-specific clonal retrieval of the same clonal identities. Even at the highest library complexity tested (∼2 million barcodes), this design requires only 55 probes across hybridisation and detection, maintaining practical scalability for spatial and cytometric applications. The fixed correspondence between long and short barcodes enables unambiguous clone identification across probe-based and sequencing workflows, while the use of a defined barcode whitelist improves demultiplexing accuracy and analytical robustness relative to fully random barcode libraries.

A key advance of X-CODE is its broad compatibility across orthogonal platforms. We demonstrate efficient barcode detection and debarcoding by mass cytometry, establishing the feasibility of high-throughput clonal quantification alongside deep phenotypic profiling. Given the similar parameter space achievable with emerging spectral flow cytometry platforms, this strategy is, in principle, extensible beyond mass cytometry. More broadly, sequencing-free clonal analysis at this scale provides sensitivity to rare populations and offers particular advantages in contexts such as liquid malignancies, where clonal structure and phenotypic diversity have historically been studied using flow-based approaches.

We further show that X-CODE is compatible with spatial imaging workflows and tissue processing conditions required for MALDI mass spectrometry imaging, enabling direct co - registration of clonal maps with spatial metabolomic profiles from the same tissue sections. While MALDI-based metabolomics combined with RNA probe detection has been reported previously^42^, our work extends these efforts by demonstrating parallel spatial detection of high-complexity RNA barcodes using the Akoya PhenoCycler-Fusion platform, repurposed for RNA detection, which to our knowledge represents the first application of this technology for RNA barcode readout at clonal scale.

Importantly, X-CODE is also compatible with probe-based spatial transcriptomics. We demonstrate barcode detection using the 10x Genomics Xenium platform, establishing the feasibility of recovering clonal identity within spatial transcriptomic workflows while preserving tissue architecture. This compatibility is particularly notable given the finite probe capacity of current spatial transcriptomics platforms and the lack, to date, of barcode systems with demonstrated applicability to Xenium at the scale required for clonal analysis.

Application of X-CODE to an *in vivo* model of androgen deprivation in prostate cancer illustrates how this integrated approach can resolve clonal responses to therapy at high spatial and phenotypic resolution. Longitudinal analysis showed that castration resistance emerges after a period of clonal stability, consistent with heterogeneous adaptive potential among lineages and with previous studies^58^. Spatial mapping revealed that this transition is accompanied by a reorganisation of tumour architecture, with resistant tumours dominated by spatially segregated clonal expansions. Integration of clonal identity with multiplex phenotyping and spatial metabolomics further showed that dominant clones adopt distinct phenotypic and metabolic states, reflecting multiple, clone-specific modes of therapy-induced plasticity rather than a single convergent resistant phenotype, consistent with observations in patients^59^.

As a static barcoding strategy, X-CODE is designed to capture clonal identity rather than lineage ancestry and therefore does not provide information on division history or enable reconstruction of full lineage trees. Probe-based readouts also require allocation of detection channels to barcode units, which can limit the number of phenotypic markers measured in a single assay, particularly in cytometry-based applications. In addition, although X-CODE is broadly compatible across platforms, optimal co-detection of barcodes and phenotypic features may require platform-specific optimisation of reagents and workflows. Together, these considerations reflect design choices inherent to probe-based, multiplatform barcoding approaches and are balanced by the scalability and flexibility that X-CODE enables across diverse experimental contexts.

Together, these results establish X-CODE as a broadly compatible and flexible framework for unified clonal tracking across readily available platforms, enabling direct integration of lineage identity with phenotype, metabolism, and transcriptional state within intact tissue architecture, while preserving compatibility with barcode-guided clonal recovery for downstream functional studies.

## RESOURCE AVAILABILITY

### Lead Contact

Further information and requests for resources and reagents should be directed to and will be fulfilled by the lead contact, Marco Bezzi.

### Materials Availability

The X-CODE 15K and 2M libraries generated in this study have not yet been deposited in any repositories; however, they are available upon request.

## ACKNOWLEDGEMENTS

We thank the Confocal Microscopy and Pathology facilities at the Institute of Cancer Research for their support, and the UK Dementia Research Institute at Imperial College London for accommodating mass cytometry experiments, with special thanks to Dorcas Cheung and Vicky Chau. This work was supported by Cancer Research UK (RCCFEL/100053 Career Establishment Award to M.B.; ICR PhD fellowship to R.R.; Convergence Science Centre PhD fellowship to E.F.; CRUK Commercial Partnerships Project Development Fund to M.B.) and Prostate Cancer UK (RIA21-EOI-020 Research Innovation Award to M.B.). Several schematic figures were created using BioRender.

## AUTHOR CONTRIBUTIONS

Conceptualisation, M.B. (Bezzi); Funding acquisition, M.B. (Bezzi), D.R., L.M., Z.T.; Methodology, E.F., R.R.; Investigation, E.F., R.R., M.T., A.D., A.T., N.P., F.G., A.W., I.A., K.L.C., C.G.; Analysis: M.B. (Bezzi), E.F., R.R., M.B. (Bourdim), E.W., M.S., J.S.M.; Software: M.B. (Bourdim), E.W., J.S.M.; Writing – original draft: M.B. (Bezzi), R.R., E.F., M.T., M.B. (Bourdim); Writing – review & editing: M.B. (Bezzi), R.R., E.F., M.T., E.W., M.S., J.S.M., A.D.; Resources: Z.T., A.D., L.G., C.G.

## DECLARATION OF INTEREST

The authors declare no competing interests.

## DECLARATION OF AI-ASSISTED WRITING

ChatGPT (OpenAI) was employed to support grammar correction and clarity improvements during manuscript drafting. All sections generated with assistance were critically reviewed and edited by the authors, who take full responsibility for the final content.

## METHODS

### Cell lines

TRAMP-C1 (ATCC; #CRL-2730), RM-1 (ATCC; #CRL-3310) and MyC-CaP (ATCC; #CRL-3255) cell lines are commercially available from ATCC. HEK-293T cell line was generously provided by the Preclinical Modelling of Paediatric Cancer Evolution team at the Institute of Cancer Research (ICR). TRAMP-C1 were grown in Dulbecco’s Modified Eagle’s Medium (DMEM; Thermo Fisher; #10313021) supplemented with 10% Fetal Bovine Serum (FBS; Sigma-Aldrich; #F9665), 4 mM L-glutamine (Thermo Fisher; #25030081), 1x Penicillin–Streptomycin (Thermo Fisher; #15140122), 0.005 mg/ml bovine insulin (Scientific Laboratory Supplies; #LZBE02-033E20) and 10 nM dehydroisoandrosterone (Sigma-Aldrich; #D4000). HEK-293T, RM-1 and MyC-CaP were grown in DMEM supplemented with 10% FBS, 4 mM L-glutamine and 1x Penicillin–Streptomycin. All cell lines were tested negative for mycoplasma using the Mycoplasma PCR detection kit (Abcam; #ab289834).

### Animal work

All animal experiments were approved and monitored by the Institute of Cancer Research Animal Welfare and Ethical Review Body (protocol PP9736772), in compliance with the UK Home Office Animals (Scientific Procedures) Act 1986, the United Kingdom National Cancer Research Institute guidelines for the welfare of animals in cancer research, and the ARRIVE guidelines. Mice were kept under standard conditions with ad libitum food and water. Tumour growth was measured twice weekly with callipers, and volume was calculated as (length × width²) / 2. Mice were monitored regularly and euthanised by CO₂ asphyxiation followed by cervical dislocation when tumour volume reached approximately 850 mm² or when any single tumour dimension exceeded 15 mm (endpoint).

#### RM-1 *in vivo* experiments

1 × 10⁶ W, Z, Y, J (1:1:1:1) RM-1 or 1 × 10⁶ RM-1-15K were injected to generate tumours used for spatial analyses and MALDI. Cells were detached and filtered to obtain a single-cell suspension, mixed in 200 µL 1:1 PBS and Matrigel (Corning; #356231) and injected subcutaneously into a single depilated and sterilised flank of C57BL/6J recipient mice (RRID: IMSR_JAX:000664). Mice were culled at experimental endpoint and tumours were collected, snap-frozen in isopentane and stored at −80 °C for subsequent analyses.

#### MyC-CaP *in vivo* experiments

30 FVB-NJ mice (RRID: IMSR_JAX:001800) 6-8 weeks old were subcutaneously injected with 1 × 10⁶ barcoded MyC-CaP cells in 200 µL of 1:1 PBS and Matrigel solution. All experimental groups excluding the uncastrated and pre-castration control groups received chemical castration via Degarelix (0.315 mg, 100 µL, every 14 days) after 21 days or when tumours exceeded 400 mm². Control groups included eight uncastrated mice sacrificed at endpoint and five mice sacrificed pre-castration. Post-castration groups were collected at early (4 days, n=5), late (10 days, n=6), and endpoint (n=6) timepoint. Mice were monitored and culled in accordance with the limits described above. After collection, tumours were sectioned into four parts: two central sections were snap-frozen, and two outer sections were used for DNA extraction and primary cell culture.

### X-CODE lines and libraries generation

#### Oligo Probes

All oligo probes reported in Supplementary Tables (Table 6 – Probe seq; Table 1 – Gibson Assembly; Table 4 – Add-ons Primers) were purchased by IDT as 25 nmol DNA Oligo, 100 nmol DNA Oligo or 4 nmole Ultramer™ DNA Oligo depending on probe size and modifications following IDT instructions.

**Table 1.**
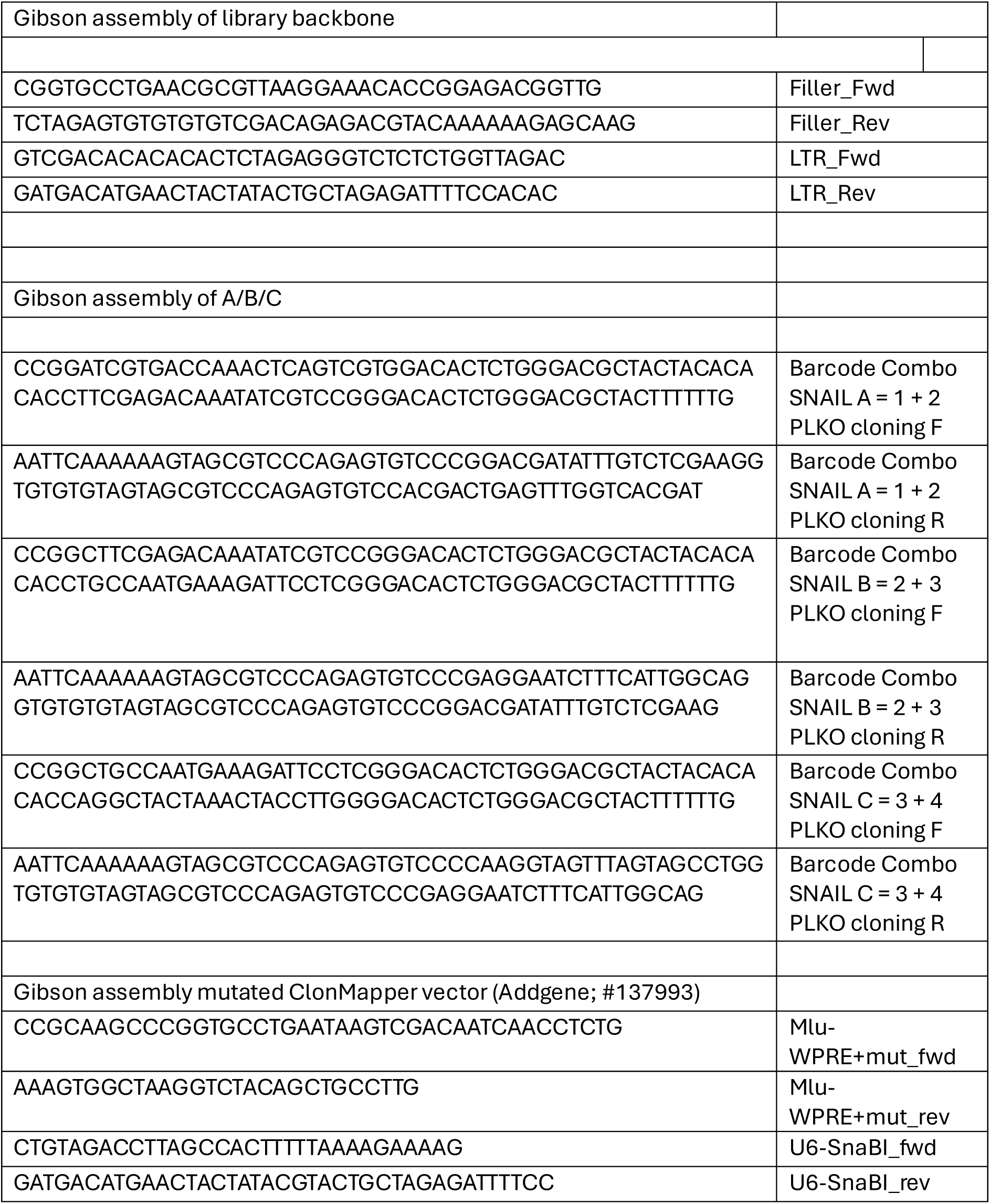
– Gibson Assembly.

#### Assembly of barcode A/B/C vectors

Primers for Gibson assembly were designed using the NEBuilder Assembly Tool (https://nebuilder.neb.com/#!/). Four sequences were divided into three pairs (Table 4 - Gibson assembly). CROPseq-Guide-puro plasmid (Addgene; #86708) was digested with MluI-HF (NEB; #R3198S) and gel-purified. The oligos were annealed, PCR-amplified using Q5 High-Fidelity PCR (NEB; #E0555L) and gel purified. Gibson assembly was performed using NEBuilder HiFi DNA Assembly Kit (NEB; #E5520S) with a 1:5 vector-to-insert ratio and extended incubation at 50 °C for 60 min. The resulting plasmid was used to transform chemically competent cells (Thermo Fisher; # C737303).

#### Chemical transformation

Chemically competent cells were thawed on ice and transformed with either 5 μL of the assembly product or ligated plasmid without the insert (negative control). After mixing, samples were incubated on ice for 30 minutes, heat-shocked at 42°C for 30 seconds, chilled for 2 minutes, and recovered in 950 μL SOC at 37°C for 1 hour with shaking. Cultures were plated on agar plates containing Ampicillin (100 μg/mL) and incubated overnight at 37°C. Individual colonies from the assembly plate were inoculated into Ampicillin-supplemented LB and grown for 24 hours at 37°C with shaking. Plasmids were extracted using the QIAprep Spin Miniprep Kit (QIAGEN; #27106) following the manufacturer’s instruction. DNA concentration was measured, and clones were validated by Sanger sequencing.

#### Electroporation

Electroporation was performed using the GENE-Pulser electroporator (Bio-Rad #165278) with the Gene Pulser Controller (Bio-Rad #165-2098) at 2.5 kV, 25 μF, 200 Ω. Electrocompetent *E. coli* (Agilent; #200227) and 2 mm cuvettes were kept on ice. For the 1st pooled ligation, 50 μL cells + 5 μL plasmid were used; for the 2nd/3rd pooled ligations, 100 μL cells + 10 μL plasmid. A negative control used 50 μL cells + 5 μL backbone (no insert). After electroporation, cells were recovered in SOC medium (950 μL or 1900 μL depending on cell volume) and incubated at 37 °C for 30 min, 250 rpm. To calculate the efficiency, the transformed cells were plated at different dilutions in agar plates containing Ampicillin (100 μg/mL) with controls plated at the highest dilution and grown overnight at 37 °C upside down. The remaining cells were grown in 450 mL LB-broth supplemented with 100 μg/mL Ampicillin and incubated overnight at 37 °C shaking. After extraction, plasmids were validated by Sanger sequencing.

#### Assembly of library backbone (CROPseq-puro-filler-LTR-MluI-HF/SalI-HF)

The primers for Gibson assembly were designed using the NEBuilder Assembly Tool. The Filler_Rev and the LTR_Frwd primers (Table 1 - Gibson assembly) were designed to harbour within the spacer regions, two new restriction sites for SalI and XbaI split by a linker region. 1 μg of CROPseq-Guide-puro (Addgene; #86708) was digested with SnaBI (NEB; #R0130S) and SalI-HF (NEB; #R3138S) and gel purified. The filler and the LTR fragments were generated through PCR from the CROPseq-Guide-puro plasmid (Addgene; #86708) and gel purified. The Gibson assembly was performed according to the NEBuilder® HiFi DNA Assembly Cloning Kit instructions in a 1:5 ratio. To increase the efficiency of the reaction the incubation at 50°C was extended from 15 to 60 minutes. The ligation product was used to transform chemically competent cells.

#### Barcode oligos design (X1, X2 and X3 sets)

The 27 variable binding sequences (VBS) (Table 2; 27_BUs) were based on the insert probes of the PLAYR technology, which have been shown to be compatible with probe-based detection strategies^43^. Of these, 23 VBSs were directly adapted from the PLAYR system, whereas 4 were generated using the same design principles (GC content; melting temperature; length). The universal binding sequence (UBS) was designed by combining the two backbone-binding regions from the same paper. Each barcode unit (BU) is 41 bp long, with the VBS (20bp) and UBS (20bp) separated by a single guanine nucleotide (G) that provides spacing to enable concurrent binding of the SNAIL and universal probes.

**Table 2.**
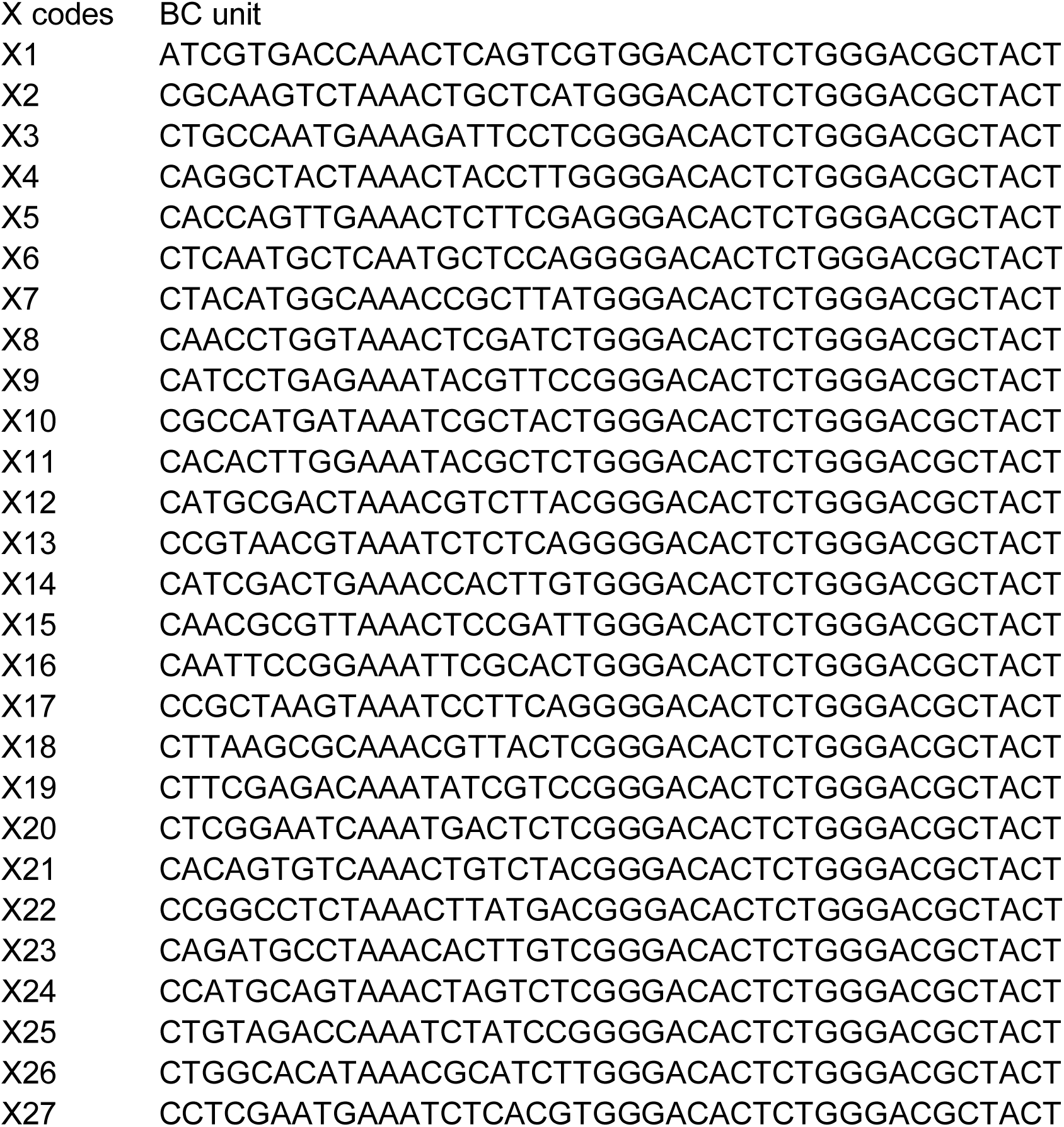
– 27_BUs.

The 127 unique oligos of each X set were generated with a “9 choose 4” combinatorial strategy. All the assembled sequences were appended with a Universal Binding Site (UBS) sequence. A linker region (ACACACAC) was inserted between the barcode sequences to facilitate the assembly of the SNAIL complex. A secondary barcode was constructed from three subsets of 4 base sequences, combined using a “4 choose 4” combinatorial design while excluding homo-tetrameric and homo-trimeric motifs. Each secondary barcode set was placed within a specific X-set, positioned downstream of the X-CODE sequences and separated from them by a BamHI–linker–EcoRI cassette. To integrate the oligos into the vector backbone, distinct restriction sites were added depending on the corresponding X-set. X1 oligos carried MluI (5′) and SalI (3′) sites, with a BbsI–spacer–cut site motif before SalI for scaffold RNA cloning downstream of Bc1. A BamHI–linker–EcoRI cassette between X1 and Bc1 enabled the three-step clonal strategy to assemble the final barcode. Both X2 and X3 oligos shared the same design, with an MfeI site at the 5′ end (compatible with BamHI after digestion) and a BglII site at the 3′ end (compatible with EcoRI). A total of 378 oligos were synthesised by IDT as MiniGene™ constructs in the pUCIDT (Amp) vector (Table 3 - Ordered_oligo_378).

**Table 3.**
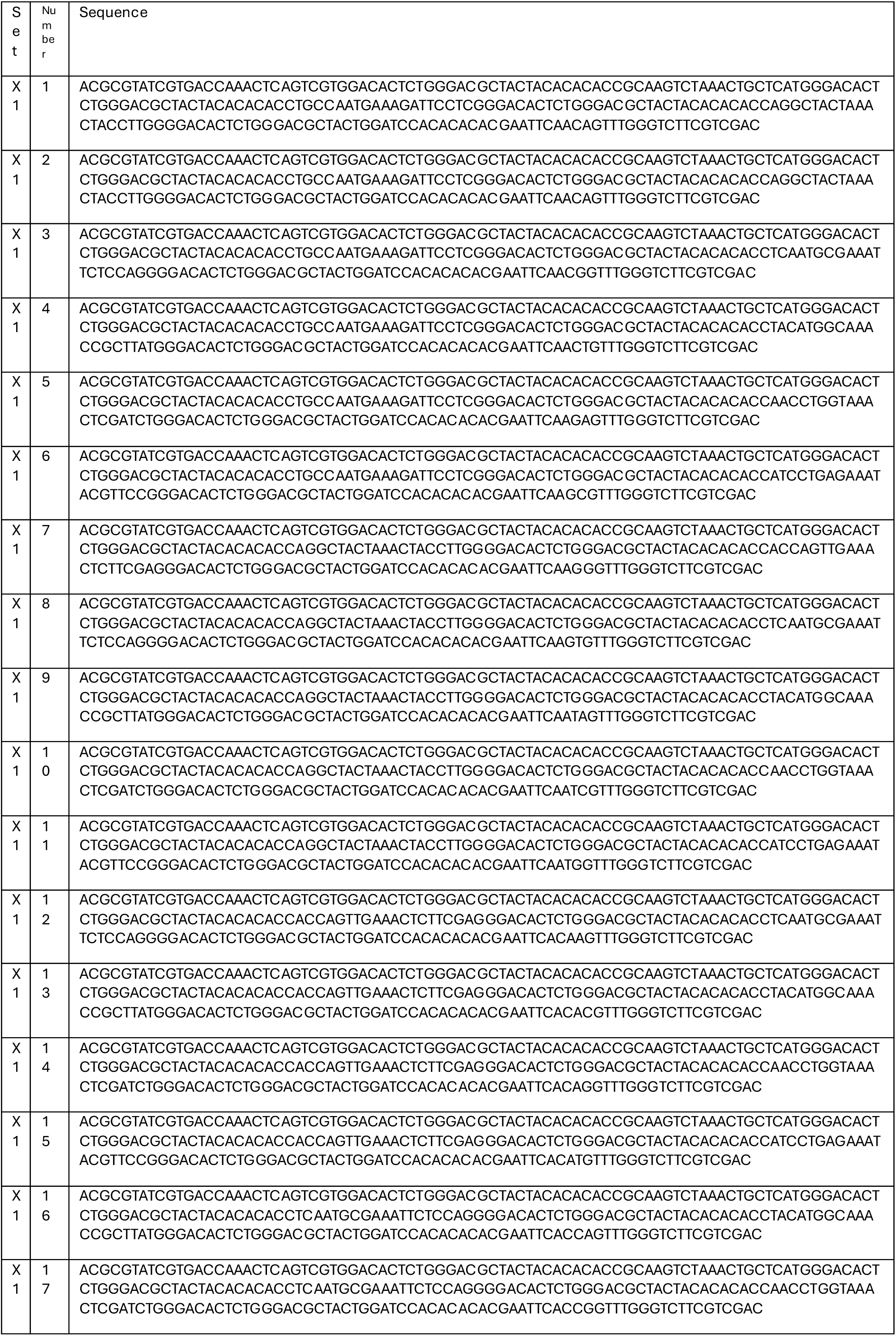
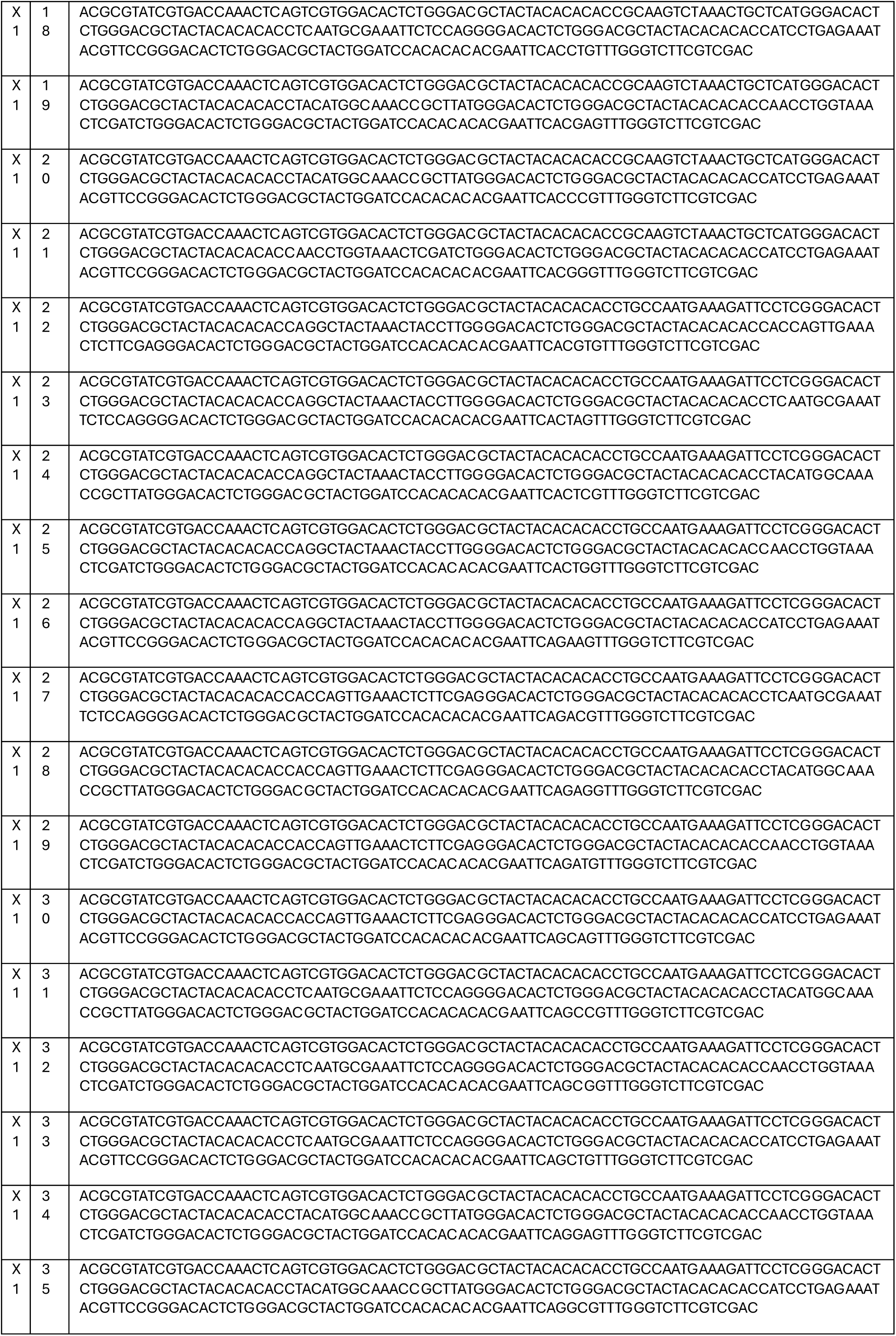
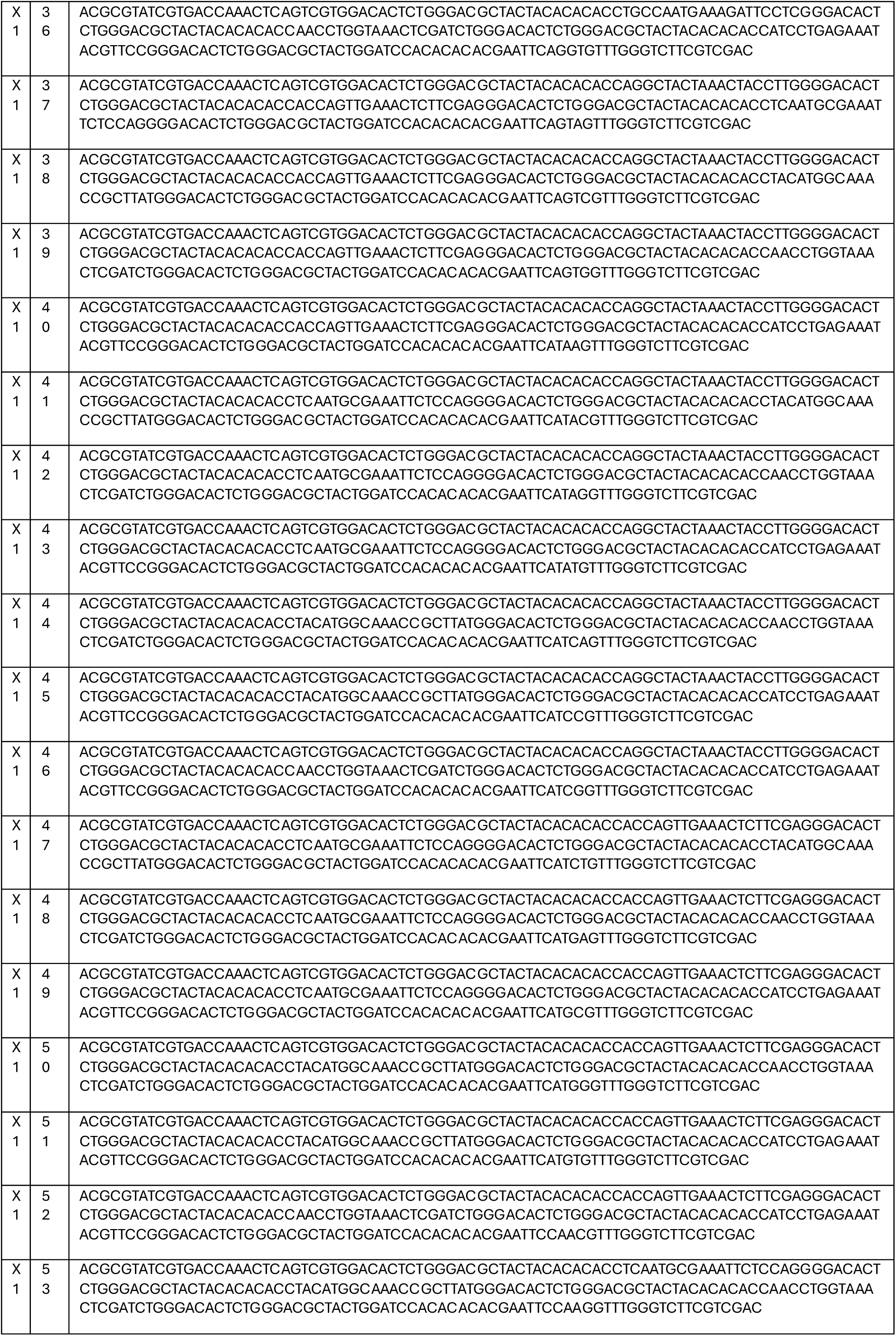
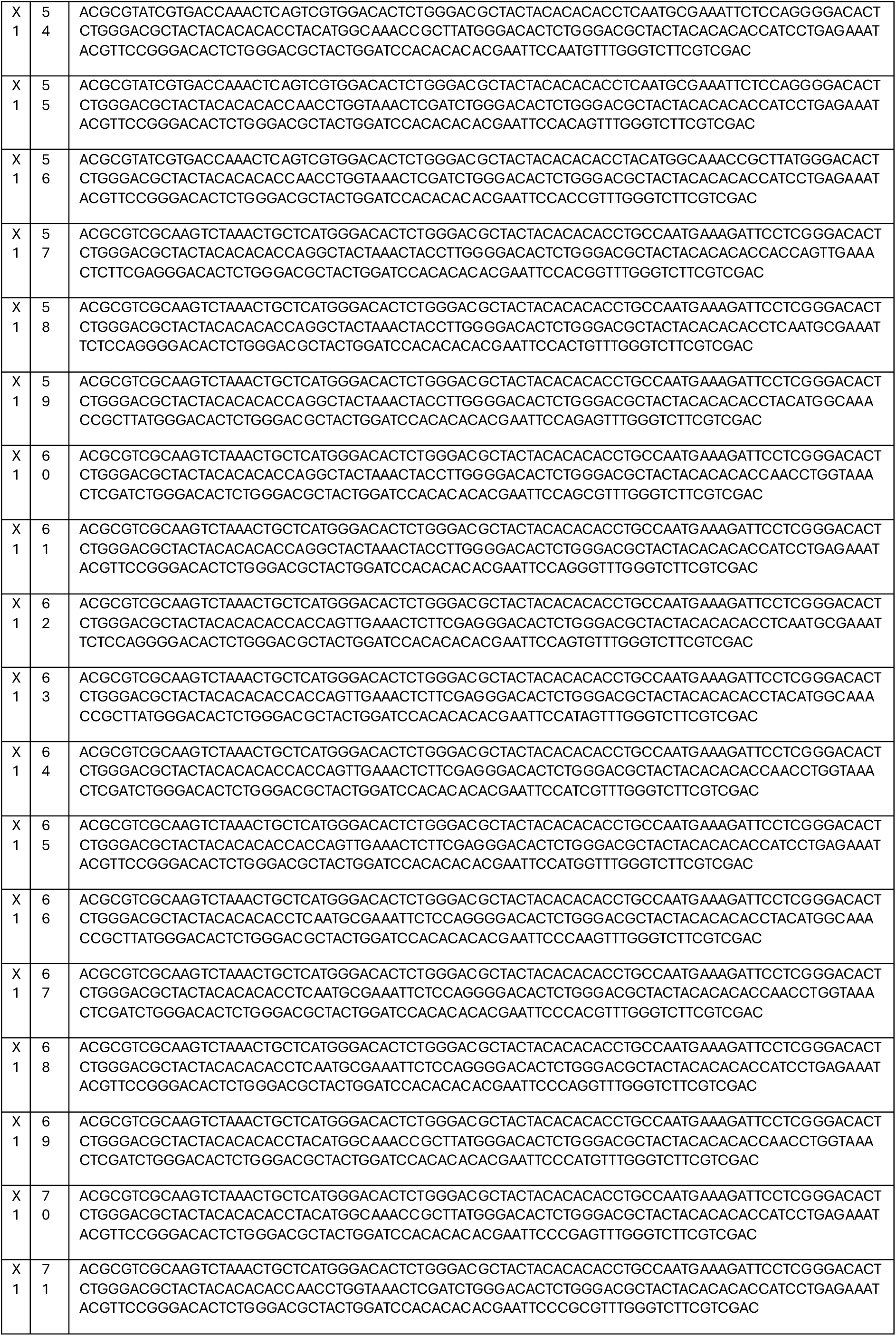
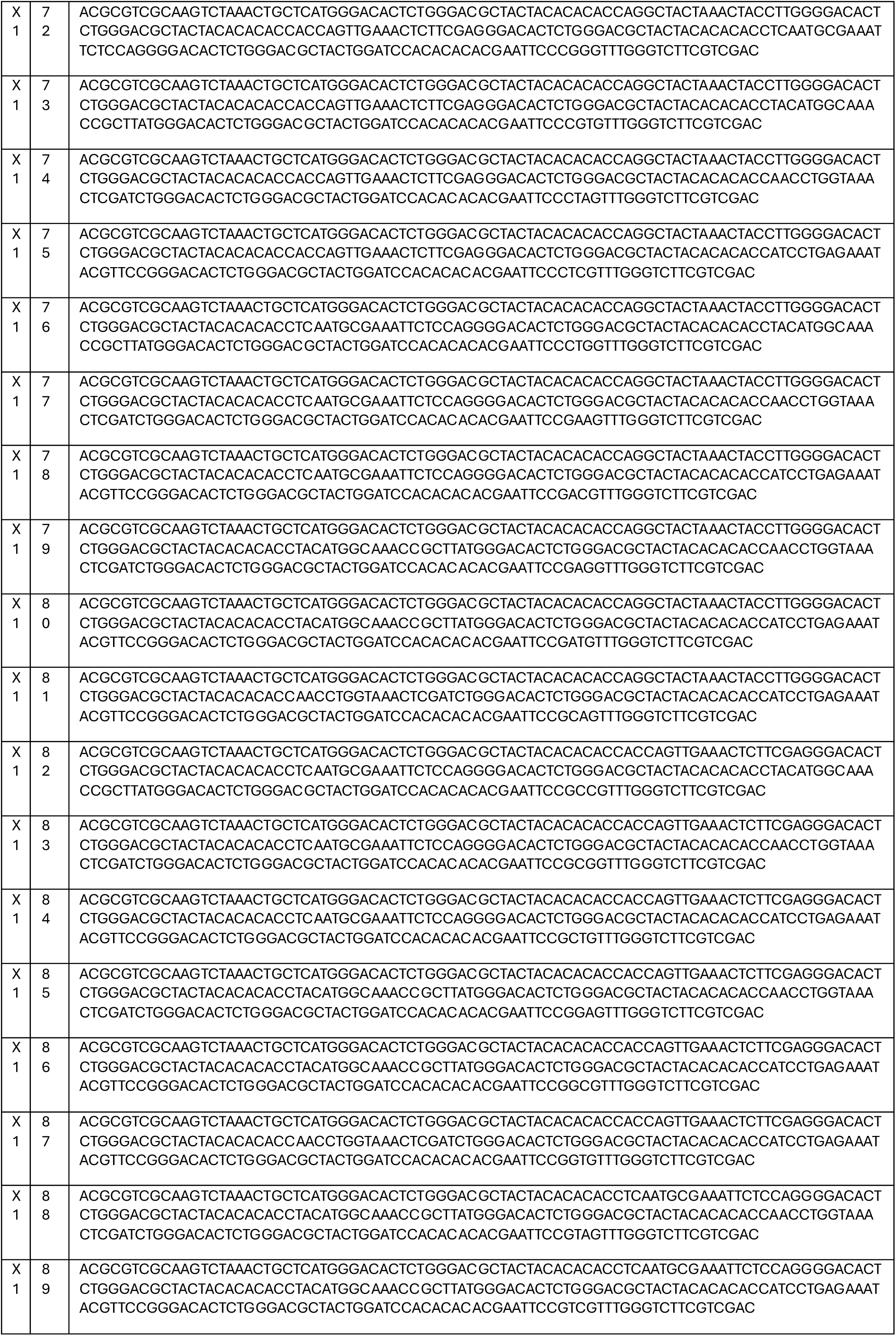
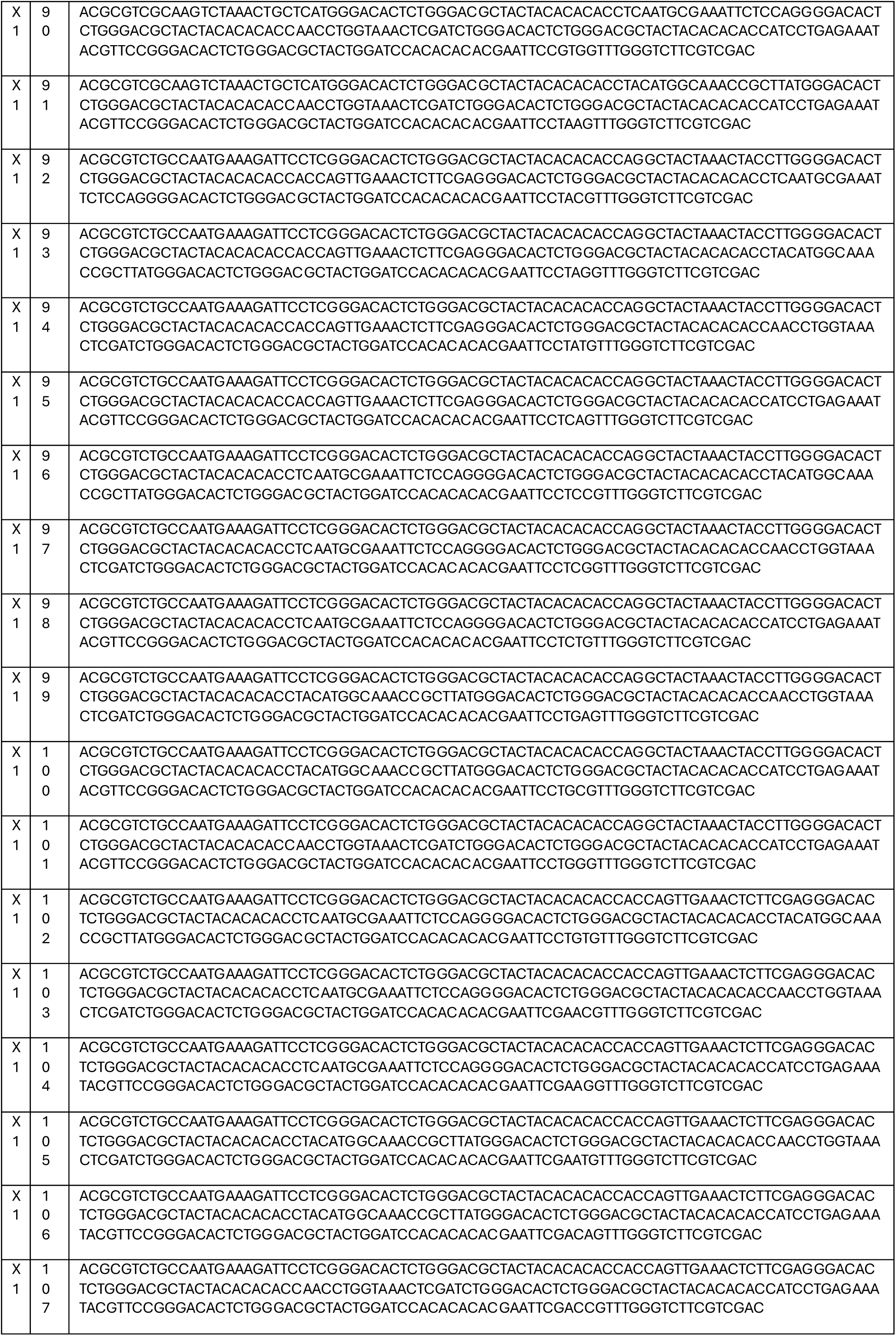
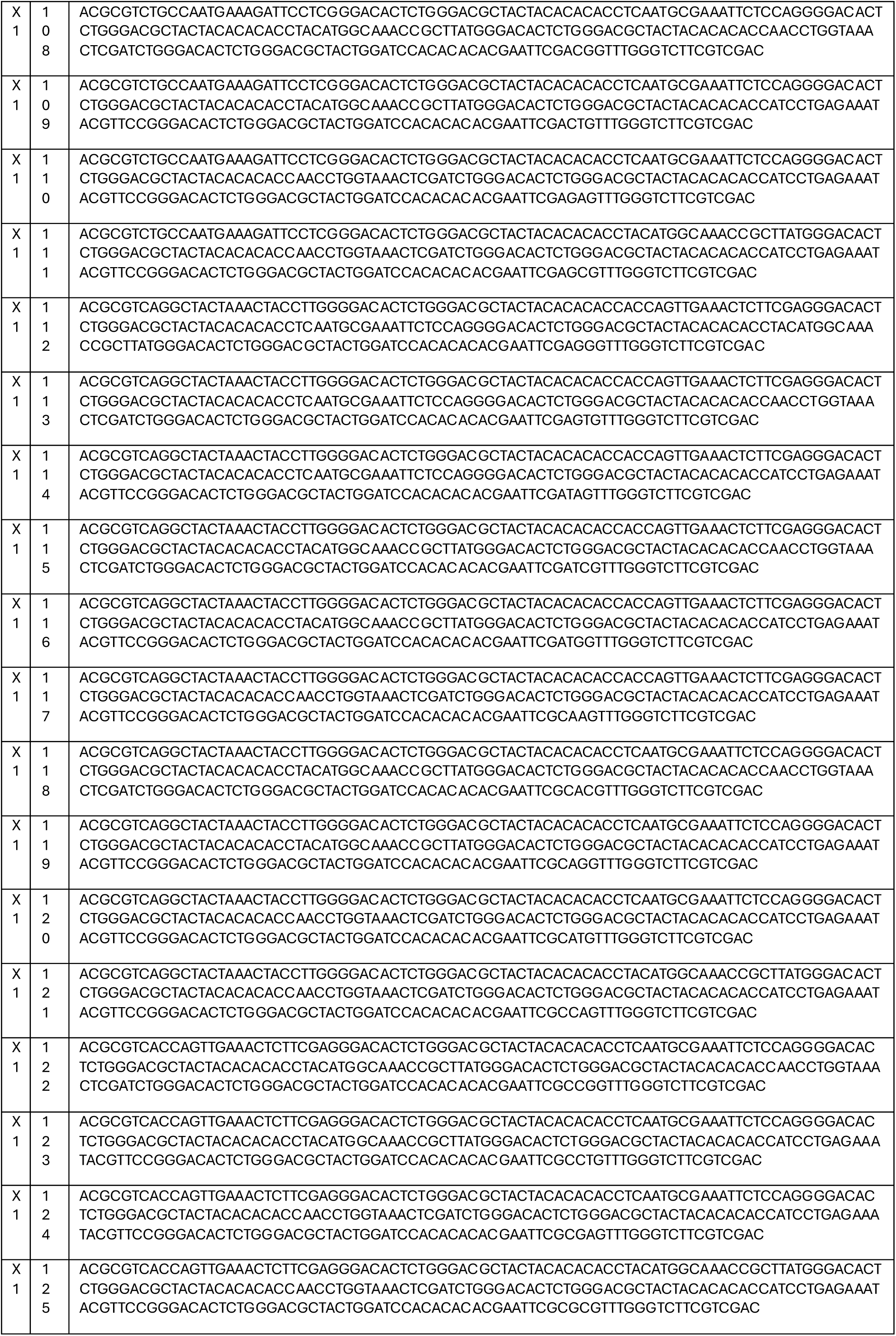
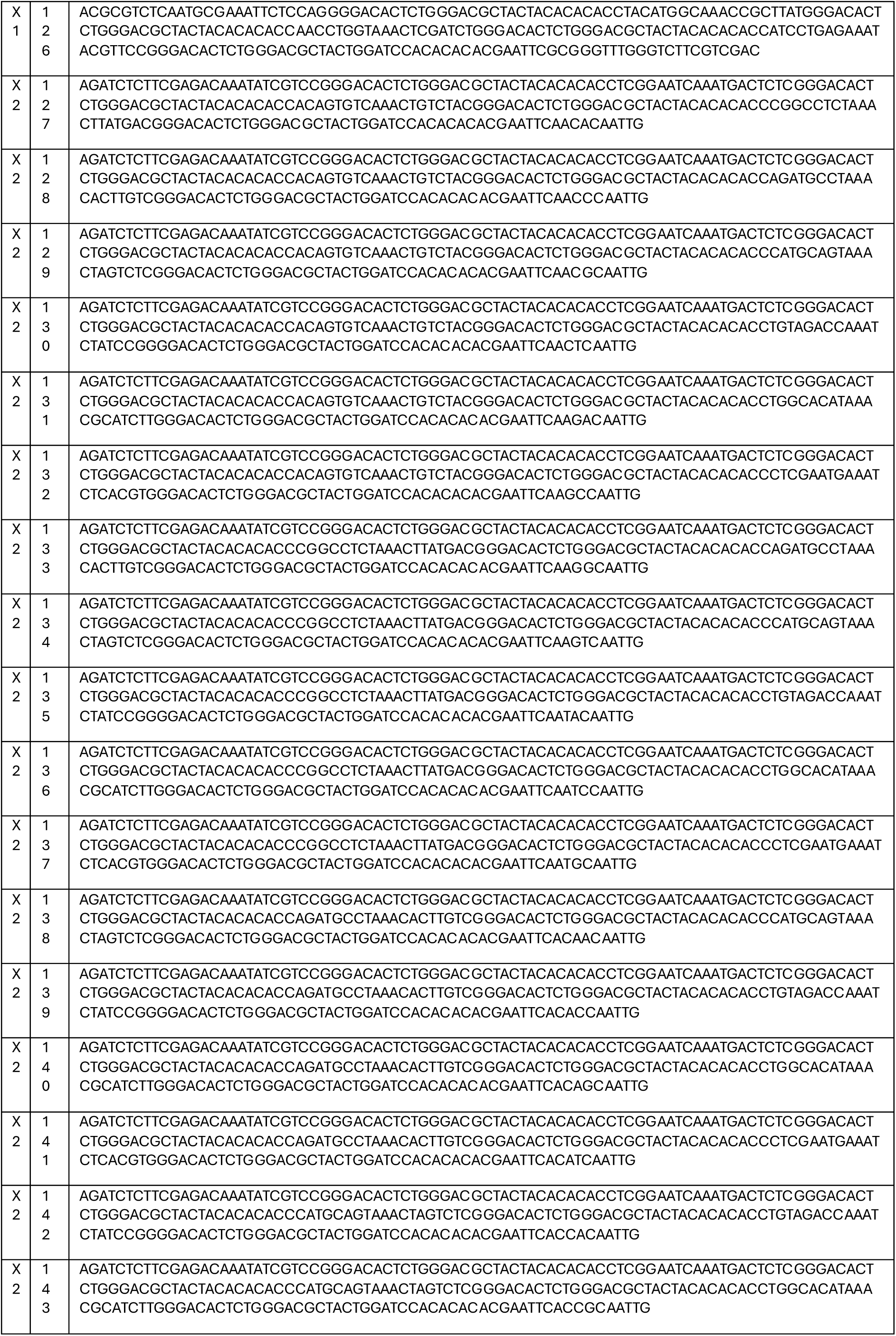
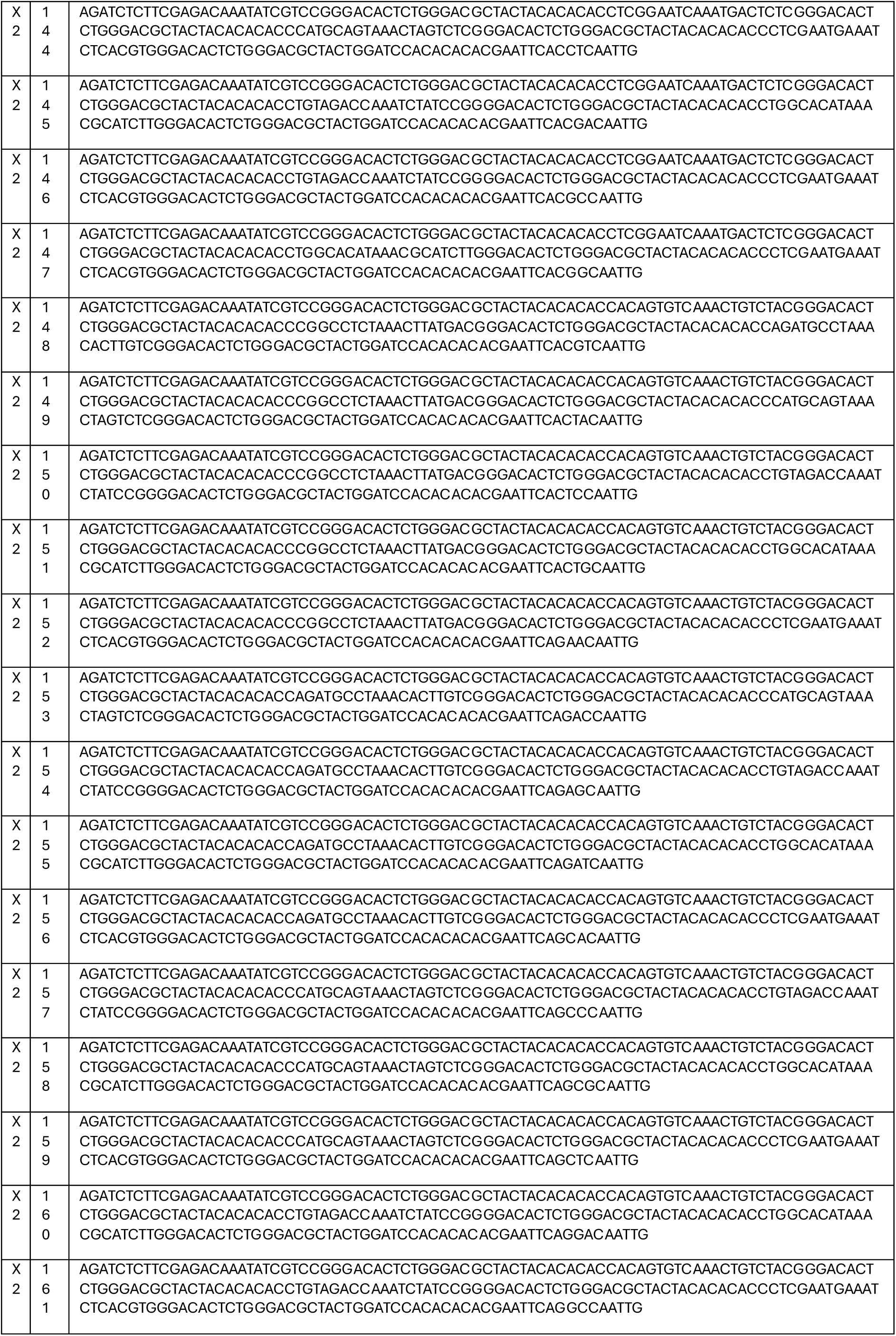
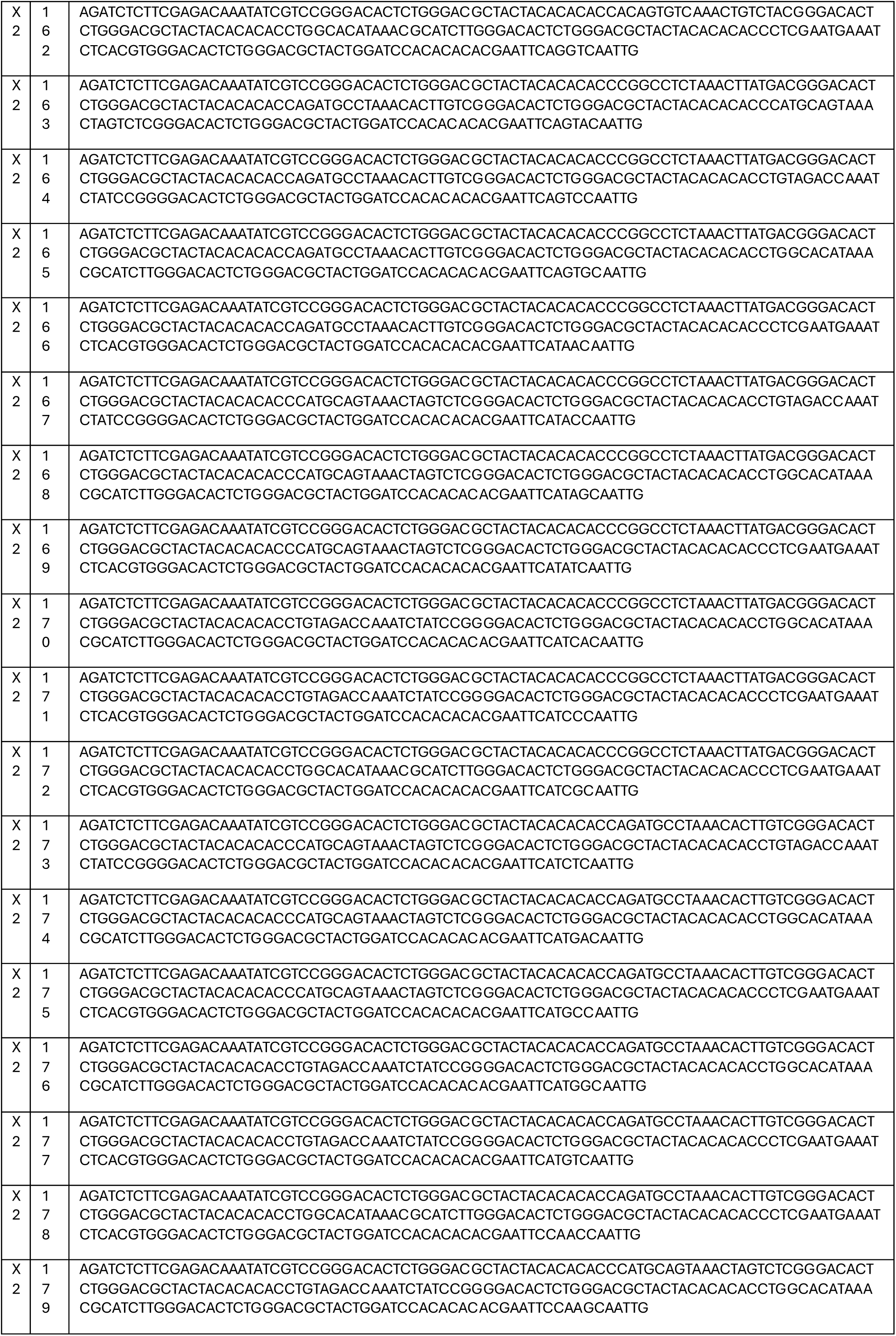
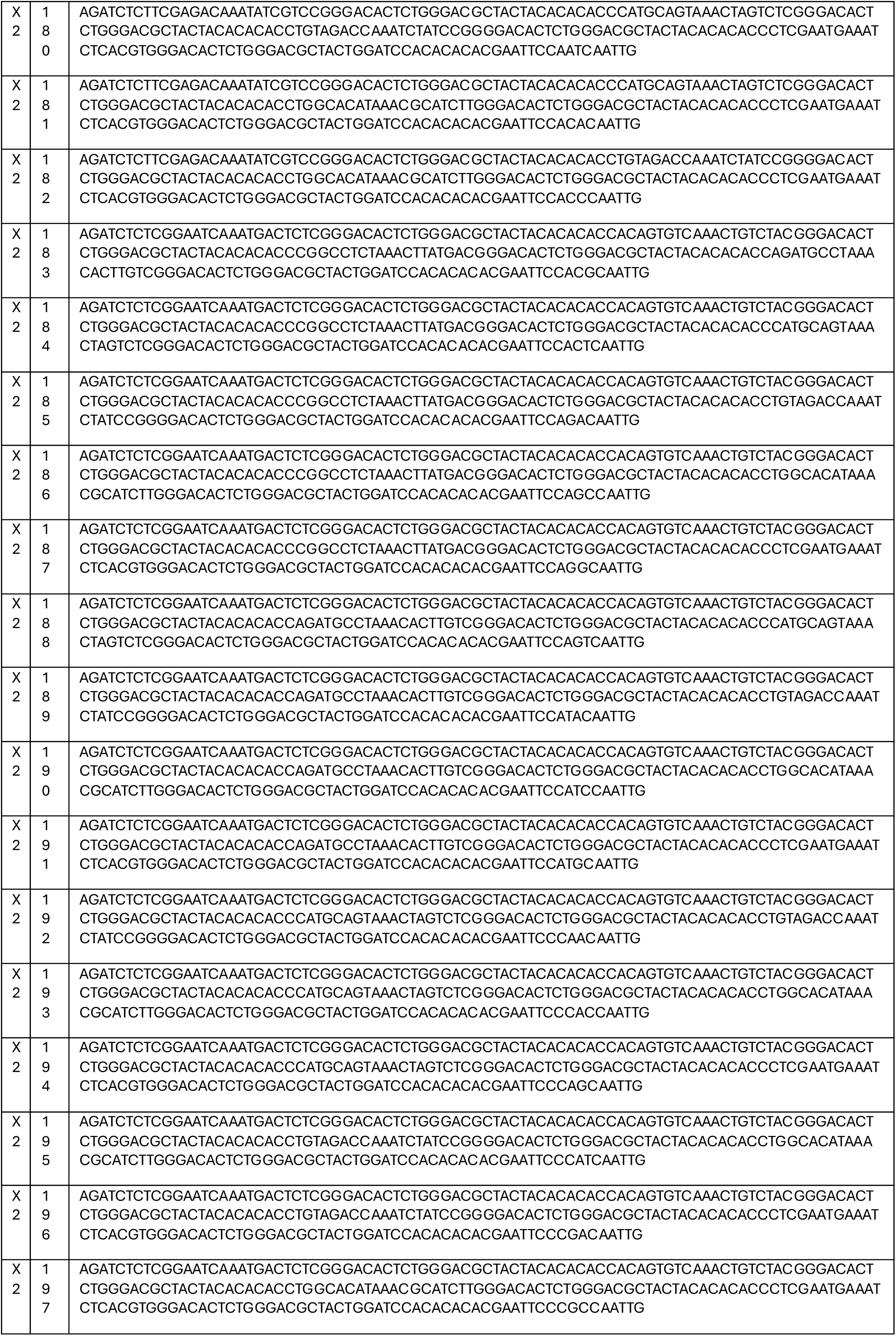
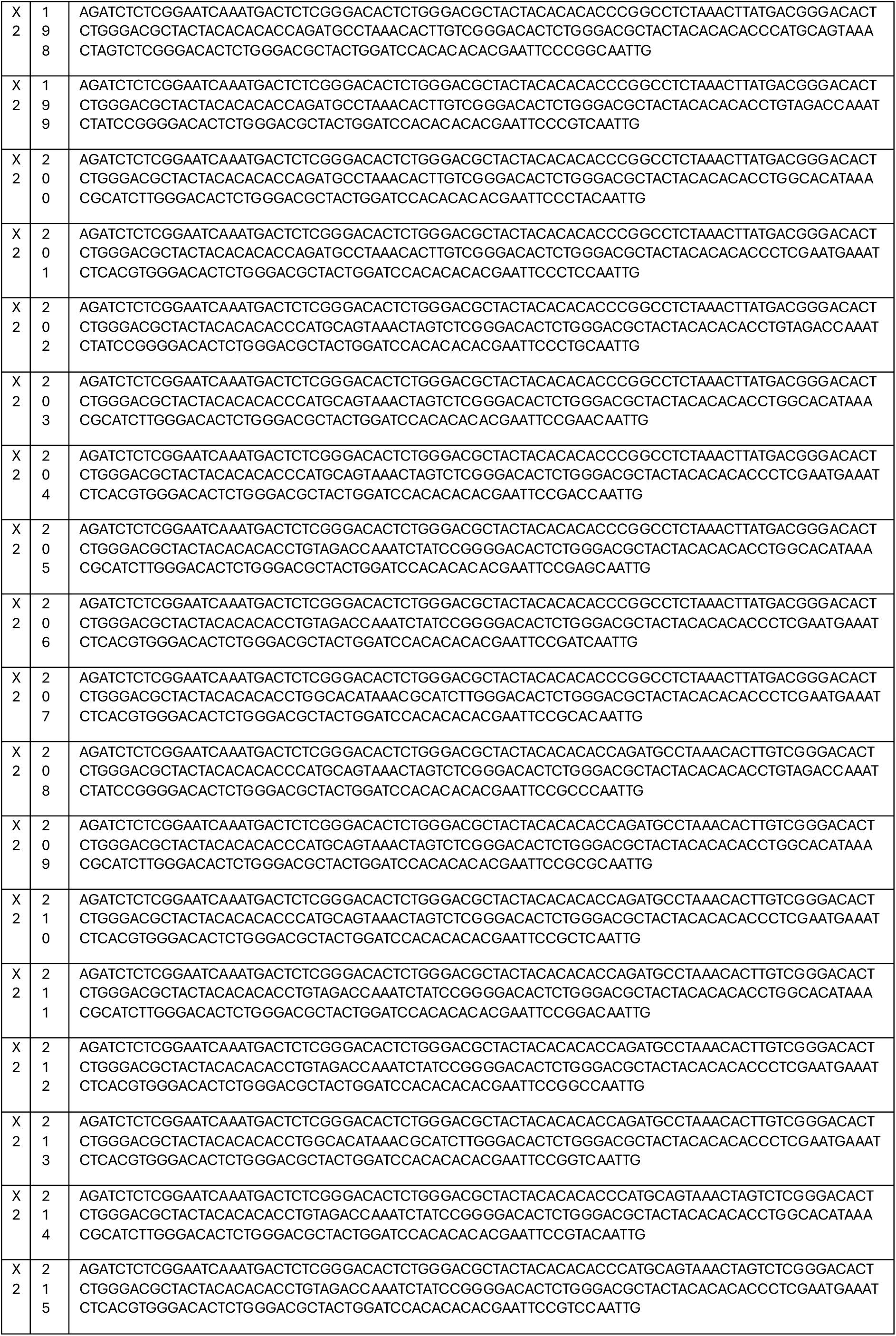
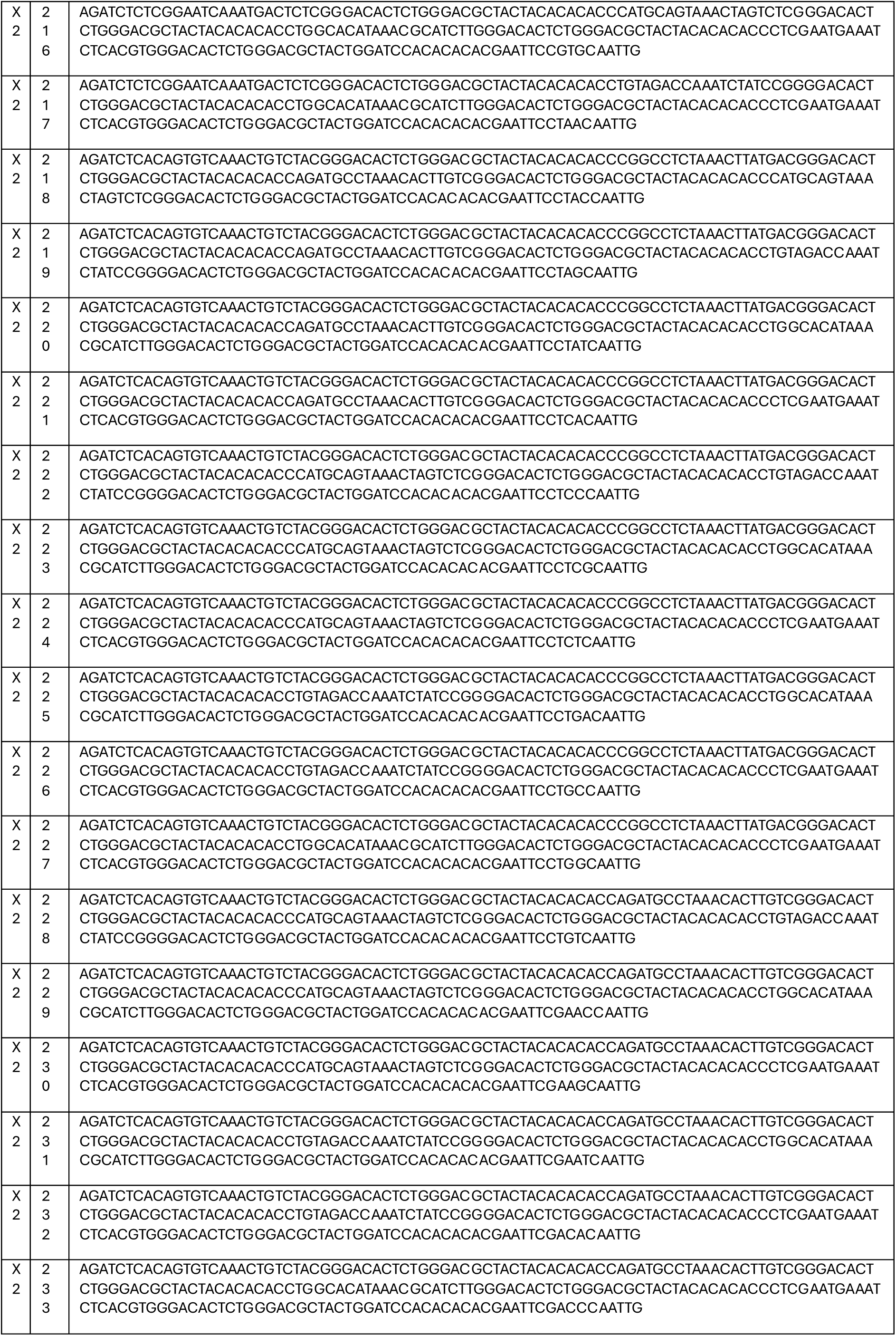
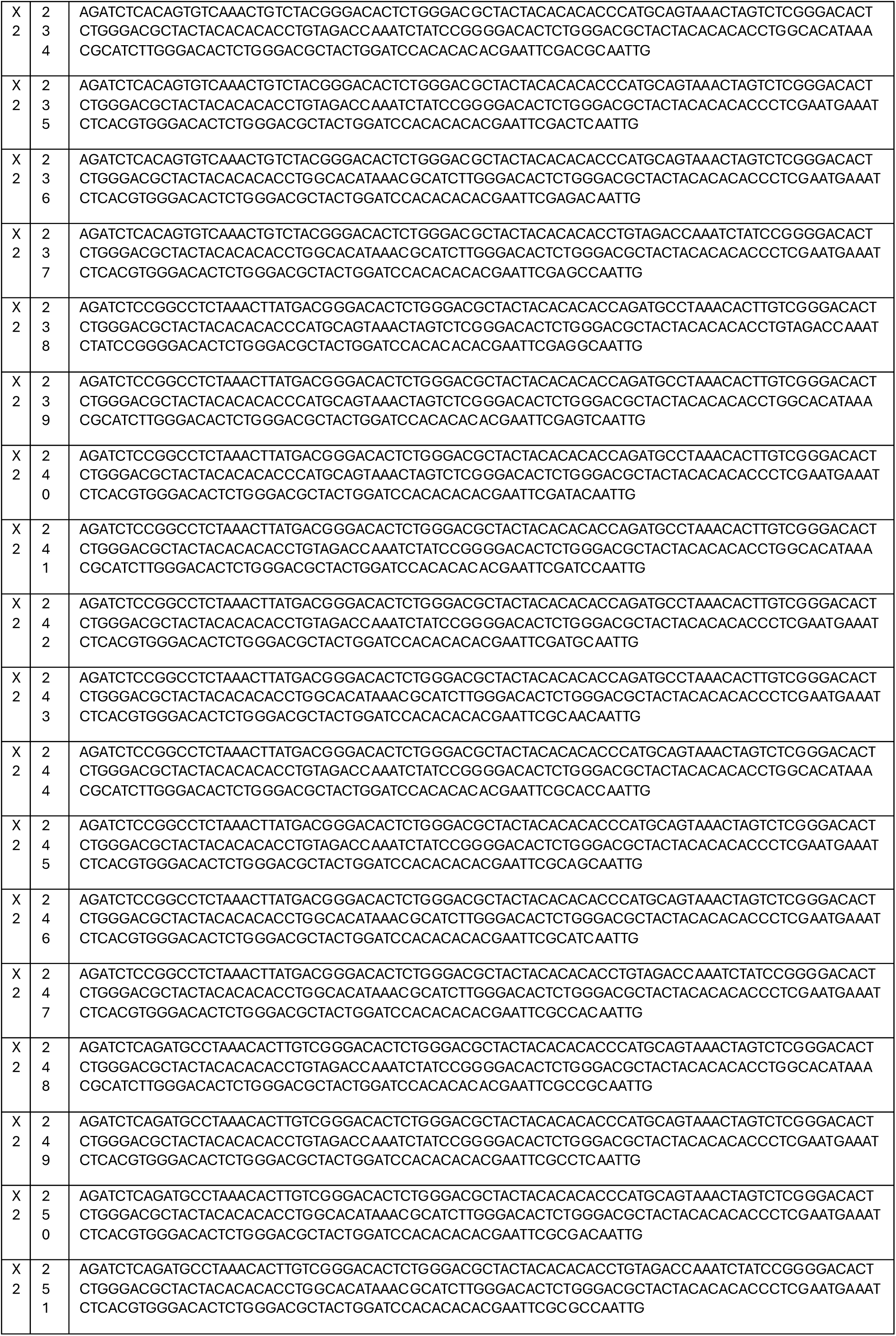
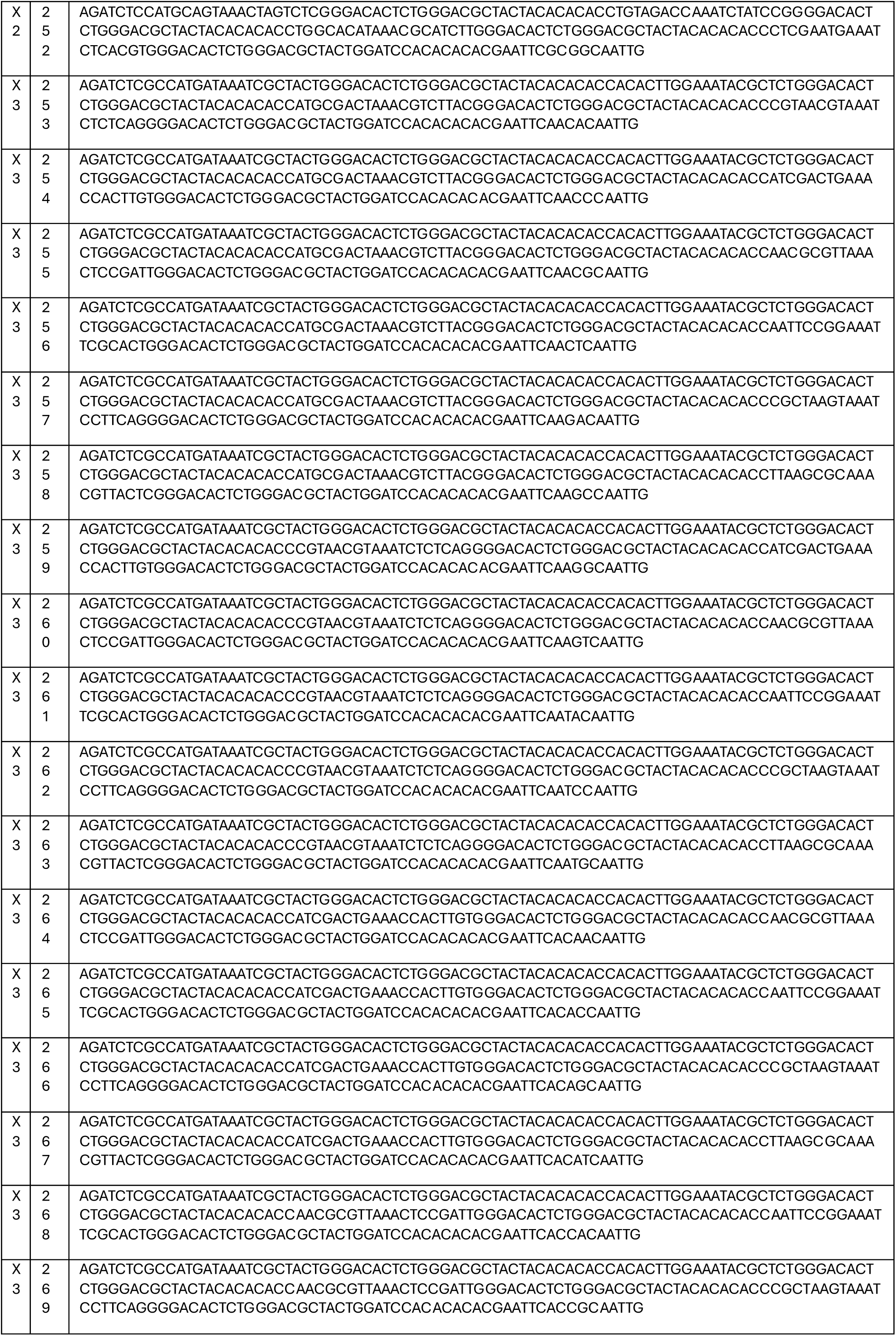
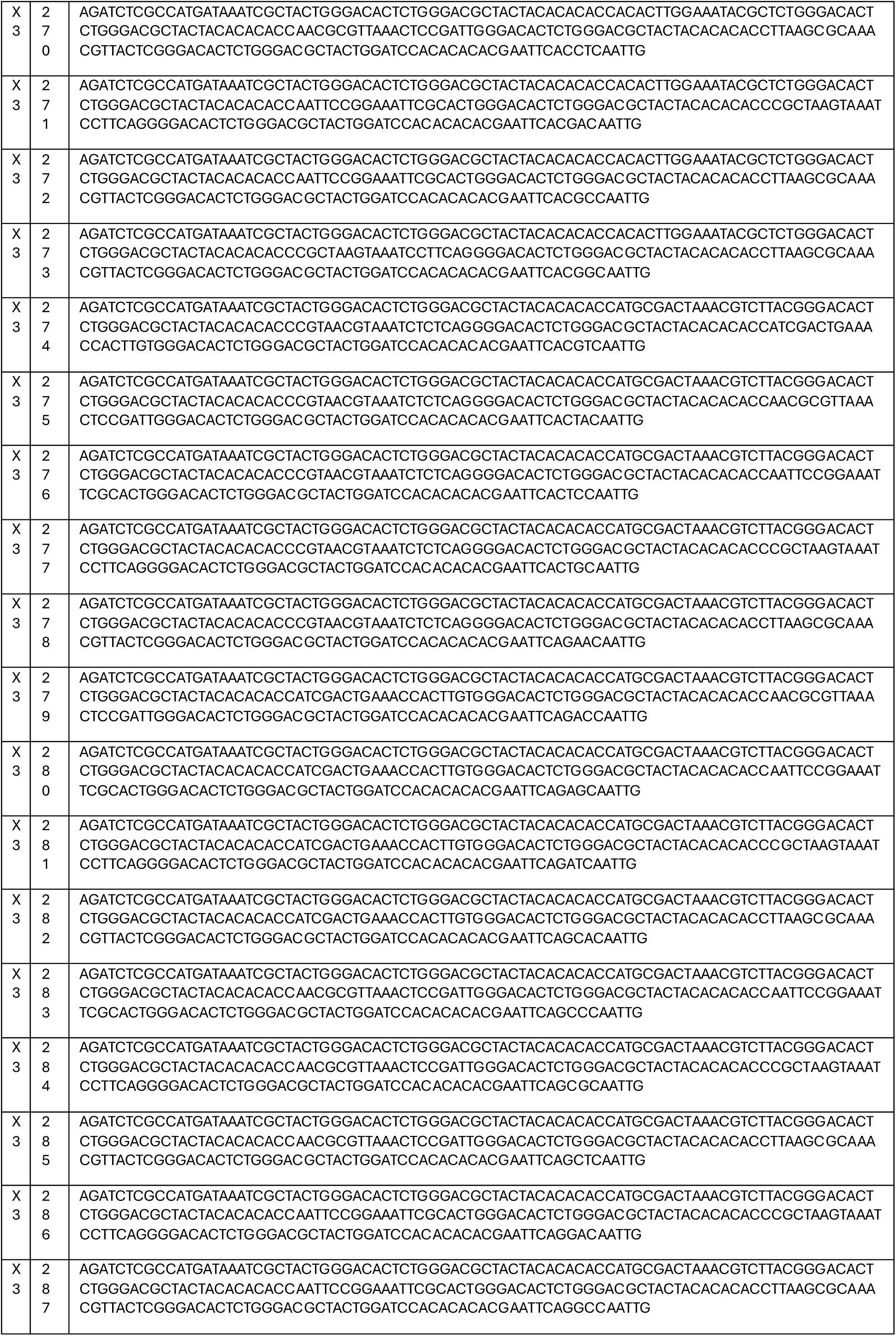
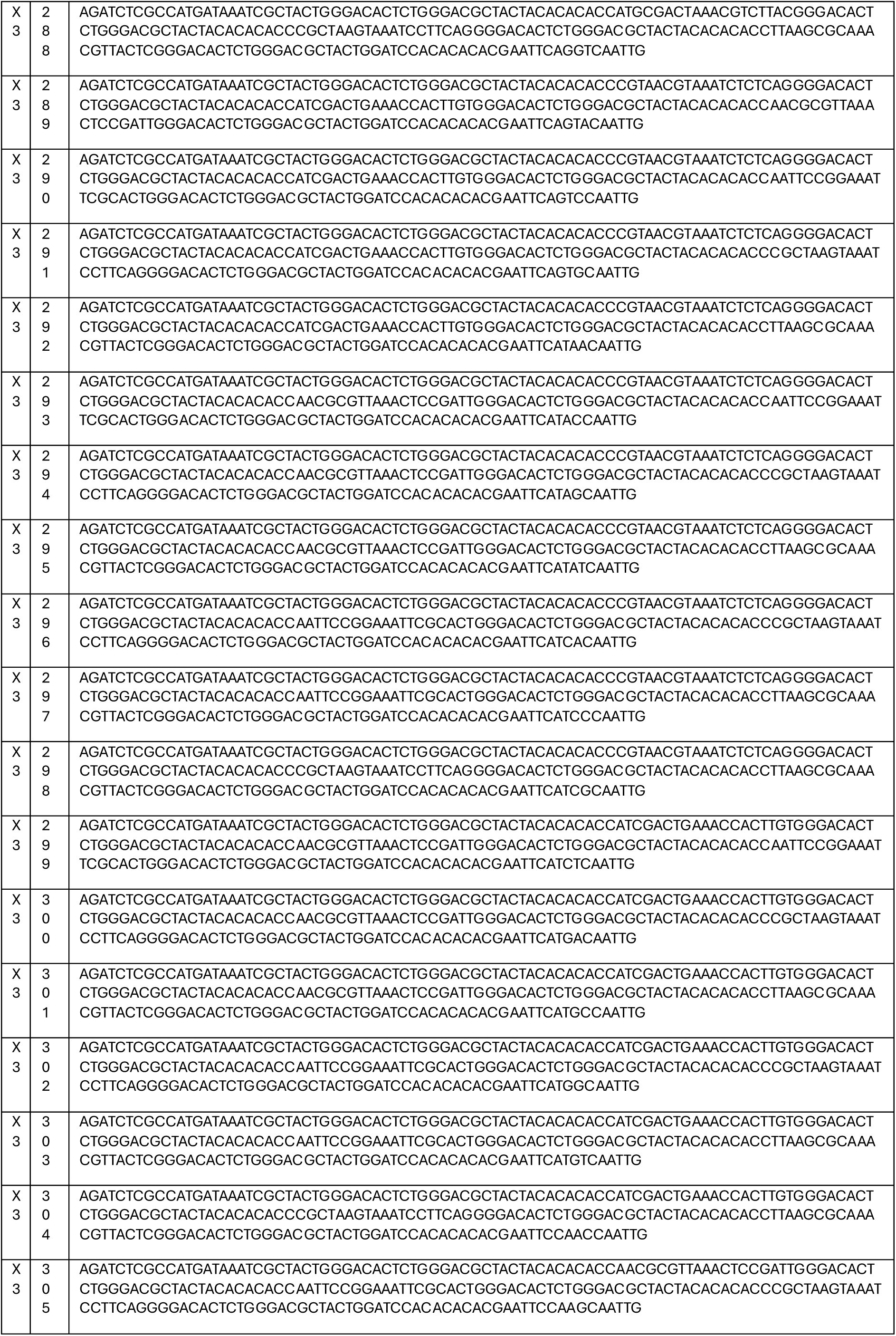
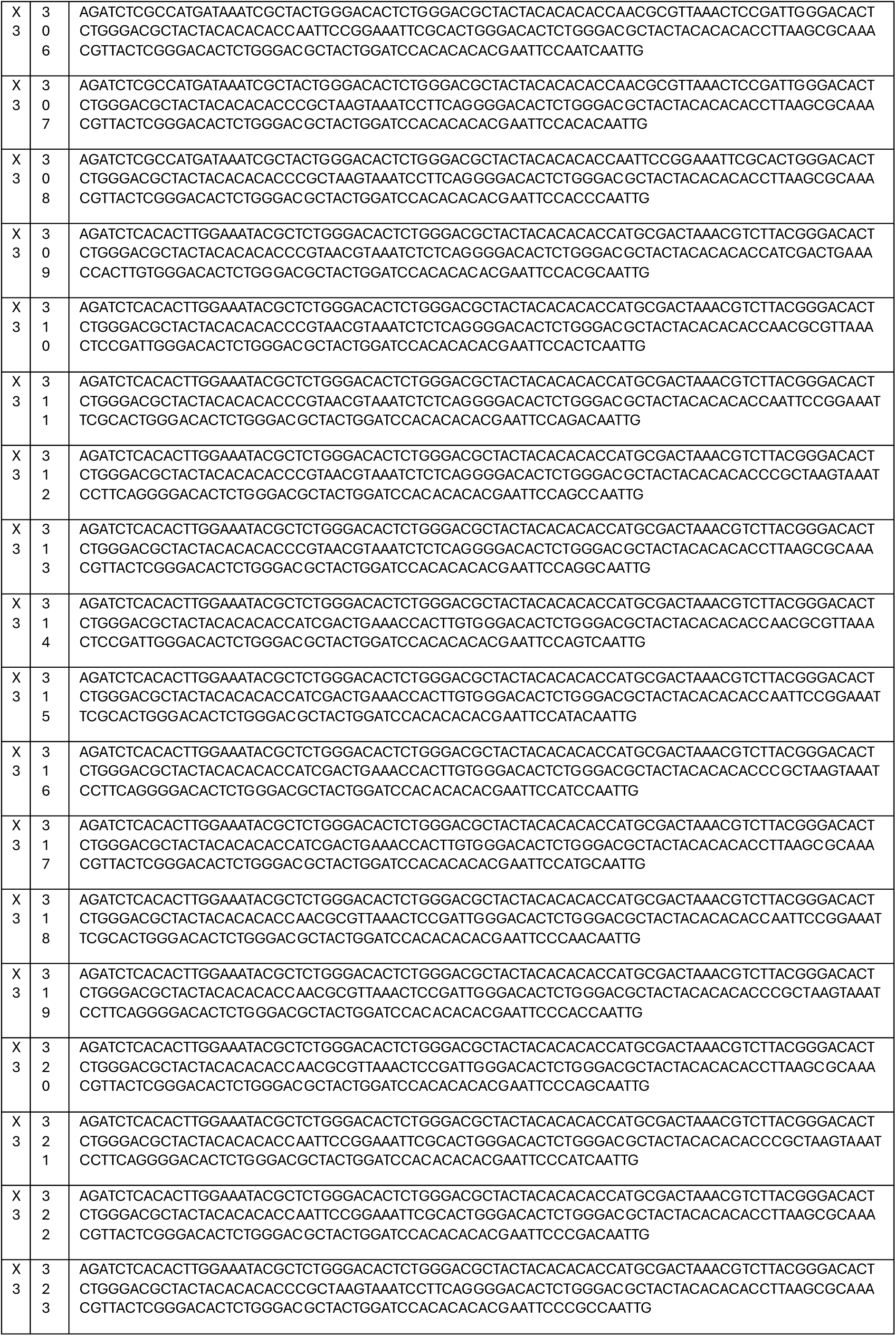
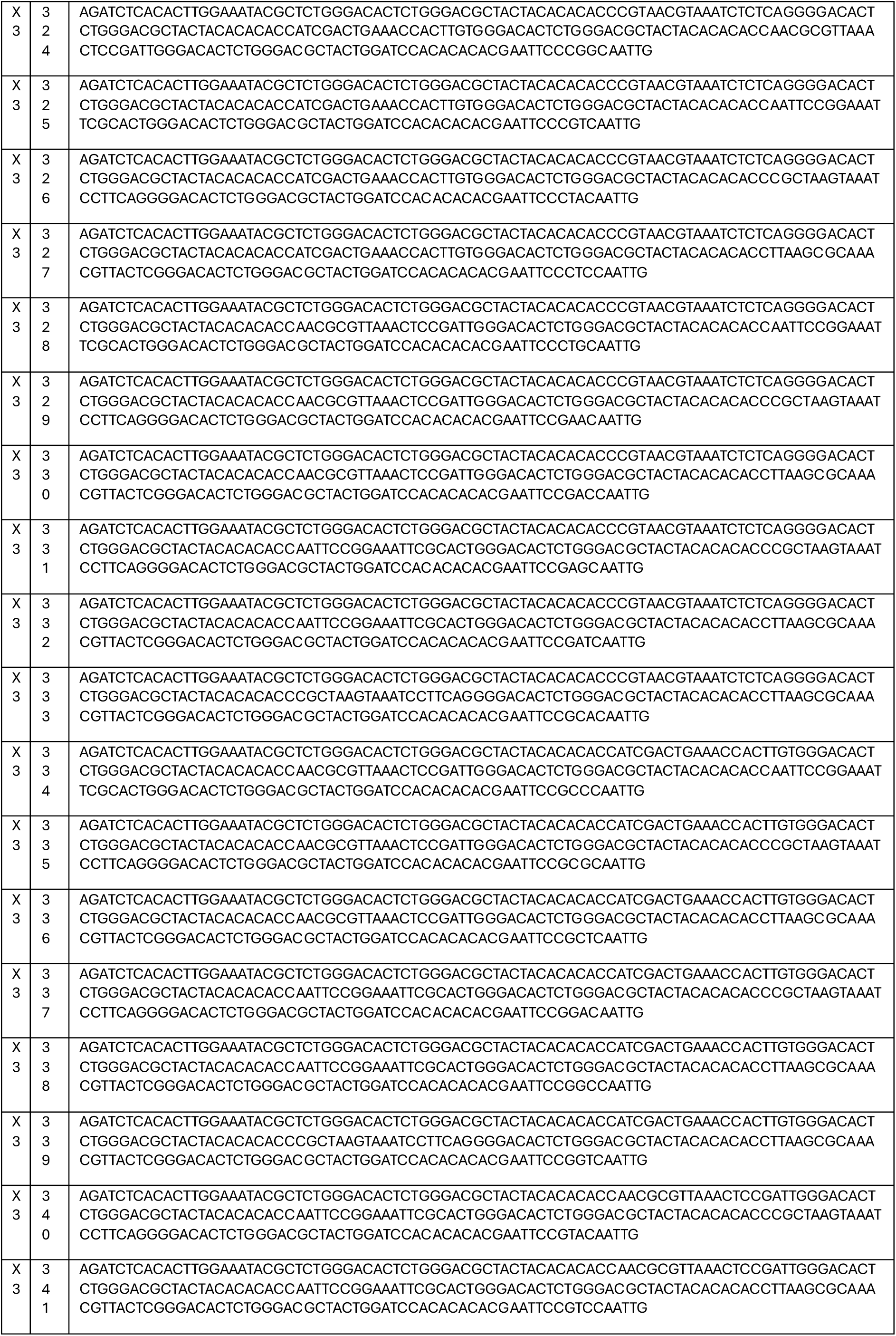
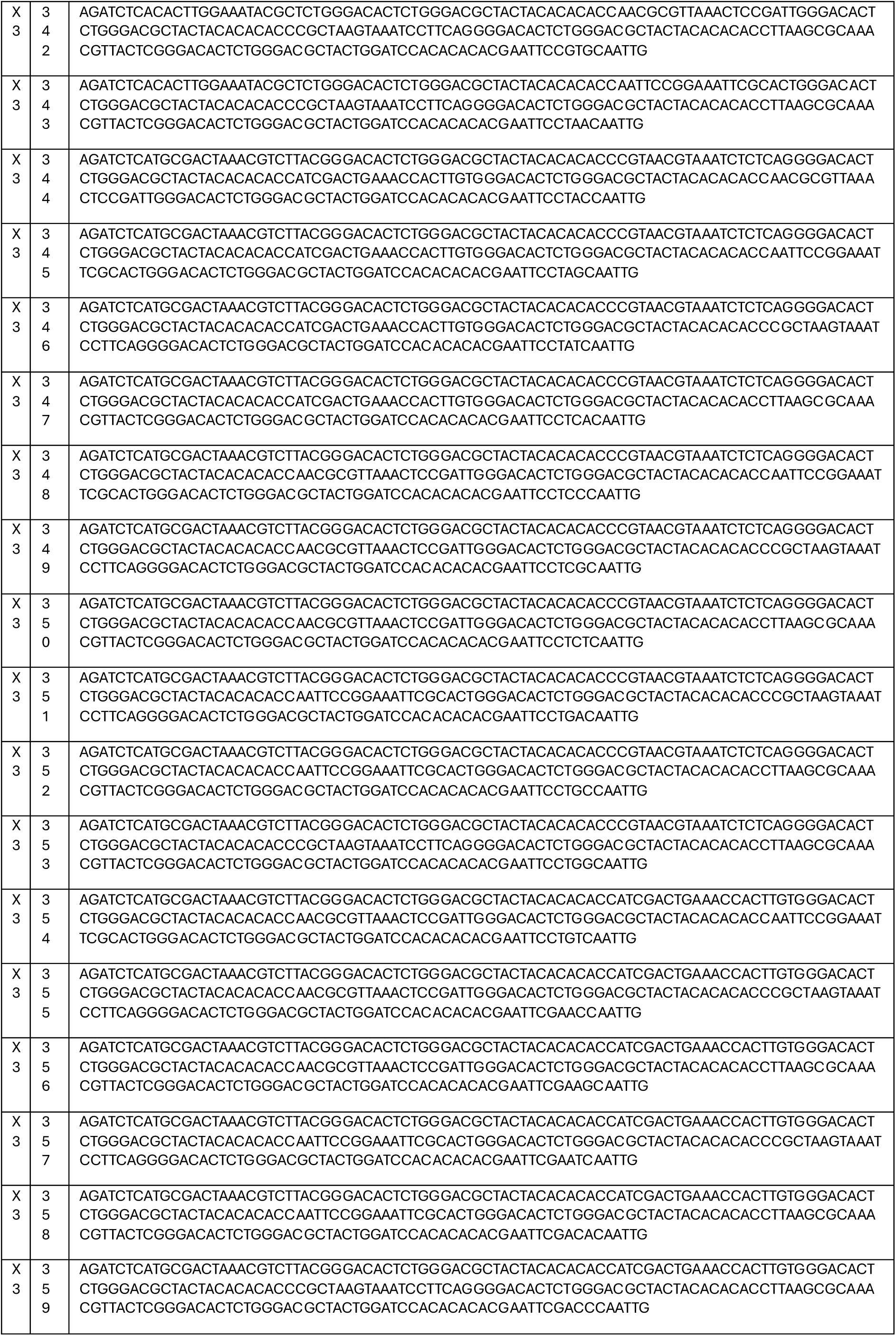
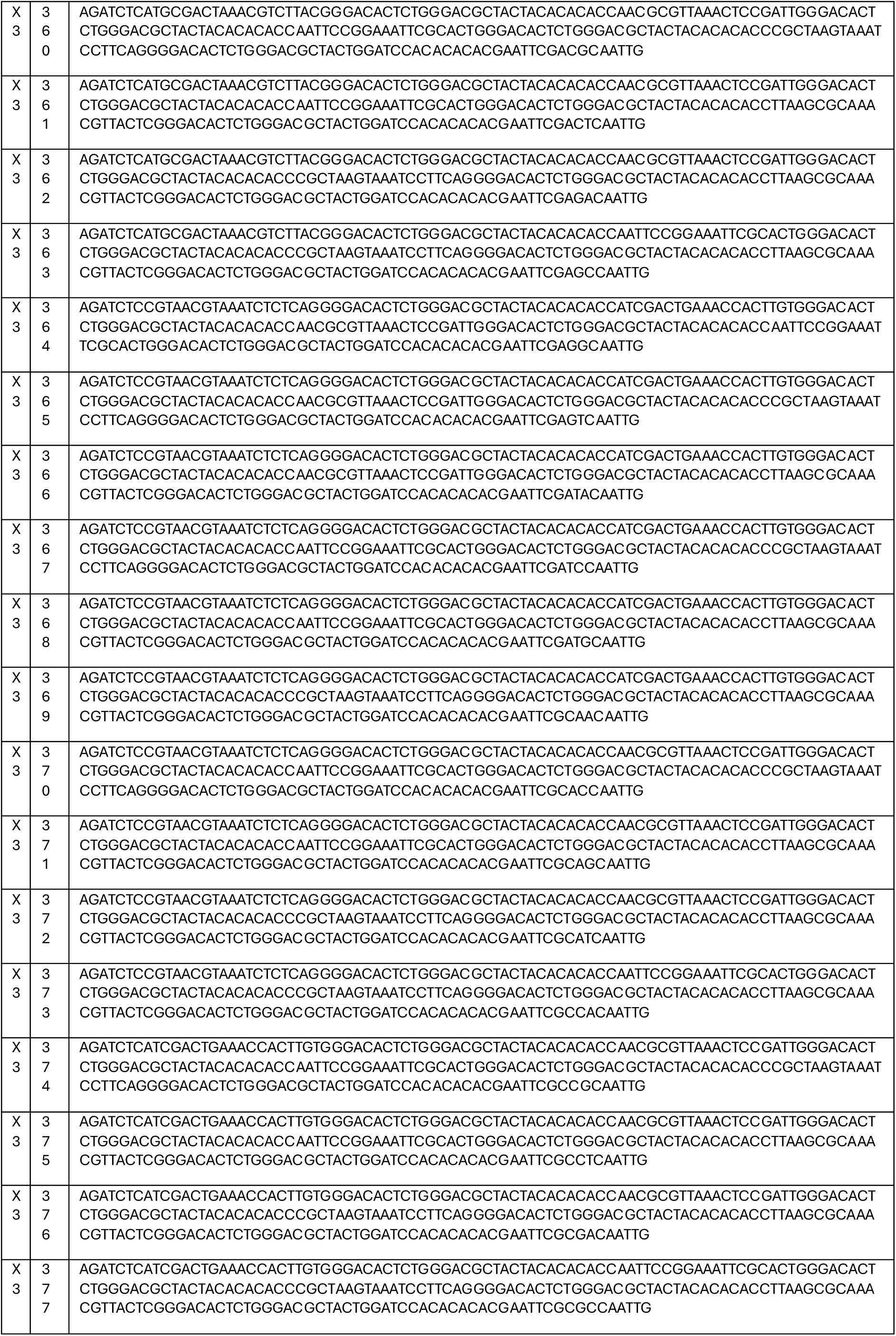
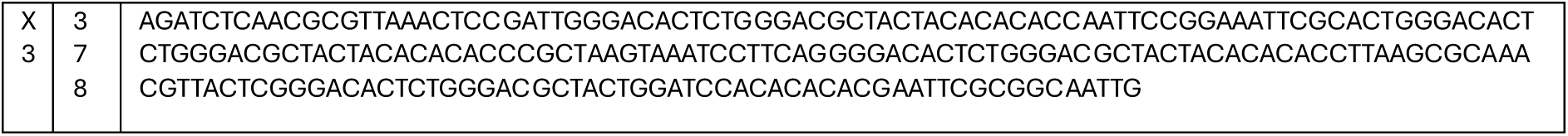
– Ordered_oligo_378.

#### X-CODE libraries assembly

A pooled ligation-based strategy was employed for X-CODE library assembly^60^. The X1 oligo mix was digested SalI-HF and MluI, while X2 and X3 were digested with MfeI (NEB; # R3589S) and BglII (NEB; #10159523). In the first ligation, gel purified X1-MluI-SalI oligos were assembled into the CROPseq-puro-filler-LTR-MluI-HF/SalI-HF backbone. The product was transformed into competent cells, amplified, and purified. The purified CROP-X1 was digested with BamHI (NEB; #R0136S) /EcoRI (NEB; #R3101S), gel purified and dephosphorylated using CIP (NEB; # M0525S), following the manufacturer’s instruction.

#### 15K library

The second ligation assembly was prepared by combining CX1BamHI/EcoRI/CIP and X2-MfeI-BglII. The ligation product was electroporated into competent cells, amplified and purified. For add-ons integration, the scaffold-gRNA segment was amplified from CROPseq-puro-v2 (Addgene; #127458) using primers (Table 4 - Add-ons Primers) containing BbsI (forward) and XbaI (reverse) sites with an ACAC linker, followed by PCR purification, digestion, and gel purification. The digested amplicons were ligated into the BbsI/XbaI–digested CROPX1X2 backbone overnight at 16 °C, purified, and electroporated into competent cells to generate CROPX1X2.1. Subsequently, the WPRE-SplitLTR-TSO-U6 fragment was PCR-amplified with primers (Table 4 - Add-ons Primers) carrying BamHI/EcoRI sites and ACAC linkers from the ClonMapper vector (Addgene; #137993), digested, and ligated into BamHI/EcoRI–digested CROPX1X2.1. The final ligation product was purified and electroporated into competent cells to generate the CROPX1X2.2 15K library. The cloning was validated through Sanger sequencing of 5 randomly picked colonies. 3 out of the 5 colonies picked for validation represented a complete set of the 18 BUs of the 15K library, and they were transduced in Tramp-C1 cells to create the F, I and H lines.

**Table 4.**
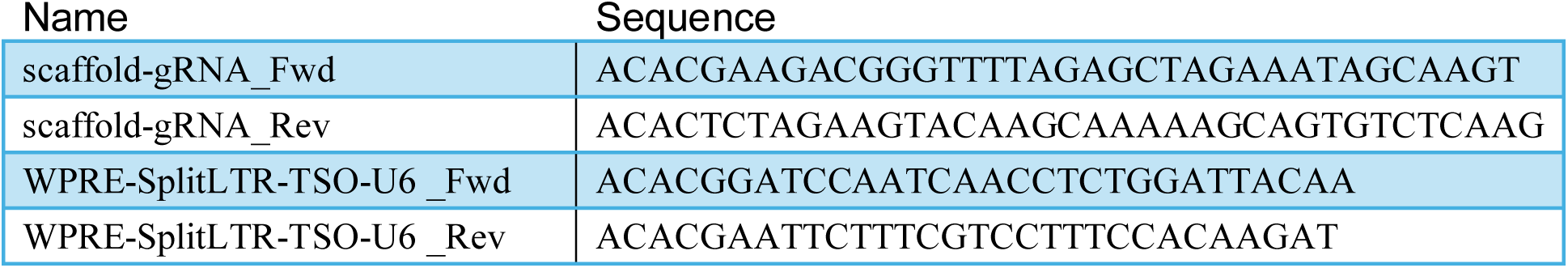
– Add-ons Primers.

#### 2M library

The third ligation assembly was prepared by combining CX1X2BamHI/EcoRI/CIP and X3-MfeI-BglII. The ligation product was electroporated into competent cells, amplified and purified. For add-ons integration, the same strategy used for the 15K library was implemented. The cloning was validated through Sanger sequencing of randomly picked colonies. 4 colonies picked for validation represented a complete set of the 27 BUs of the 2M library, and they were transduced in RM-1 cells to create the W, Z, Y and J lines. Sanger sequencing also revealed a high level of recombination that was not compatible with library scale-up. Accordingly, a different strategy was adopted for the assembly of the 2M library. The ClonMapper vector (Addgene; #137993) was mutated through mutational PCR and Gibson assembly (Table 1 – Gibson Assembly) to remove the BglII site from within the WPRE-SplitLTR-TSO-U6 sequence. The mutated WPRE-SplitLTR-TSO-U6 fragment was PCR-amplified with primers (Table 4 - Add-ons Primers) carrying BamHI/EcoRI sites and ACAC linkers, digested, and ligated into BamHI/EcoRI–digested X3 minigenes. The X3-mutWPRE-SplitLTR-TSO-U6 was then extracted from the minigenes using MfeI and BglII. The third ligation assembly was prepared by combining CX1X2.1 BamHI/EcoRI/CIP and X3-mutWPRE-SplitLTR-TSO-U6 - MfeI-BglII. The ligation product was PCR purified and used directly for lentiviral production. All the intermediate clonings were validated through Sanger sequencing.

#### Lentiviral production and transduction

HEK293T cells were seeded into 10 cm plates at a density of 20%. The following day, cells were transfected with the expression plasmid, VSVg, GAG and REV packaging plasmids, using Lipofectamine 3000 (Thermo Fisher; #L3000015) following the manufacturer’s instructions. Media was replaced after 16 h and collected after 48 h, filtered through a 0.45 μm filter, and concentrated by ultracentrifugation at 24,000 rpm for 2.5 h at 4 °C. The resulting pellet was resuspended in HBSS (Thermo Fisher; #14170112) and stored at –80 °C. For infection, 20,000 cells were seeded in 10 cm dishes and exposed to varying concentrations of lentiviral particles in the presence of 5 μg/mL Polybrene (Fisher Scientific; #12965169). After 48 hours, cells were selected with 1 μg/mL Puromycin (Thermo Fisher; #A1113803). Once the selection was completed, we calculated the MOI of the infections. An MOI of <0.2 was maintained, except in specified cases where MOI <0.3 was used.

#### Clonal retrieval

Clonal retrieval was performed following the ClonMapper protocol^31^. Briefly: barcode arrays were generated using three pairs of overlapping oligonucleotides encoding the desired barcode sequence flanked by overlapping overhangs. Complementary oligos were annealed by heating to 80 °C and cooling to room temperature, producing three dsDNA blocks. These blocks were phosphorylated and ligated sequentially to assemble the final barcode array. The ligation product (∼170 bp) was selected by size via 2% agarose gel electrophoresis and gel purified. The purified array was inserted into the Recall-miniCMV-sfGFP plasmid backbone (Addgene; #137995), previously digested with BbsI (NEB; #R3539L), using T4 ligase at a 10:1 insert:vector ratio at 37°C for 1h. The assembly was transformed into chemically competent cells. Correct insertion of the barcode array was confirmed through Sanger sequencing. For functional recall, cells were transfected with the assembled recall plasmid and dCas9-VPR plasmid (Addgene; #63798) using Lipofectamine 3000 following the manufacturer’s instructions. After 16–18 h, medium was replaced and after 48h post-transfection, cells were harvested, washed, and filtered through a 35 µm strainer. GFP positive cells were gated as recalled barcoded lineages and sorted for downstream expansion or analysis.

### X-CODE assay for CyTOF

RNA detection by CyTOF was performed following an adapted version of the PLAYR protocol^43^. 1.0–1.5 × 10⁶ cells were counted and collected per sample; this range is optimised for TRAMP-C1 and RM-1 cells, and may require adjustment for other cell lines. Cells were pelleted (before fixation: 500g, 3 min, 4 °C), washed once in PBS, and resuspended in 200μL of 1μM Cisplatin-194Pt (Standard BioTools; #201194) in PBS for 5 min at room temperature.

#### Preparation of metal-conjugated detection oligonucleotides

Metal conjugation to detection oligonucleotides was performed following the PLAYR protocol^43^. Briefly, maleimide-activated Maxpar metal-chelating X8 polymers (Fluidigm; #201300) were loaded with metals and purified according to the manufacturer’s instructions. Detection oligonucleotides containing a 5′ thiol modifier (Table 1 – Probe Seq) were resuspended in DEPC-treated water (250 μM) and reduced with 50 mM TCEP for 30 min at room temperature. Reduced oligonucleotides were ethanol precipitated, resuspended in C buffer, and conjugated to X8 polymer at 2 nmol oligonucleotide per reaction for 2 h at room temperature. Unconjugated oligonucleotides were reduced by addition of TCEP to a final concentration of 5 mM for 30 min. Conjugates were purified using 30-kDa centrifugal filter units, washed twice with DEPC-treated water, resuspended at 1 μM in DEPC-treated water, and stored at 4 °C.

#### Optional extracellular antibody staining

Cells were washed once in ice-cold Cell Staining Buffer (CSB; Standard BioTools; #201068), resuspended in 100 μL of antibody (Table 5 - Antibodies) cocktail in CSB and incubated on ice for 30 min with frequent agitation. Cells were then washed with CSB, followed by PBS, and resuspended in 100 μL of freshly prepared 5 mM BS3 (Thermo Fisher; #10249963) in PBS, followed by 30 min incubation on ice with regular agitation. Cells were resuspended in ice-cold 1.6% PFA in RNase-free PBS (∼1mL per 1 × 10⁶ cells) for 10 min on ice. After centrifugation, the pellet was loosened, and ice-cold methanol (100μL per 1 × 10⁶ cells) was added dropwise while vortexing to prevent clumping. Cells were incubated for 10 min on ice, then stained as described below. Alternatively, cells were stored in methanol at –80 °C for several months without significant RNA degradation.

**Table 5.** – Antibodies.

| **CyTOF** |  |  |  |
| --- | --- | --- | --- |
| **Antibodies** | **Clone** | **Metal** | **Cat#** |
| pRB | J112-906 | 150Nd | 3150013A - Standard Biotools |
| Ki67 | B56 | 209Bi | Custom conjugation - Standard Biotools |
| pERK1/2<br>[Thr202/Tyr204] | D1314.4E | 167Er | 3167005A - Standard Biotools |
| p4EBP1<br>[Thr37/46] | 236B4 | 149Sm | 3149005A - Standard Biotools |
| CD24 | M1/69 | 160Gd | 101802 - BioLegends |
| CD49f | GoH3 | 141Pr | 313602 - BioLegends |
| **PhenoCycler**<br>**(Akoya)** |  |  |  |
| **Antibodies** | **Bx** | **reporter** | **Cat#** |
| CK5 | #049 | Atto 550 | 5250019 - Akoya |
| CK8 | #070 | Alexa Fluor 647 | 5550024 - Akoya |
| CD45 | #007 | Alexa Fluor 488 | 4150002 - Akoya |
| Vimentin | #031 | Alexa Fluor 488 | 46173 - CST |
| AR | #052 | Atto 550 | 5250012 - Akoya |
| Ki67 | #047 | Atto 550 | 4250019 - Akoya |
| Na-ATPase | #017 | Atto 550 | ab167390 - Abcam |
| CD11b | #025 | Alexa Fluor 488 | 4150015 - Akoya |
| CD3 | #021 | Alexa Fluor 647 | 4550109 - Akoya |
| CD19 | #020 | Atto 550 | 4250014 - Akoya |
| MHC II | #014 | Atto 550 | 4250003 - Akoya |
| F4/80 | #006 | Alexa Fluor 647 | F401 - Leinco |
| Ly-6C | #037 | Alexa Fluor 488 | L400 - Leinco |
| NKP46 | #046 | Alexa Fluor 488 | N320 - Leinco |
| CD44 | #028 | Alexa Fluor 488 | C501 - Leinco |
| FOXP3 | #041 | Alexa Fluor 647 | F405 - Leinco |
| CD4 | #026 | Atto 550 | 4250016 - Akoya |
| CD11c | #030 | Alexa Fluor 647 | 4350013 - Akoya |
| CD8a | #029 | Atto 550 | 4250017 - Akoya |
| Ly6g | #024 | Alexa Fluor 647 | 4350015 - Akoya |
| CD31 | #002 | Atto 550 | ab282746 - Abcam |
| CC3 | #001 | Alexa Fluor 488 | 9661 - CST |
| SMA | #019 | Alexa Fluor 488 | ab242395 - Abcam |
| CD206 | #003 | Alexa Fluor 647 | ab64693 - Abcam |
| Synaptophysin | #034 | Alexa Fluor 488 | ab14692 - Abcam |

Upon removal from –80 °C storage, cells were thawed for 5 min on ice then pelleted (after permeabilisation: 800g, 3 min, 4 °C), washed once with 500 μL PBSTR+ (0.1% tween20 (Sigma-Aldrich; #P2287), 40 U/ml RNAsin Plus (Promega; #N2615) in PBS) and immediately resuspended in 500 μL of hybridisation buffer (1% Tween20, 2.5% PVS (Sigma-Aldrich; #278424), 1x SSC (Fisher Scientific; #10418353), 100 μg/ml salmon sperm (Thermo Fisher; #15632011), 20 mM RVC (NEB; # S1402S), 40 U/ml RNAsin, and a hybridisation probe mix: 100 nM X-CODE probes (Table 6 – Probe Seq), 1 μM universal probe (Table 6 – Probe Seq), 100 nM PuromycinR and β-actin probes (Table 6 – Probe Seq) in RNase-free water (probe mix was incubated at 90 °C for 5 min, then cooled on ice before use)).Cells in Hybridisation buffer were incubated at 40 °C for 2 h in a thermal mixer with agitation at 1,400 rpm. Cells were then washed twice with 500 μL ice-cold PBSTR (0.1% Tween20, 4 U/ml RNAsin in PBS) and incubated in 500 μL of post-hybridisation buffer (4x SSC, 0.1% Tween20, 4U/ml RNAsin in PBS) at 40 °C for 20 min. Cells were washed twice with PBSTR and resuspended in 500 μL of Ligase buffer (1X ligase T4 buffer, 4 U/ml RNAsin, 5 U/ml T4 ligase (Thermo Fisher; #B69) in RNase-free water). Cells were incubated at 21 °C for 1 h at 1,100 rpm. Cells were washed twice with PBSTR. Cells can be kept overnight in PBS after ligation without significant signal loss. Cells were centrifuged and resuspended in 500 μL of RCA buffer (1x EQPhi buffer, 250 μM dNTP, 1 mM DTT (Fisher Scientific; #10699530), 5 U/ml EquiPhi DNA polymerase (Thermo Fisher; #B39) in RNase-free water) and incubated at 37 °C for 5 h with agitation at 1,100 rpm. Cells were washed twice with CSB and incubated in a CSB solution containing 3 nM of each isotope-labelled detection probes (Table 6 - Probe seq) for 1 h at 37 °C with agitation at 1,100 rpm.

**Table 6.**
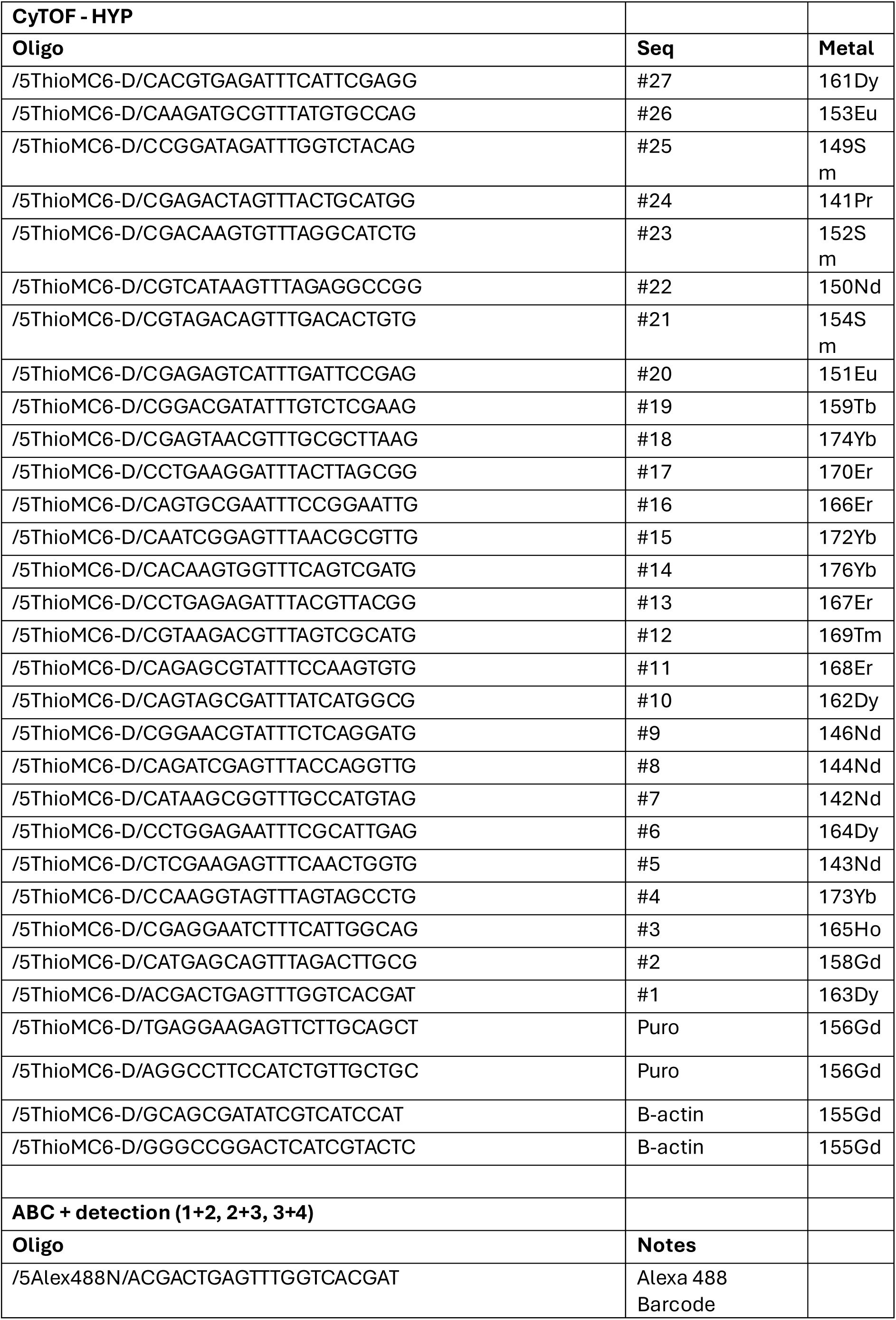
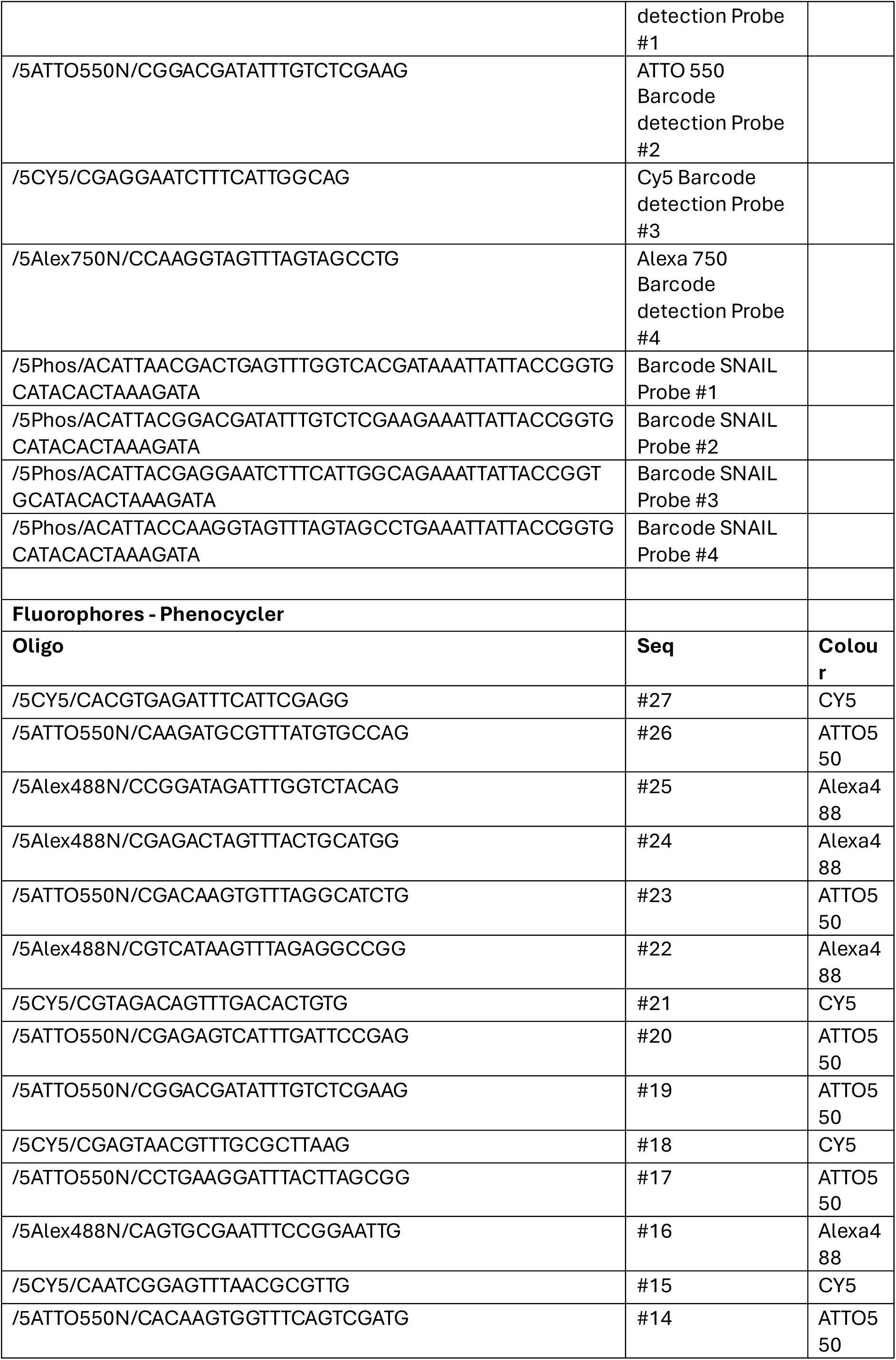
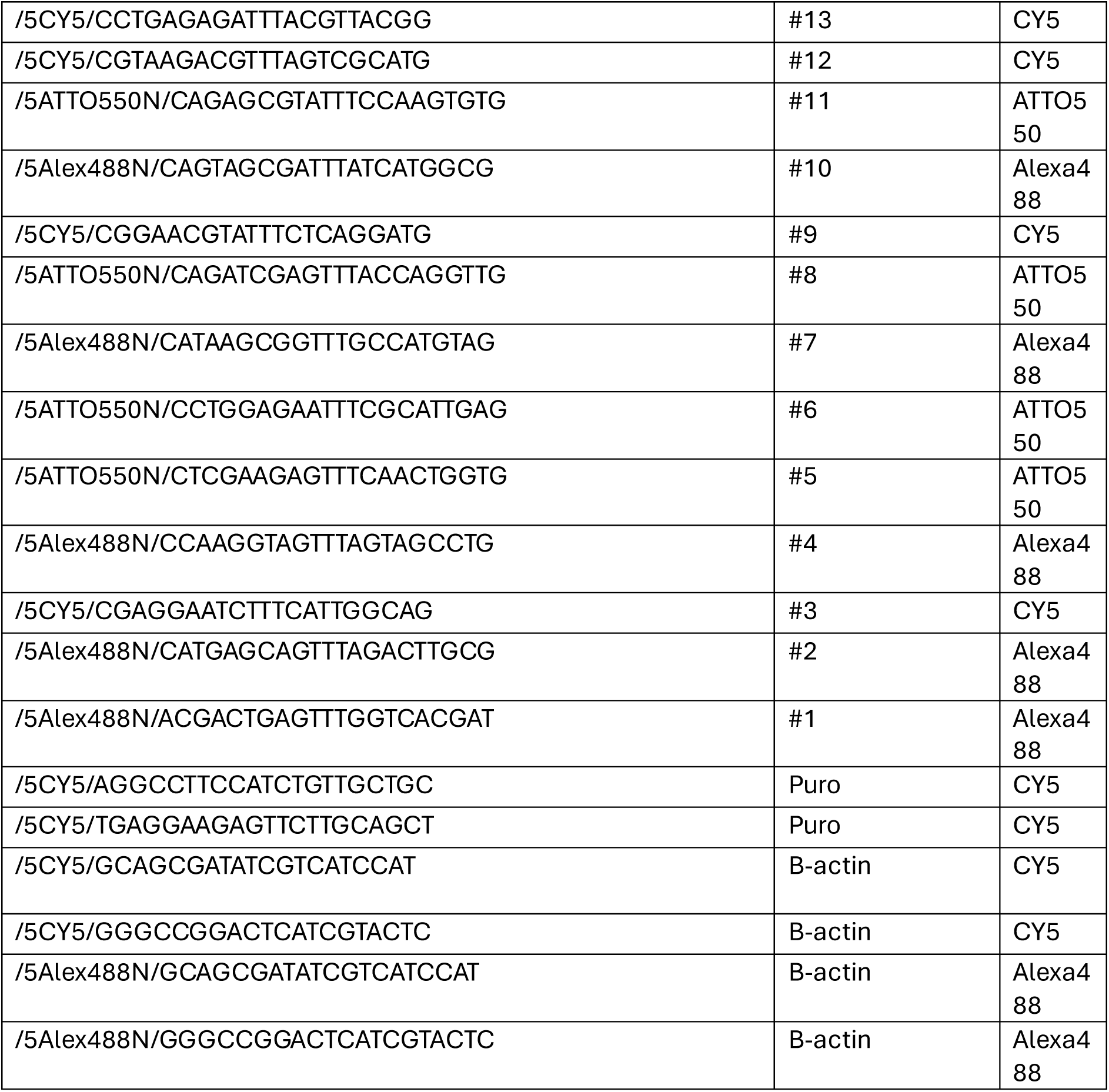
– Probe Seq.

#### Optional intracellular antibody staining

Cells were washed twice in CSB then resuspended in 100 μL intracellular antibody (Table 5 - Antibodies) cocktail on ice for 1 h with frequent agitation. Cells were washed twice with CSB and incubated in 500 μL of 62.5 nM Ir 191-193 (Standard BioTools; #201192B) in 1.6% PFA, diluted with PBS, for 1 h at room temperature, or for up to three days at 4 °C with minimal signal loss.

For CyTOF sample acquisition, the Helios mass cytometer (Standard Biotools) was calibrated according to the manufacturer’s protocols. Cells were then washed in 2 mM EDTA in CSB, then CSB, followed by Cell Acquisition Solution (CAS; Standard Biotools; #201240). Immediately before acquisition, cellswere diluted to 0.5-1 × 10⁶ cells/mL in a 10% EQ Four Element Calibration Beads (Standard BioTools; #201078) solution with 2 mM EDTA in CAS. Cells were filtered through a 35 µL filter and acquired at 100-400 events per second.

### CyTOF data debarcoding

#### Software implementation

To facilitate the debarcoding of CyTOF X-CODE data, we developed xcode_debarcode, a Python library available at https://github.com/ICR-Data-Science/xcode_debarcode/. The library uses AnnData objects throughout to ensure interoperability with the scverse ecosystem and is organized into modules for I/O (FCS and H5AD file handling, channel mapping), preprocessing (log and arcsinh transformations, intensity filtering), debarcoding (GMM, PC-GMM, PreMessa, and manual threshold methods), postprocessing (Hamming clustering, confidence filtering, Mahalanobis distance-based cluster cohesion scoring), and interactive visualizations. The design philosophy separates preprocessing, debarcoding, and postprocessing into distinct modules, allowing users to flexibly combine different filtering strategies and confidence thresholds with the debarcoding assignments. All random operations use fixed seeds for reproducibility, and each processing step writes results to AnnData columns to facilitate inspection of intermediate states.

#### PC-GMM debarcoding

Pattern-Constrained Gaussian Mixture Model (PC-GMM) debarcoding fits a two-component Gaussian mixture model to each barcode channel independently, then enforces valid 4-of-9 pattern constraints when assigning cells to barcodes. For each channel g, the model estimates the parameters of ON (high intensity) and OFF (low intensity) distributions:

p(xg) = πOFF N(xg; μOFF, σOFF²) + πON N(xg; μON, σON²)

For each cell i and channel g, the method computes the log-likelihood of the channel being in the ON or OFF state. Within each 9-channel block, all 126 valid 4-of-9 patterns are enumerated and the total log-likelihood for each pattern is computed as the sum of per-channel log-likelihoods consistent with that pattern. The maximum likelihood valid pattern is selected for each block, and the full barcode is the concatenation across blocks. The confidence score is the product of per-block posteriors. For each block, the posterior is computed as the softmax of block pattern log-likelihoods:

P(pattern | x) = exp(LLpattern) / Σp’ exp(LLp’)

This score represents the probability of the selected pattern given the observed intensities and enables ranking cells for coverage-accuracy trade-offs.

#### PreMessa debarcoding

PreMessa is a debarcoding method based on iterative top-4 selection with per-channel normalisation, adapted from the original algorithm2 to apply to the X-CODE barcode design. Within each 9-channel block, the method identifies the 4 channels with highest intensity, which by construction guarantees a valid 4-of-9 pattern without requiring bimodal distributions. The separation score is computed as the difference between the 4th and 5th highest intensities. The method then iteratively normalises each channel by the 95th percentile of its ON population and repeats the selection on normalised data. The confidence score is the minimum separation delta across blocks.

#### Hamming clustering correction

Hamming clustering is a postprocessing step that reassigns cells from low-count patterns to nearby high-count patterns based on Hamming distance. The algorithms are adapted from Starcode^61^, a clustering tool for DNA sequence barcodes, and tailored to the X-CODE 4-of-9 block structure. Low-count patterns near high-count centres are likely to be noisy children arising from measurement error rather than true barcodes. When the experimental sub-library is sparse relative to the full valid pattern space, true barcodes dominate their Hamming neighbourhoods, enabling effective error correction. Two clustering methods are available. The message passing method (default) iteratively merges small patterns into larger neighbours through chain formation: patterns are processed from smallest to largest, and each pattern that passes the count ratio test is merged into its qualifying neighbour. Chains are resolved by following each pattern to its final centre. The sphere method identifies local maxima as cluster centres and assigns non-centre patterns directly to the nearest qualifying centre without iterative chain formation. Invalid patterns (those violating the 4-of-9 constraint) are merged unconditionally into the nearest valid neighbour or centre. Key parameters include the Hamming radius (maximum distance for reassignment; default 2) and the count ratio threshold (minimum ratio of centre count to peripheral count for reassignment). When multiple centres qualify for a given pattern, ties are broken using Linear Discriminant Analysis (LDA), which computes the optimal Gaussian boundary based on the intensity distributions of the candidate centres on the channels that differ between them.

#### Simulation generative model

The simulation module generates synthetic CyTOF barcode data with known ground truth for method validation. We ran 984 simulation configurations (492 for 18-channel and 492 for 27-channel), with 10,000 cells per replicate, sub-library sizes from 1 to 3,000 barcodes, target per-channel error rates of 0.01 to 0.15, Dirichlet concentration parameters (alpha) from 0.2 to 5.0, and three replicates per configuration. The generative process proceeds as follows. First, n_barcodes valid 4-of-9 patterns are randomly selected from the full library. Cells are assigned to patterns according to a Dirichlet distribution with concentration parameter alpha. For each cell and channel, intensities are sampled from N(μON, σON) or N(μOFF, σOFF) depending on pattern state. The separation between modes is computed from the target error rate and varied by a factor of 0.7 to 1.3 across channels. An exponential transformation max(0, exp(x) - 1) produces realistic CyTOF-scale intensities.

#### PreMessa vs PC-GMM benchmark on simulations

We evaluated PC-GMM and PreMessa across the simulation parameter space. Coverage-accuracy curves were generated by ranking cells by confidence and selecting the top N%. Supplementary Figure 3F shows that PC-GMM achieves higher accuracy than PreMessa at matched coverage levels in the bimodal regime (sublibrary sizes ≥30), with an advantage of 2 to 5 percentage points at 80% coverage. Both methods detect >80% of barcodes on average even for large sub-libraries (up to 3,000), and Spearman correlation exceeds 0.75 across the tested range.

### Figure generation pipelines

#### 2M dataset

Data were loaded and channels mapped to the 27-channel configuration. Log transformation was applied. Mahalanobis distance filtering on channel sum and channel variance at the 95th percentile removed debris and doublets. PC-GMM and PreMessa debarcoding were run on log-transformed data. Sphere Hamming clustering was applied with radius=4, ratio=2, and min_count_center=500. Although the minimum Hamming distance between any two of the four expected barcodes is 10, allowing for a very aggressive radius, we used radius=4 to mimic a typical workflow where the user would estimate sub-library size via data exploration rather than having access to the exact minimum pairwise distance. Given the small expected sub-library and large valid pattern space (2M library) an aggressive ratio of 2 was appropriate.

#### 15K dataset

Data were loaded and channels mapped to the 18-channel configuration. Ground truth DNA barcodes were loaded from NGS data. Log transformation and Mahalanobis distance filtering on channel sum and channel variance at the 90th percentile were applied. PC-GMM and PreMessa debarcoding were run on log-transformed data. Patterns with count <3 were filtered out.

#### Data and Code Availability

The xcode_debarcode library, analysis code (figure generation and simulation benchmarks), and all datasets (2M, 15K, and DNA reference) are available at https://github.com/ICR-Data-Science/xcode_debarcode/.

### Sample preparations for spatial analysis

#### Chamber slide seeding

8 or 12-well (Ibidi; #80841, #81201) chamber slides with removable wells were coated with 50 µg/mL poly-D-lysine (Sigma-Aldrich; #A003E) using the manufacturer’s recommended volume per well, for 3 h to overnight at room temperature in a biological safety cabinet. Each well was washed three times with PBS, and the chamber slide was allowed to dry entirely for several hours to overnight before use. Cultured cells were counted and plated onto the chamber slides in growth media at the manufacturer’s recommended cell suspension concentration to achieve even coverage of the well area within 1–2 days.

#### OCT sample preparation

For OCT embedding, fresh-frozen samples were placed in pre-chilled OCT compound (Fisher Scientific; #23-730-571) within cryomolds and frozen over a dry-ice-chilled isopentane bath for approximately 2 min. OCT blocks were stored long-term at –80 °C in sealed containers. Before cryosectioning, artists’ brushes used to flatten sections were cleaned with an RNase decontamination solution (Thermo Fisher; #AM9780). OCT blocks, brushes, slides, and cryo-blades were acclimatised to the cryostat chamber (–20 °C) for 30 min. 10 µm sections were adhered to SuperFrost Plus microscope slides (VWR; #631 - 0448) and promptly moved to pre-chilled slide boxes on dry ice. Sectioned slides were vacuum-packed and stored at –80 °C for several weeks before X-CODE staining.

#### FFPE sample preparation

Freshly resected tissues, placed in 4% PFA, were fixed on a roller for approximately 24 h at 4 °C, then transferred to PBS. Samples were paraffin-embedded and sectioned at 5 µm according to standard histological protocols. FFPE blocks and slides were stored at 4 °C to minimise RNA degradation, and downstream X-CODE staining was performed promptly after sectioning.

#### HPMC/PVP sample preparation

HPMC/PVP embedding was performed as previously described by Dannhorn et al.^62^. Briefly, hydroxypropyl methylcellulose (HPMC) and polyvinylpyrrolidone (PVP) were weighed and dissolved in deionised water to final concentrations of 7.5% (w/v) HPMC and 2.5% (w/v) PVP. The solution was mixed thoroughly and stored overnight at 4 °C before use. For embedding, fresh-frozen samples were placed in pre-chilled HPMC/PVP within cryomolds and frozen over a dry-ice-chilled isopropanol bath for 2 min. Before transfer to a dry-ice-chilled isopentane bath for approximately 30 s. HPMC/PVP blocks were stored long-term at –80 °C in sealed containers. Cryosectioning was performed as described for OCT blocks, and vacuum-packed sectioned slides were stored at –80 °C for several weeks before X-CODE staining.

### X-CODE assay for spatial readout

#### Chamber slide preparation

Growth medium was aspirated from chamber slide wells, and cells were washed twice with PBS, then fixed with 4% PFA in PBS for 10 min at room temperature. The fixative was removed, and cells were permeabilised with ice-cold 100% methanol, dropwise, and incubated for 15 min at –80 °C before immediately commencing X-CODE staining. Alternatively, cells on chamber slides can be stored at –80 °C in sealed containers for several weeks without significant RNA degradation. To perform X-CODE staining, chamber slides were thawed on ice for ∼5 min, then the methanol was removed.

#### OCT and HPMC/PVP section preparation

Slides were thawed at 40 °C for 1 min, then immediately fixed in 4% PFA for 30 min at room temperature. Slides were washed in PBS, then submerged in 1% SDS (Promega; #V6551) in RNase-free water for 2 min, before washing twice in PBS. Slides were then incubated in pre-chilled 70% methanol on ice for 1 hour, washed twice in PBS, before immediately commencing to X-CODE staining.

#### FFPE section preparation

Slides were baked at 60 °C for 1 h, deparaffinized using xylene and rehydrated through an ethanol series (100%, 90%, 70%) in RNase-free water. To permeabilize the tissue, a freshly prepared pepsin (Sigma-Aldrich; #P7012) solution of 0.5 mg/mL in cold 0.1 M HCl was applied to cover each section entirely. Slides were incubated for 30 min at room temperature (optimisation is required for different tissue types, incubation between 5-60 min). Slides were washed twice in PBS, wicked of excess liquid, and then air-dried for 5 min before commencing X-CODE staining.

#### X-CODE spatial assay

OCT, HPMC/PVP, or FFPE-prepared slides were wiped of excess moisture, and then a hydrophobic barrier was drawn around the tissue sections using a PAP pen (Vector Laboratories; #H-4000), which was allowed to dry completely. Immediately, PBSTR+ was added inside the barrier or to the chamber slide wells for 2 min. The PBSTR+ was carefully removed from inside the barrier or wells, then Hybridisation buffer was added and incubated for 3 h at 40 °C in a humidified oven (Advanced Cell Diagnostics, HybEZ Oven). Slides were washed twice with PBSTR. Slides were then incubated with post-hybridisation buffer for 20 min at 40 °C followed by two washes with PBSTR. Slides were incubated for 2 h at room temperature in Ligase buffer, washed twice with PBSTR, and then RCA buffer was added. Slides were incubated overnight at 30 °C in a humidified oven. Following rolling-circle amplification, the protocol stages are specific to the detection technology used.

#### Detection by fluorescence microscopy

Slides were washed twice with PBST (0.1% Tween20 in PBS) for 2 min, then incubated with 200 nM X-CODE fluorescent detection probes (Table 6 – Probe Seq) in PBST for 2 h at 37 °C in a humidified oven. Following two PBST washes, chamber wells were removed if applicable, then slides were mounted using mounting medium containing DAPI (Thermo Fisher; #00-4959-52), sealed, and dried in the dark. Images were acquired by fluorescence microscopy (confocal Zeiss LSM980 instrument and Zeiss Axio Scan Z1 slide scanner).

#### Detection by Akoya PhenoCycler-Fusion

OCT and HPMC/PVP samples were funnelled into Akoya’s antibody staining workflow for fresh-frozen samples with minor adaptations to the manufacturer’s instructions, using Akoya Staining (#7000008) and Sample Kits (#7000017) for PhenoCycler-Fusion. Briefly, slides were washed twice in PBS for 2 min, chamber wells were removed if applicable, then incubated in Staining Buffer for 20–30 min at room temperature. Next, slides were incubated with an antibody cocktail (Table 5 - Antibodies) conjugated to Akoya PhenoCycler barcodes for 3 hours at room temperature in a humidity chamber. Post-staining, slides were washed with Staining Buffer twice for 2 min each, then fixed in 1.6% PFA for 10 min at room temperature. Slides were washed three times with PBS before a 5-min incubation in 100% methanol at 4 °C. Slides were washed three times in PBS, then incubated with Fixative Reagent in PBS for 20 minutes at room temperature in a humidity chamber. Following three additional PBS washes, the slide underwent PhenoCycler-Fusion flow cell assembly and was loaded into the instrument according to the manufacturer’s instructions. If applicable, PAP pen barriers were promptly removed prior to flow cell assembly using a xylene-soaked cotton swab in a fume cupboard, without touching the tissue section, and the slides were then washed a further three times in PBS. The PhenoCycler-Fusion was calibrated, and the Reporter Plate was prepared and loaded according to the manufacturer’s instructions. The Reporter Plate included the appropriate Akoya PhenoCycler fluorescent Reporters for visualisation of antibody staining. In addition, 5 μL of 10 µM X-CODE fluorescent detection probes (Table 6 – Probe Seq) were incorporated into the Reporter Plate configuration, analogous to the inclusion of Akoya Reporters. X-CODE detection probes and Akoya Reporters can be used within the same imaging cycle, provided that three distinct fluorophores are applied per cycle, consistent with the instrument’s capacity.

#### Detection by Imaging Mass Cytometry (IMC)

For IMC, samples were washed twice with Cell Staining Buffer (CSB) for 2 min. Isotope-conjugated X-CODE detection probes (Table 6 – Probe Seq) were then added and incubated at 37 °C for 60 min. Samples were subsequently washed twice with CSB for 2 min. For nuclear staining, samples were incubated with Intercalator-Ir diluted 1:400 in PBS (300–500 µl per ∼20 mm² section) for 30 min at room temperature in a humidified chamber. Sections were rinsed twice in RNase-free water for 5 min, air-dried at room temperature, and chamber wells were removed if applicable. Slides were imaged using a Hyperion imaging mass cytometer (Standard Biotools), calibrated according to the manufacturer’s protocols.

### Mass spectrometry imaging by MALDI

MSI analysis was undertaken using a time-TOF fleX (Bruker Daltonics, Bremen, DE) instrument in positive ion mode without TIMS. The MALDI matrix, 10 mg/mL 2,5 - dihydroxyacetophenone (DHAP; Fisher Scientific; #A12185.06), in 69.95% (v/v) acetonitrile (Fisher Scientific; #A955500), and 29.95% 200mM ammonium acetate (Merck; #631-61-9) in LC grade water (Fisher Scientific; #7732-18-5), 0.1% formic acid (Sigma-Aldrich; #Z0797502 21) was used. The matrix was applied using a TM HTX3 sprayer (nozzle temperature = 50 °C, gas pressure = 10 psi, flow rate = 120 μL/min, velocity = 1200 mm/min, track spacing = 2.5 mm, number of passes = 8 in a criss-cross pattern with 2 s drying time). The instrument was mass and mobility calibrated using direct infusion ESI of Agilent tune mix (Agilent Technologies; G2431). The ion source parameters were set as follows: for MALDI, MALDI offset = 50 V and shots = 150, rate = 10 kHz. For direct infusion ESI, an end plate offset of 500 V and capillary voltage of 3000 V were used, with the nebulizer set to 0.3 bar, the dry gas at 3.5 L/min, and the dry temperature at 200 °C. The instrument parameters were as follows (for a mass range of 300–1350 m/z, using a quadrupole mass filter at 300 m/z with 850 Vpp collision RF): deflection 1 delta = −70 V, funnel 1 RF = 350 Vpp, funnel 2 RF = 350 Vpp, multipole RF = 300 Vpp, isCID = 0v, ion energy = 5 V, CID = 5 V. The TOF transfer time was 85 μs, and the pre-pulse storage was 10 μs. Imaging parameters such as pixel size and x,y coordinates were set using Flex Imaging (Bruker Daltonics) software. The majority of images were acquired at a 10 μm pixel size.

#### Post-MSI sample preparation for X-CODE staining

Slides were thawed at room temperature for 2–5 min within the vacuum packaging, then removed and immediately submerged in 2% PFA in PBS for 1 h at room temperature. Following fixation, the slides were washed in PBS for 10 min, then incubated in pre-chilled 30% methanol on ice for 10 min, or until residual matrix material was fully dissolved. Slides were washed in PBS for 10 min, after which the X-CODE spatial assay was performed as described, beginning with application of the hydrophobic barrier.

### MALDI data analysis

All mass spectrometry imaging files were converted from .d format to .mz5 format using ProteoWizard/msConvert (version 3.0.24066-8d99481) and imported into MATLAB 2024b using in-house developed code.

#### Comparison of laser power

For each of the 4 samples analysed with different laser powers, a tissue mask was determined to exclude the background pixels, and the mean spectrum across the tissue pixels was determined across the full m/z range at 0.001 intervals. The 4 mean spectra were rescaled and averaged and peak picking was performed which resulted in a total of 1613 peaks. Principal components analysis was performed on total intensity normalised (multiplied by 1000) and log10 transformed data with an added offset of 1. The non-zero pixel count for each of the 1613 variables was determined as a proportion of the total number of tissue pixels. Variables were sorted into centiles according to their mean intensity, and the 25-50-75th percentiles of peak frequency were determined.

#### MyC-CaP sample analysis

A tissue mask was determined that separated each of the 12 sections from the background, and the mean spectrum was determined across all pixels for each section. Peak picking resulted in a total of 2296 features detected across at least two of the 12 tissue sections. To account for pixel sparsity across the acquisition, image smoothing was performed using a 15 x 15 pixel Gaussian filter (sigma = 3). The dataset was normalised and transformed as described above prior to multivariate analysis using principal components analysis (PCA) and uniform manifold approximation and projection (UMAP). For univariate analysis to determine variable specificity, the data was normalised but not log10 transformed.

#### Co-registration with the fluorescence image

Clone information was exported from QuPath into geoJSON format, along with a down-sampled RGB fluorescence image. An image for each clone was generated from the geoJSON coordinates at the same size as the fluorescence image (12672 x 5184 pixels). The UMAP image of components 1-3 for each suitable section was generated and co-registered to its matching fluorescence image. Characteristic regions across each pair of images were identified and used to define affine transformations that maximised the overlap of these regions. The clone image for each of these sections was similarly warped with final image dimensions of 4407 x 2520 matching the dimensions of the mass spectrometry imaging dataset. The final outputs are a fluorescence image and clone images that map directly to the mass spectrometry image. Any tissue sections containing clones with fewer than 20 pixels were discounted. The smallest group was 25 pixels for clone 8466 in sample 303; the largest was 14339 pixels for clone 8958 in sample 285.

#### Variable specificity

Each of the 2296 variables was analysed to estimate its specificity relative to a particular clone across all sections, and to a particular clone in a single section. Only pixels with a clone annotation following co-registration were analysed. Due to the imbalance in pixel counts across clones and samples, a repeated under-sampling approach was employed. A subsample of observations was randomly selected, with which the ROC curve was generated for the group with the highest mean rank (from Kruskal Wallis test) versus all other groups. From this, the AUC and optimal FPR and TPR rates were determined. This procedure was repeated 100 times, and the sample size (per group) for each of the 6 clones was 1000 observations, and for each of the 21 unique clone/sample combinations was 20 observations. The mean and standard deviations across the 100 iterations was determined. Variables were ranked according to the mean true positive rate but discounted if there was no agreement between the clone across both comparisons, i.e. the group with the highest mean rank for the clone-only analysis must match that from the clone/sample analysis. Tentative annotation of variables was performed using the bulk structure search function of LIPID MAPS structural database (LMSD) with a tolerance of ±0.05 for all [M-H]^-^ ions (https://lipidmaps.org/resources/tools/bulk-structure-search/create?database=LMSD).

### X-CODE detection by Xenium spatial transcriptomics

#### Xenium sample preparation

Cells were attached to Xenium slides using the STAMPing protocol as previously described^52^, with minor method and equipment adaptations. Briefly, Xenium slides were assembled in supplied cassettes (10x Genomics; #1000460), and coated with 50 µg/mL poly-D-lysine (Sigma-Aldrich; #A003E) in PBS for 1 h to overnight at 37 °C. Slides were washed three times with RNase-free water, allowed to dry, before silicone barriers pre-cut to the Xenium capture area were adhered to the slide. Cells were fixed with 4% PFA in PBS (1mL per 1 × 10⁶ cells) for 30 min at room temperature, then washed twice with RNase - free water (800g, 3 min, 4 °C), and resuspended in RNase-free water containing 0.01% Triton-X-100 (Sigma-Aldrich; #T8787). The cell number and resuspension volume were estimated using ChatGPT 4o to cover the area inside the barrier adequately. The cell suspension was applied evenly to the slide, and slides were dried sequentially at 4 °C (30 min), room temperature (5 min), and on a 42 °C hot plate (1.5 h). Finally, the adhered cells were permeabilised and stored in 100% methanol at –80 °C overnight within the cassettes. Alternatively, OCT blocks were sectioned onto Xenium slides according to the manufacturer’s protocols (10x Genomics; CG000579).

#### Xenium assay

The Genomics Facility (LMS MRC) thawed the STAMPed-slide cassette assembly at room temperature for 5 minutes, removed the methanol, then washed the cells three times with PBS before proceeding directly to Probe Hybridisation as per the manufacturer’s protocols (10x Genomics; CG000582). OCT tissue slides were processed accordingly (10x Genomics, CG000579, followed by hybridisation, ligation, and rolling circle amplification (10x Genomics; CG000582). The predesigned 379-gene Xenium Mouse Tissue Atlassing Panel and the custom panel for detection of X-CODE BUs were applied. Imaging, signal decoding and analysis were performed by the Xenium Analyzer instrument (10x Genomics).

### Next-generation sequencing of barcoded samples

Genomic DNA was extracted from barcoded cells or frozen tumour tissues using the QIAGEN DNeasy Blood & Tissue Kit (QIAGEN; #69504), following the manufacturer’s instructions. For the complete library characterization, the X-CODE plasmid pool was used directly as the template for PCR amplification. Barcode sequences were amplified by PCR and subsequently gel purified. The purified amplicons were submitted for sequencing using the Amplicon-EZ service provided by Azenta, according to the company’s protocol.

#### Next generation sequencing analysis

Raw barcode sequencing data were processed using Cutadapt (https://github.com/marcelm/cutadapt; v1.18) to remove adapter sequences (5′: AGATCGGAAGAGCACACGTCTGAACTCCAGTCAC; 3′: AGATCGGAAGAGCGTCGTGTAGGGAAAGAGTGT). The trimmed R1 and R2 reads were decompressed to .fastq format and aligned using the NGmerge tool (https://github.com/jsh58/NGmerge; v0.5). Aligned reads were then compared to a predefined barcode whitelist (Table 7 – 15K_key), retaining only perfect matches. A list of the 15,876 unique barcodes with corresponding counts were generated and used for downstream barcode distribution analysis and plotting.

**Table 7.**
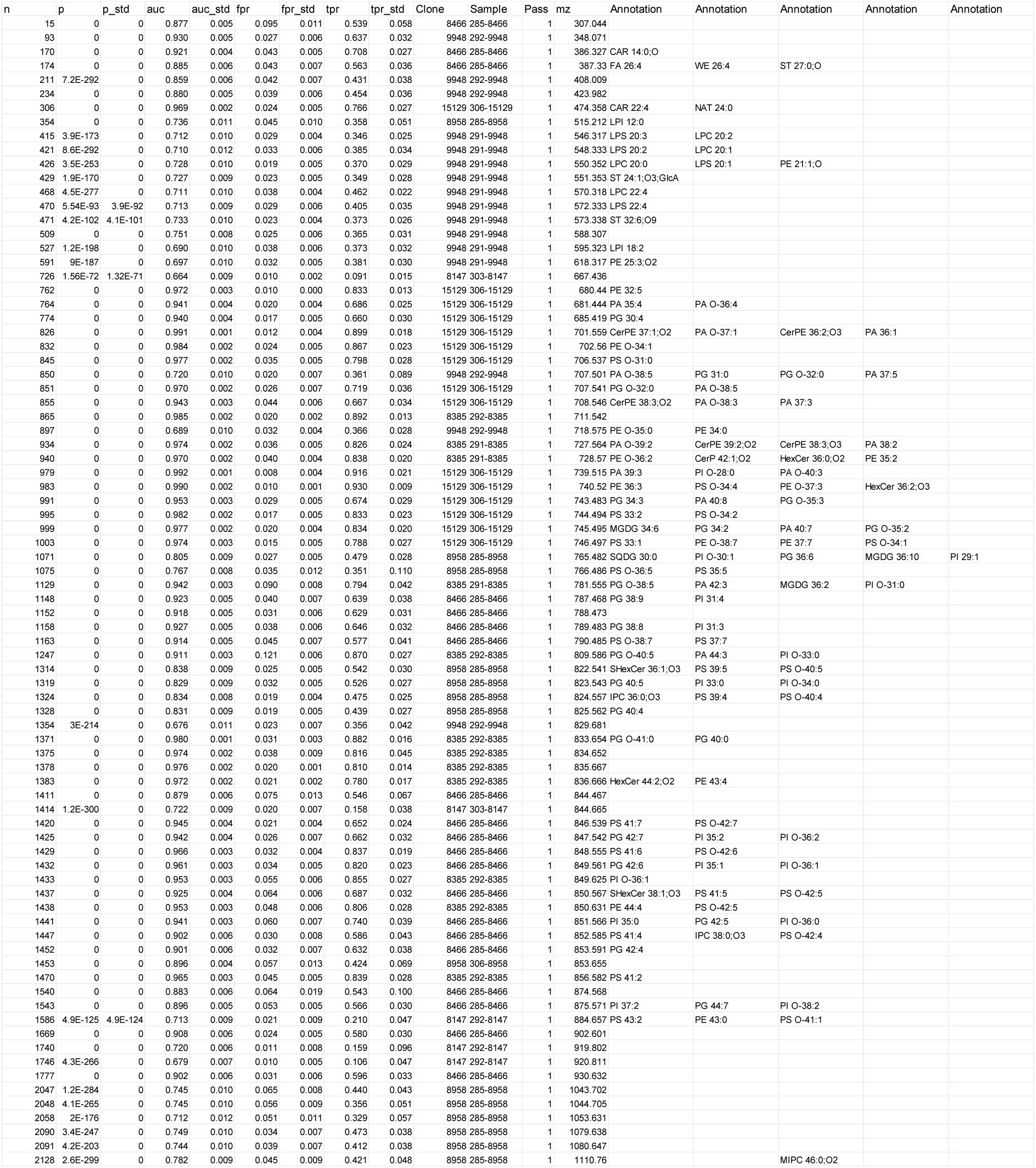
– MALDI.

### Spatial X-CODE analysis

#### Cell segmentation

To quantify X-CODE RNA amplicons (fluorescent spots) in images acquired by confocal microscopy, slide scanning, and PhenoCycler-Fusion, raw image files were visualised using QuPath 0.6.051^63^. Cellular segmentation was performed with StarDist 2D (model: dsb2018_heavy_augment; channel: DAPI) in QuPath, applying percentile normalisation (1–99%), a detection threshold of 0.5, a pixel size of 0.5 µm, and a cell expansion of 5 pixels^64^. Shape and intensity measurements were recorded for each segmented cell.

#### Spot detection

Spot detection was performed independently for each image channel using QuPath’s built-in Subcellular Detection tool. Detection parameters were optimised to ensure inclusion of visually validated true signals while minimising false-positive spot detection from background fluorescence. As standard, the minimum spot size was set to >4 pixels, and per-channel intensity thresholds were manually adjusted based on visual inspection. Additional detection parameters, including smoothing, shape- and intensity-based spot splitting, and cluster handling, were applied. QuPath’s scripting interface was used to export the total estimated number of spots and the mean intensity of each spot for each channel (representing a barcode unit (BU)) per cell. Subsequent calculations of per-cell mean fluorescence intensity for each channel were performed in Python.

#### 15K library debarcoding

Cellular BUs measurements and coordinates were exported as .csv files for downstream analysis. Decoding of barcodes was performed in Python (v3.11). To reduce the amount of any inaccurate BU detections, only the 11 most frequent BUs were considered for each cell. Each cell was then assigned a 18-digit long sequence with ones (1) and zeroes (0) corresponding to the presence of a BU. Each sequence was compared against a whitelist of 15,876 valid combinations derived from the 18 BUs in the 15K library (Table 7 – 15K_key). Cells matching a valid combination were assigned the corresponding barcode. Because of the unlikeliness that cells close together with very similar sequences/combinations would express different barcodes, a secondary classification is made. In a further classifier extension, any unassigned cell that is located within 50 µm of an assigned cell and its sequence only differs with a maximum of 2 BUs from the valid barcode, will also be assigned that same barcode. As a control, each cells’ original sequence and the top 11 adjusted sequence is saved to a .csv file together with the assigned barcode. The assigned barcodes were then imported into QuPath v0.6.0 again as a classification for each cell, allowing the cells to be visualised and categorised by the barcode.

#### Clonal phenotyping and cell population analysis

To detect single cells and quantify marker expression, manual thresholding was performed using HALO (Indica Labs). Distinct cell populations were identified based on the expression of cell-type-specific markers (Table 5 – Antibodies). Summary object data were exported from HALO across all tumours. Clone annotations were imported as layered data for downstream clone-specific cell population analysis.

### Data visualization and figure generation

Flow cytometry and mass cytometry (CyTOF) data, including t-SNE, histograms, scatter (dot / density), contour and density plots, were visualised using FCS Express 7 (v7.28.0035, De Novo Software). Bar plots and heatmaps were generated in GraphPad Prism (v10.1.0) for statistical summarisation and aesthetic customisation. UMAP projections derived from the spatial transcriptomics output (Xenium projection UMAP output file) were plotted in R using ggplot2^65^ in RStudio (v4.3.1). Alluvial plots were constructed in Plotly (v5.16.0) for Python to display cell-state transitions or lineage relationships. DNA sequencing data were visualised with SnapGene (v8.2.0) for sequence mapping and annotation, while quantitative DNAseq results and growth curves were plotted using ggplot2 in RStudio (v4.3.1). Schematic diagrams, composite figure layouts, and cartoon-style illustrations were composed using BioRender (version as of 2025). All BioRender-derived icons, templates, and assemblies were exported under a publication license and will be cited in figure legends as “Created with https://BioRender.com” in accordance with BioRender’s licensing guidelines.

---

**Supplementary Figure 1.**
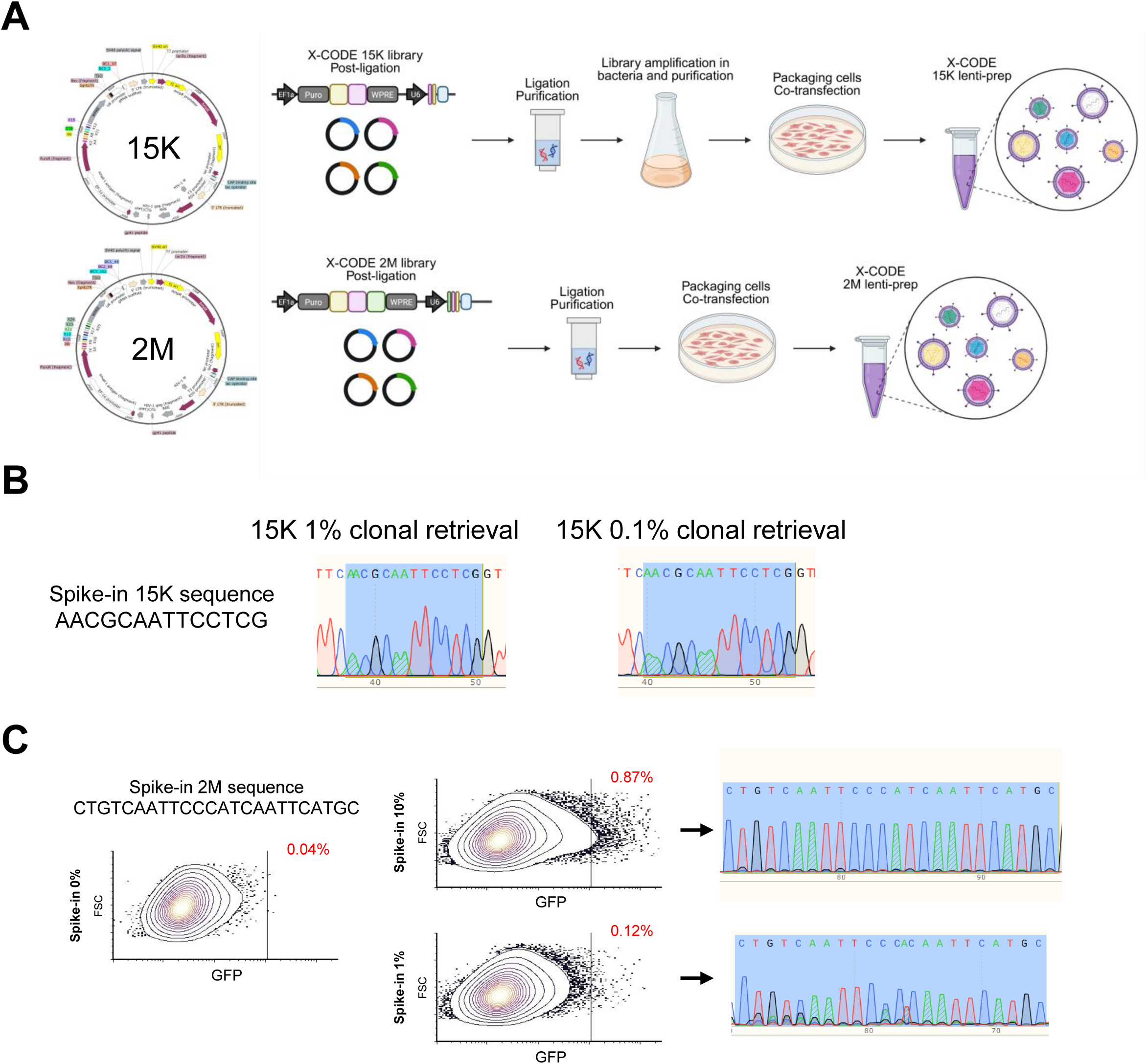
The X-CODE concept and generation. **A**) Schematic representation of the experimental workflow followed for the 15K and 2M barcode libraries (created with https://BioRender.com). **B**) Sanger sequencing chromatograms showing retrieval of the 15K-A spike-in barcode (AACGCAATT CCTCG) from sorted cells under two clonal spike-in conditions: 1% clonal retrieval (left) and 0.1% clonal retrieval (right). The recovered barcode sequence is highlighted in blue. **C**) Flow cytometry plots showing GFP expression in cells spiked with the 2M barcode (CTGTCAATTCCCATCAATTCATGC) at 0.1% (top) and 1% (bottom) frequencies. The percentage of spike-in cells detected in each condition is indicated in red. Sorted populations were subjected to Sanger sequencing, and the recovered barcode sequence is highlighted in blue in the chromatograms (right).

**Supplementary Figure 2.**
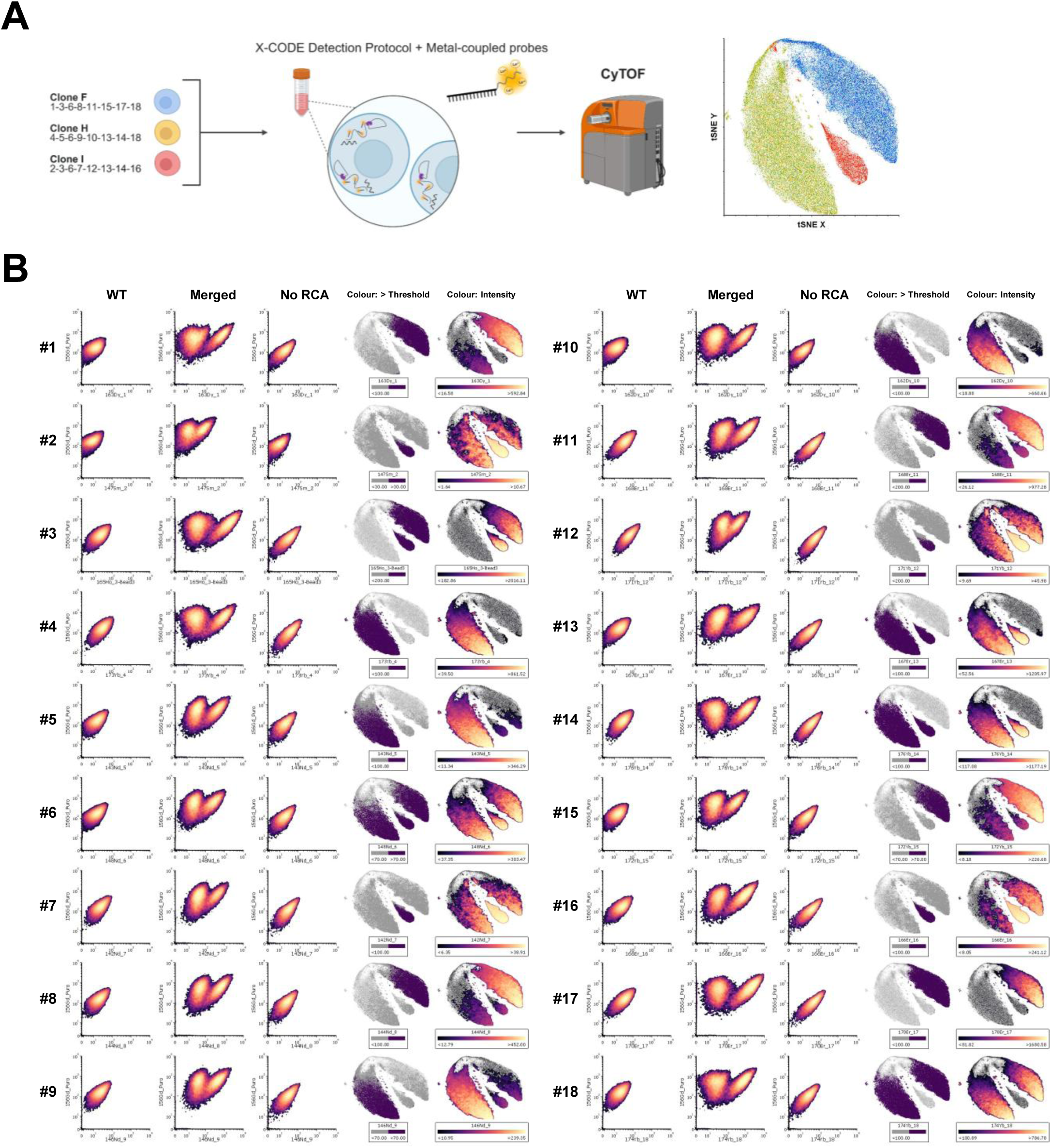
High-throughput X-CODE readout by CyTOF. **A**) Workflow illustrating validation of 15K BU detection using the X-CODE probe–based protocol adapted for CyTOF (created with https://BioRender.com). Three barcoded clones (F, H, and I), together representing the complete set of 18 BUs, were stained with detection probes, conjugated in-house to metal isotopes for mass cytometry readout. Each clone was analysed in a separate run, and the resulting datasets were merged and visualised by t-SNE, revealing three distinct clusters corresponding to the three clones. **B**) Density plots and t-SNE visualisations summarising BU-specific signal patterns for each of the 18 barcode units (BUs) in the 15K library. For every BU, panels show: (i) density plot of signal from WT cells, (ii) density plot from merged F/I/H clone data, (iii) density plot from control samples processed without the RCA step, (iv) t-SNE of merged data colour-coded for BU-positive cells above a defined signal threshold, and (v) t-SNE of merged data colour-coded by BU signal intensity.

**Supplementary Figure 3.**
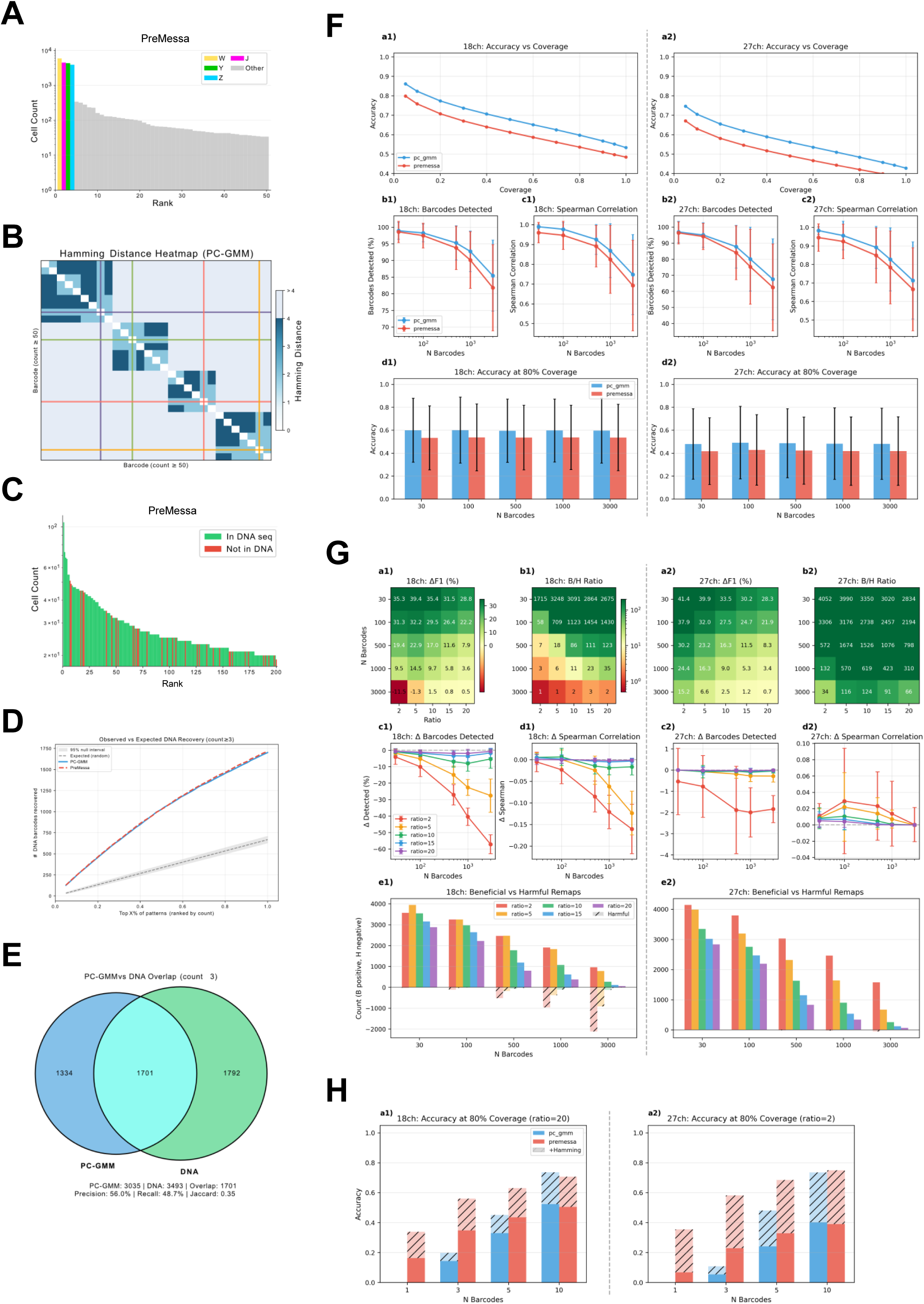
Debarcoding of X-CODE readout by CyTOF. **A**) Rank-count histogram for PreMessa debarcoding (CTV: 0.068). Expected barcodes (J, W, Y, Z) are coloured; grey bars indicate non-expected patterns. CTV = (1/2) Σi |pi - qi|, where pi is observed proportion and qi = 1/4 is expected uniform proportion. **B**) Hamming distance heatmap for patterns with ≥50 cells, hierarchically clustered. Coloured crosshairs mark expected barcodes. **C**) DNA concordance validates CyTOF debarcoding in RM-1 15K dataset. Rank-count histogram for PreMessa debarcoding with patterns coloured by DNA membership. Barcodes detected by PreMessa (count ≥ 3: 3073), 166 matching DNA barcodes in the top 200. **D**) DNA recovery as a function of coverage (top X% of patterns by count). Solid lines show observed overlap with DNA set; dashed line shows hypergeometric expectation; grey band shows 95% null confidence interval. **E**) Venn diagram showing overlap between PC-GMM detections (3,035 patterns with count ≥3) and DNA barcode set (3,493 barcodes). Overlap = 1,701. Jaccard index J = |A ∩ B| / |A ∪ B| = 0.35. **F**) Method comparison on simulated data. Set 1: 18 channels 15K library. Set 2: 27 channels 2M library. a1,a2, Coverage-accuracy curves for PC-GMM (blue) and PreMessa (red) on 18-channel and 27-channel simulations. b1,b2, Percentage of true barcodes detected as a function of sublibrary size. c1,c2, Spearman correlation between predicted and true barcode counts. d1,d2, Accuracy at 80% coverage. **G**) Message passing Hamming clustering performance. Set 1: 18 channels 15K library. Set 2: 27 channels 2M library. a1,a2, ΔF1 heatmaps showing change in F1 score after Hamming clustering for 18-channel and 27-channel configurations across sublibrary sizes (rows) and ratio parameters (columns). b1,b2, Beneficial/Harmful ratio heatmaps (log scale). c1,c2, Δ barcodes detected. d1,d2, Δ Spearman correlation. e1,e2, Beneficial and harmful remap counts. **H**) Unimodal regime comparison. Accuracy at 80% coverage for PC-GMM (blue) and PreMessa (red) with sublibrary sizes ≤10.

**Supplementary Figure 4.**
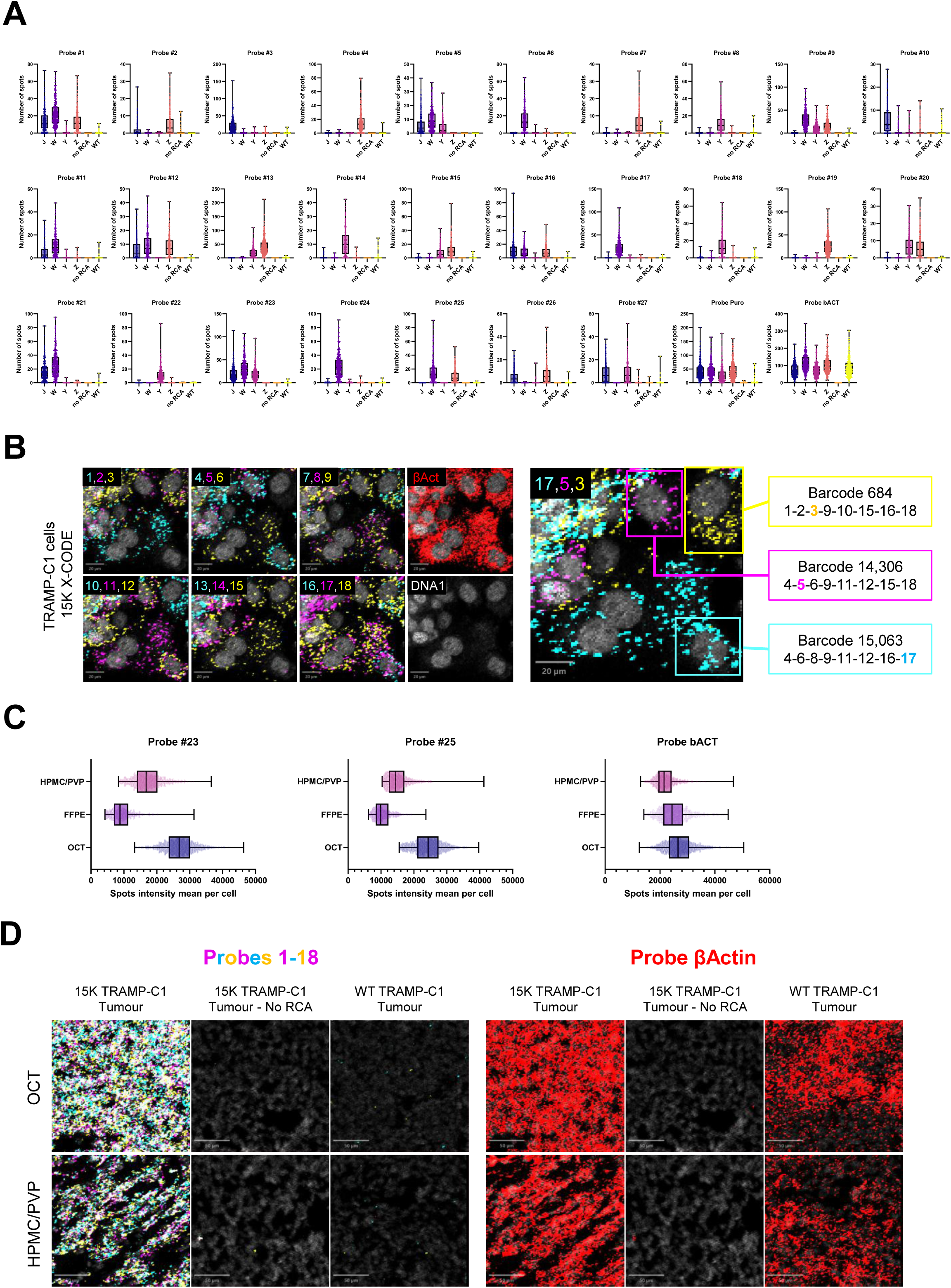
Imaging-based detection across cells, tissues and MALDI compatibility. **A**) Box plots showing the distribution of mean spot intensity per cell for all 27 probes across six experimental conditions: J, W, Y, Z, WT, and no-RCA. **B**) Representative imaging mass cytometry images of TRAMP-C1 cells expressing the X-CODE 15K library. The 18 probes used to detect the 15K library are displayed in groups of three (labelled 1–18), with individual probes shown in distinct colours. The right panel highlights three spatially localised clones within the field of view, each distinguished by a single probe (indicated), illustrating clone-specific barcode detection. Scale bars, 20 μm. **C**) Box plots comparing mean spot intensity per cell for representative probes (#23, #25, β-actin) in mixed J, W, Y, Z tumours processed using three tissue embedding methods: OCT, FFPE, and hydroxypropyl-methylcellulose/polyvinylpyrrolidone-enriched hydrogel (HPMC/PVP). **D**) Representative imaging mass cytometry images of TRAMP-C1 tumours grown in C57BL/6 mice, derived from cells expressing the X-CODE 15K library, alongside no-RCA controls and tumours generated from wild-type TRAMP-C1 cells. Samples were embedded using OCT or HPMC/PVP hydrogel. Left panels display the combined signal from the 18 X-CODE probes used to detect the 15K library, shown as grouped overlays (six probes per colour). Right panels show detection of the β-actin probe as a control. Scale bars, 50 μm.

**Supplementary Figure 5.**
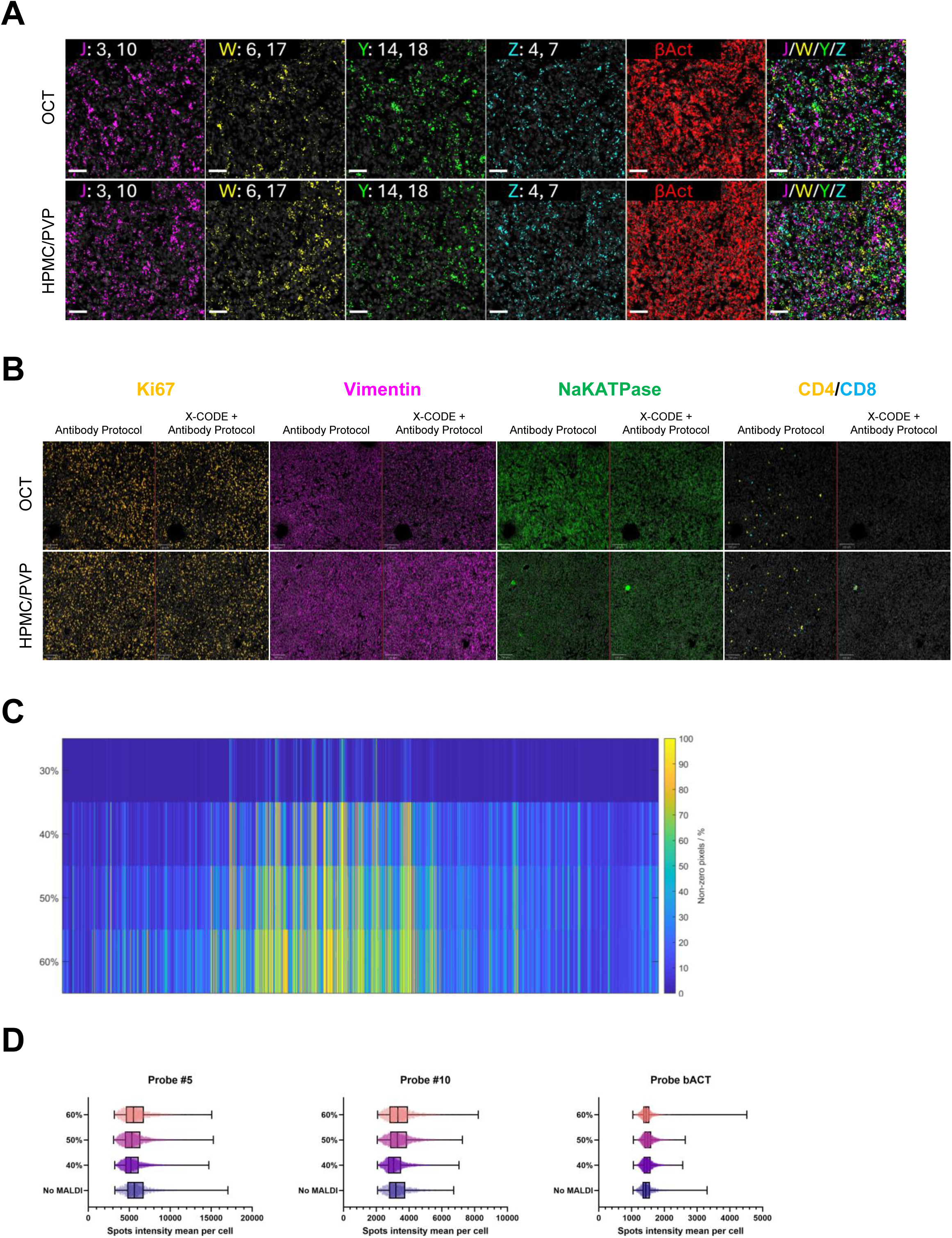
Imaging-based detection across cells, tissues and MALDI compatibility. **A**) Representative Akoya PhenoCycler images of RM-1 tumours generated by injecting an equal mixture of 2M X-CODE barcoded J, W, Y, and Z cell lines into C57BL/6 mice. Tumours were embedded using either OCT or HPMC/PVP hydrogel and processed using the X-CODE detection protocol in combination with an antibody staining workflow. For each clone, two BU probes specific to that clone are shown, illustrating clone-specific barcode detection. β-actin staining is shown as a control, and composite overlays highlight the spatial distribution of all four clones within the same tumour section. Scale bar, 50 μm. **B**) Representative Akoya PhenoCycler images of the same RM-1 tumour samples shown in the previous panel. Antibody staining is shown for samples processed using either the standard antibody protocol or an antibody staining workflow performed following the X-CODE barcode detection protocol in a consecutive tissue section. Scale bars, 100 μm. **C**) Heatmap showing feature sparsity across MALDI laser power settings (30%, 40%, 50%, 60%). Each column represents a variable, and each row corresponds to a MALDI condition. Colour coding indicates the percentage of non-zero pixels per probe, reflecting signal occupancy. **D**) Box plots showing mean spot intensity per cell for representative probes (#5, #10, β-actin) in RM-1-15K tumours processed with MALDI pre-treatment at different laser power settings (No MALDI, 40%, 50%, 60%).

**Supplementary Figure 6.**
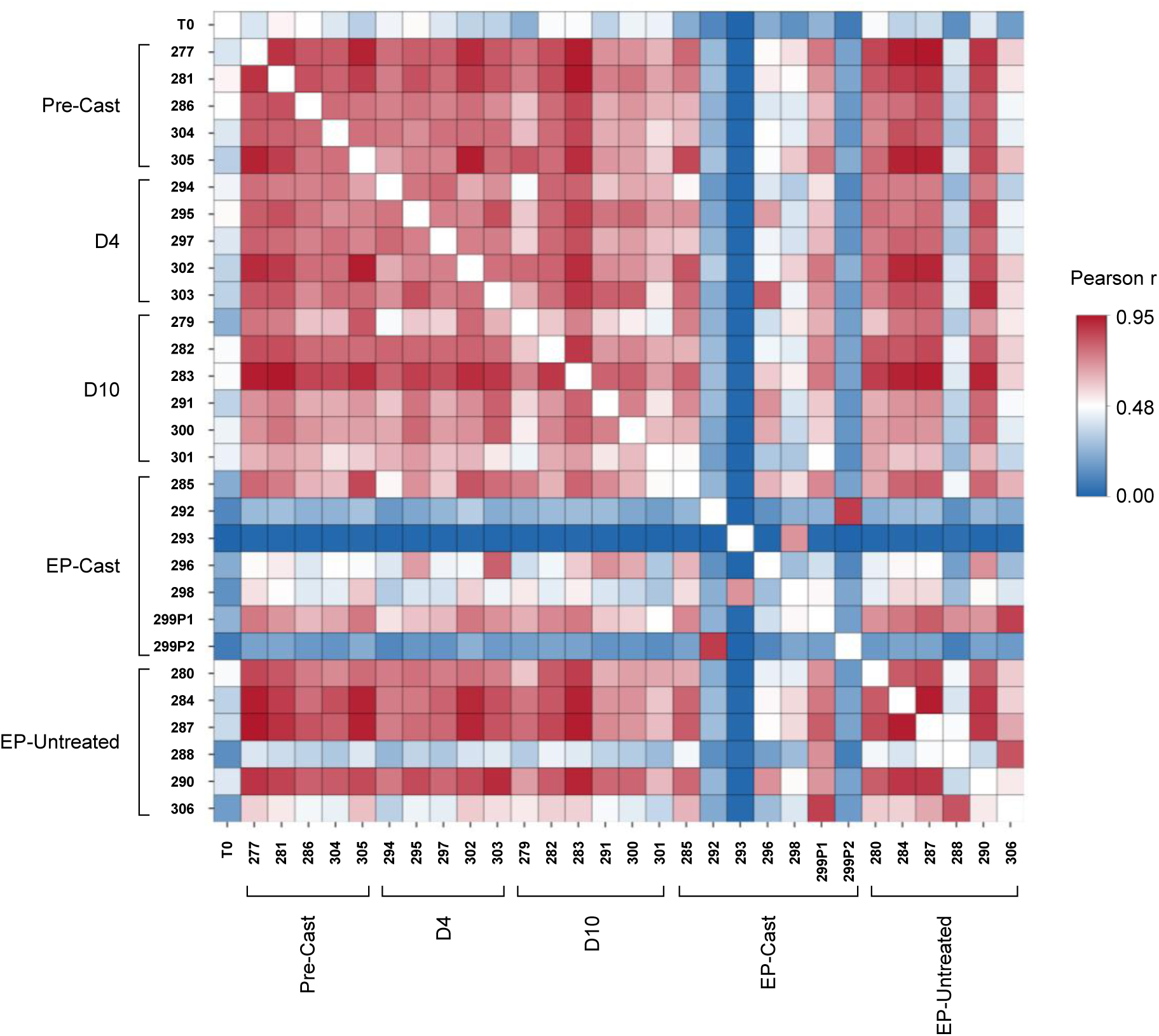
Barcode composition correlations across samples based on DNA sequencing. Heatmap of pairwise Pearson correlation coefficients across all samples. The heatmap displays Pearson correlation scores calculated between all sample pairs. Each row and column represents an individual sample, grouped by experimental condition: pre-castration (Pre-Cast), untreated endpoint (EP-Untreated), 4 days post-castration (D4 Cast), 10 days post-castration (D10 Cast), and castration endpoint (EP-Cast).

**Supplementary Figure 7.**
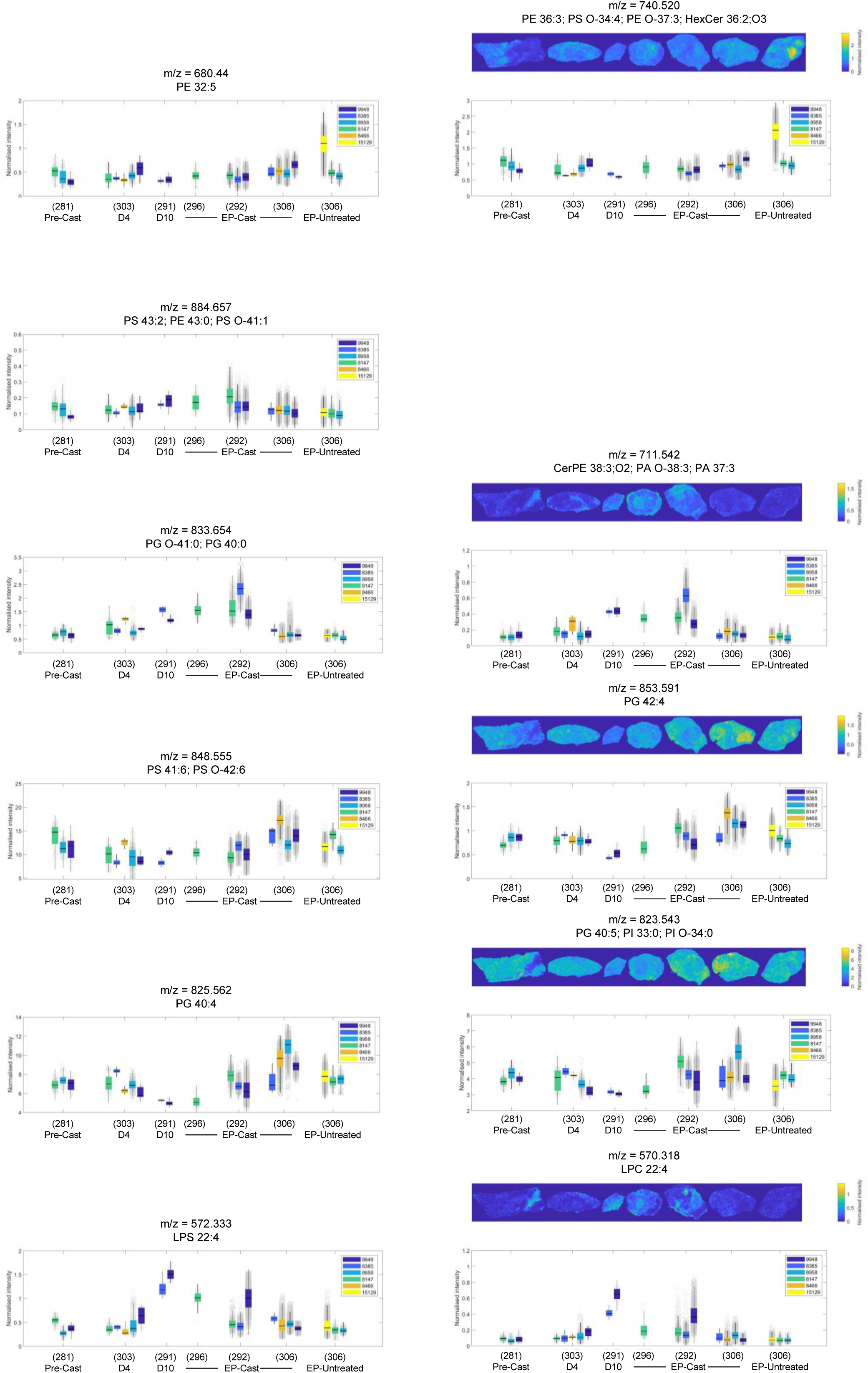
Clone-associated metabolic features identified by spatial metabolomics. Representative MALDI ion images and corresponding quantitative summaries for selected m/z features associated with dominant clones across experimental conditions. Putative annotations are indicated for each m/z value. Spatial distributions are shown for pre-castration (Pre-Cast), untreated endpoint (EP-Untreated), 4 days post-castration (D4 Cast), 10 days post-castration (D10 Cast), and castration endpoint (EP-Cast) tumours, illustrating clone-linked metabolic heterogeneity. Colour scales indicate relative ion intensity.

## Notes

### Competing Interest Statement

The authors have declared no competing interest.

